# Myelin governs hippocampal network function through the extracellular space it occupies, not only the axons it insulates

**DOI:** 10.64898/2026.11.02.763564

**Authors:** Xin Jia, Cuidie Zeng, Lingchen Liu, Tengao Gao, Jing Bi, Zhiming Zeng, Lei Liang, Chenxiang Qi, FuYuan Lin, Fangxiao Cheng

## Abstract

Myelin, the multilamellar sheath wrapping neuronal axons, enables saltatory conduction. Beyond electrical insulation, myelin occupies volume, and therefore shapes the extracellular space (ECS) through which interstitial fluid flows and neurotransmitter diffuses. How these two roles interact is unknown. We built a multi-scale model of hippocampal CA1 coupling lattice Boltzmann simulations of ECS flow to detailed networks of reconstructed neurons and glia. Demyelination enlarged the ECS volume fraction from 28 to 43 per cent up to 43 to 70 per cent across layers, lowered tortuosity from about 1.7 to 1.4, raised effective diffusivity from 0.35 to 0.50 of free solution and shortened solute breakthrough times by roughly a third, while raising flow speed yet lowering viscous dissipation. Because transmitter concentrationscales inversely with that volume, synaptic driving force fell: postsynaptic potential amplitude declined with wrap count and transmission failed entirely below four of sixteen wraps. In 200-cell networks, demyelination halved firing (pyramidal 7.9 to 4.4 Hz), rendered discharge burst-dominated, reduced band power by up to 8.7-fold and displaced the population state, which a classifier separated at 100 per cent accuracy, whereas an inflammatory glial census left every functional measure unchanged. Driving myelin continuously from intact to 12.5 per cent and back recovered the same threshold near four wraps, and firing rate and spectral power returned fully to baseline. Myelin thus governs circuit function through the space it vacates as well as the axons it insulates, with opposite consequences for clearance and for signalling, a dissociation that may matter in demyelinating disease.

## 1 Introduction

The brain’s ECS, as volume transmission pathway, constitutes approximately 20% of brain volume and plays crucial roles in maintaining homeostasis (Lei et al., 2017; Syková & Nicholson, 2008; Tønnesen et al., 2023), facilitating neurotransmitter clearance (Syková & Nicholson, 2008), and supporting glymphatic waste removal. The glymphatic system, a brain-wide clearance pathway(Iliff et al., 2012), relies on ECS flow to remove metabolic waste products including amyloid-β (Aβ) and tau proteins whose accumulation is associated with neurodegenerative diseases (Iliff et al., 2014; Ishida et al., 2022; Xie et al., 2013) and neuroinflammatory factors such as cytokines and chemokines whose dysregulation is linked to chronic stress-mediated depressive disorders (chronic neuroinflammatory conditions) (Gu et al., 2022). Recent evidence suggests that ECS architecture and flow dynamics are dynamically regulated by cellular structures(Tønnesen et al., 2023), particularly myelination (Wang et al., 2019; Wang et al., 2021); Myelinated axons comprise approximately half of the human brain volume (Stadelmann et al., 2019).

Myelin, the multilamellar sheath wrapping neuronal axons, is essential for rapid signal propagation through saltatory conduction (Nave, 2010; Stadelmann et al., 2019). In addition to its well-established role in electrical signalling, myelin shapes the geometric architecture of the ECS (Vargová & Syková, 2014). The compact myelin layers reduce ECS volume fraction and create tortuous pathways for fluid flow and molecular diffusion (Kamali-Zare & Nicholson, 2013). However, the quantitative relationship between myelination and ECS flow dynamics remains poorly understood, despite growing recognition that ECS dysfunction contributes to neurological disorders.

Computational fluid dynamics (CFD) approaches have been increasingly used to model ECS flow and molecular transport (Agudelo-Toro, 2013; Asgari et al., 2015; Bohr et al., 2022; Boster et al., 2022; Daversin-Catty et al., 2022; Guo et al., 2025; Jin et al., 2016; Kedarasetti et al., 2022; Li et al., 2025; Poulain et al., 2023; Troyetsky et al., 2021). Recent lattice Boltzmann method (LBM) studies have simulated waste clearance pathways, including the glymphatic system and blood-brain barrier, revealing their complementary roles in removing metabolic waste products such as Aβ (Yousofvand & Tithof, 2025). However, most studies have focused on simplified geometries or specific brain regions, and systematic investigations of myelination effects on ECS flow are limited. Experimental studies using two-photon microscopy and MRI have shown that ECS volume fraction and tortuosity vary with brain region and pathological conditions (Han et al., 2014; Hrabetova et al., 2018; Jiang-Xie et al., 2024), but the underlying mechanisms remain unclear.

Electrophysiological studies have extensively characterized how myelination affects action potential propagation and conduction velocity. The Hodgkin-Huxley model and its extensions, implemented in NEURON, have successfully predicted myelination-dependent changes in conduction velocity (Hodgkin & Huxley, 1952). However, the relationship between myelination, postsynaptic potential (PSP) properties, and extracellular ionic waves remains less explored. How the ECS geometry set by myelin feeds back onto synaptic transmission is a separate and largely unexamined question. Because the concentration reached by a released transmitter depends on the volume into which it disperses, an enlarged ECS should dilute it and weaken the postsynaptic response (Rusakov & Kullmann, 1998), but whether that geometric route is strong enough to alter network activity has not been tested.

Despite these advances, several critical gaps remain. First, the quantitative relationship between myelination and ECS flow dynamics has not been systematically investigated using computational approaches. Second, how myelination simultaneously affects electrophysiological changes and ECS flow dynamics remains unclear. Third, how myelination affects network-level dynamics and whether these effects scale from single-synapse to network level is unknown. Here, we address these questions by combining computational fluid dynamics (LBM) and electrophysiological modelling (NEURON) across multiple scales: (1) ECS flow and molecular transport at the tissue level, (2) single-synapse electrophysiology, and (3) network-level dynamics. We systematically compare low myelination (MY-) and high myelination (MY+) conditions and, in the dynamic models, drive myelination continuously across that range to establish the mechanistic link between myelination, electrophysiological signal properties, and ECS flow dynamics.

This paper is organised as follows. We first investigated ECS flow dynamics and molecular transport using LBM, comparing MY-and MY+ conditions. We then examined single-synapse electrophysiology, quantifying how myelination affects PSP properties and conduction velocity. Finally, we extended these findings to the network level, first contrasting two fixed myelination states in a factorial design that also varied the glial census, and then driving myelin continuously from intact through maximal loss and back, so as to locate the threshold at which network function fails and to test whether it recovers.

## 2 Materials and Methods

### 2.1 Computational Domain and Geometry

We simulated extracellular space (ECS) flow, molecular transport and electrophysiological activity in computational domains representing the hippocampal CA1 region, encompassing all four anatomical layers: stratum oriens (SO), stratum pyramidale (SP), stratum radiatum (SR) and stratum lacunosum-moleculare (SLM). Domain sizes were selected based on a representative volume element (RVE) analysis of electron-microscopic reconstructions of hippocampal neuropil at the 500-nm scale (Fig. 1a), and were then scaled to 20 × 20 μm² so as to contain a statistically representative number of cellular profiles while preserving realistic ECS geometry (Kinney et al., 2013; Mishchenko et al., 2010; Nicholson & Hrabětová, 2017). For the lattice Boltzmann method (LBM) fluid-dynamics simulations we used two-dimensional square domains of 20 × 20 μm per layer with a uniform lattice spacing of 0.01 μm (10 nm), i.e. 2000 × 2000 lattice nodes per layer. For the electrophysiological simulations the domain was extended to 20 × 20 × 500 μm so as to accommodate full axonal morphologies spanning all four layers (Bezaire et al., 2016).

**Figure 1.**
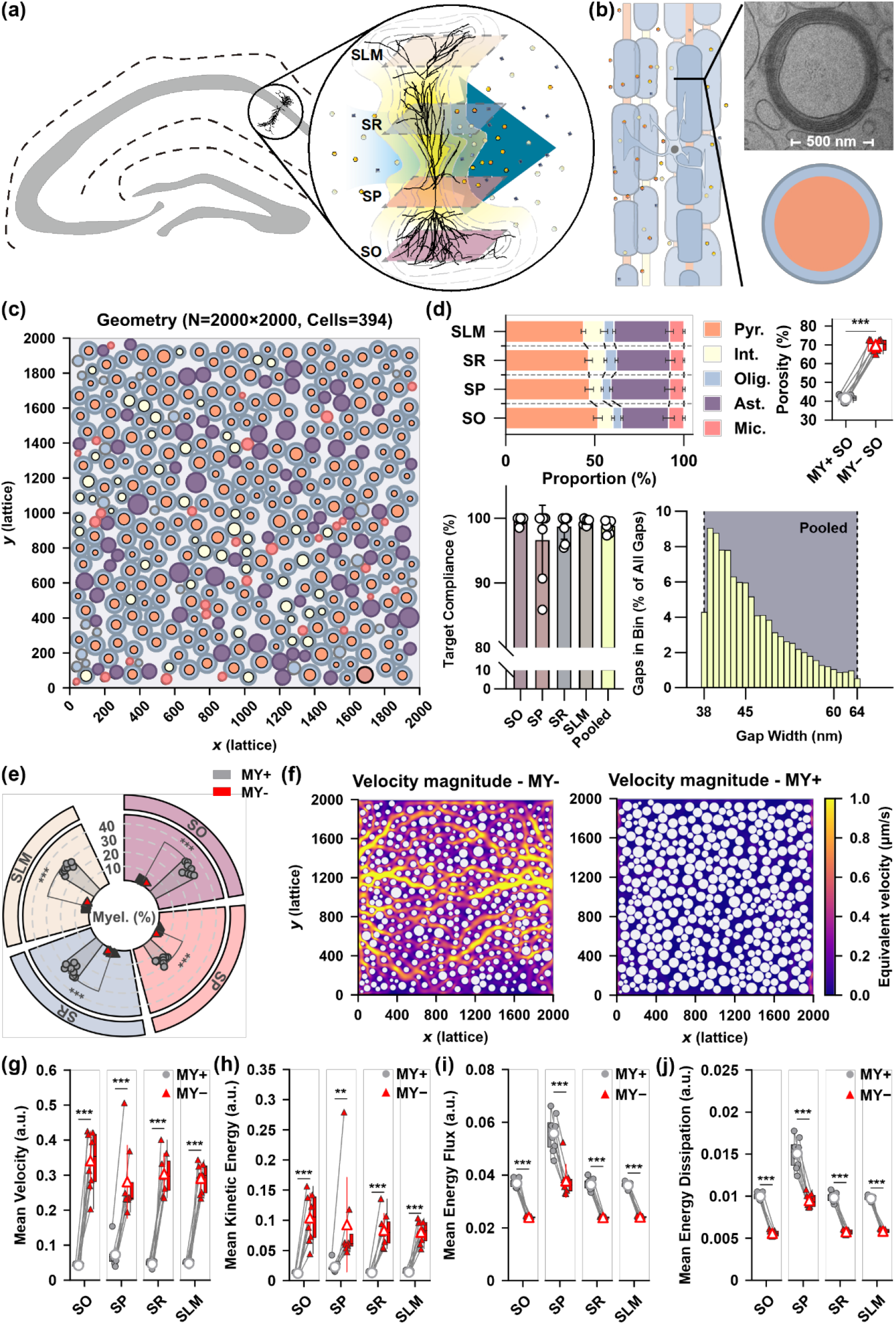
Model geometry, myelin ultrastructure, and demyelination increases interstitial fluid velocity across all CA1 sublayers. (a) Schematic of the hippocampal CA1 region illustrating the four sublayers (stratum oriens, SO; stratum pyramidale, SP; stratum radiatum, SR; stratum lacunosum-moleculare, SLM) and the reconstructed pyramidal neuron morphology used to define sublayer boundaries. Yellow traces indicate the electrophysiological (synaptic transmission) axis of the study; blue arrows indicate the fluid-dynamic (ECS transport) axis, reflecting the two complementary mechanisms examined throughout this study. (b) Ultrastructural basis of the myelin model: transmission electron micrograph of a myelinated axon (500 nm scale bar) and its abstraction into a concentric-cylinder model (orange, axon; blue, myelin sheath), with periodic internode geometry shown schematically. (c) Representative 2D computational domain for one simulation repeat (lattice size 2000 × 2000, 399 reconstructed cell processes), used as the basis for all lattice-Boltzmann flow simulations. (d) Composition and calibration of the synthetic cell-process geometry: schematic cross-sections of the modelled process types (top), target ECS volume-fraction (porosity) compliance across sublayers and pooled (bottom left; mean ± SD, \*\*\**p* < 0.001, paired comparison, the only panel in (d) subjected to formal statistical testing — the remaining descriptive panels serve as geometric calibration checks against literature targets), and the pooled distribution of cell-process gap widths (bottom right). (e) Sublayer-specific myelination ratio (%) under MY+ and MY− conditions (mean ± SD; **/\*\*\**p* < 0.01/0.001, paired *t*-test, *n* = 8 per sublayer). (f) Representative interstitial flow-velocity fields within the computational domain, illustrating streamlines (left) and velocity magnitude (right) under the demyelinated configuration. (g–j) Demyelination significantly increases mean interstitial fluid velocity (g) and mean local kinetic energy (h), while significantly decreasing mean energy flux (i) and mean viscous energy dissipation (j), consistently across all four CA1 sublayers (SO, SP, SR, SLM; grey circle, MY+; red triangle, MY−; paired comparisons, *n* = 8 independent geometries per sublayer; \*\**p* < 0.01, \*\*\**p* < 0.001).

Each layer was assigned a distinct cell-type composition reflecting the anatomical specialisation of rat CA1 (Bezaire & Soltesz, 2013). Layer-specific placement weights for pyramidal cells, interneurons, astrocytes and oligodendrocytes were: SO, 55/25/16/4 %; SP, 60/20/16/4 %; SR, 50/25/20/5 %; SLM, 45/25/25/5 %. Microglia were drawn separately as 10–20 % of the total glial quota. These layer weights act as placement probabilities and are applied under two global constraints: the neuronal fraction of all cells was drawn from the interval 0.45–0.55, and an explicit neuron-to-glia balancing constraint was enforced, so that the overall neuron-to-glia ratio was approximately 1:1 in every realisation (Keller et al., 2018). Cellular process diameters were drawn from truncated normal distributions: neuronal (axonal) processes, 0.5 ± 0.1 μm (truncated to 0.3–0.7 μm); astrocytic processes, 0.8 ± 0.15 μm (0.6–1.1 μm); oligodendrocytic processes, 0.6 ± 0.1 μm (0.4–0.8 μm); microglial processes, 0.5 ± 0.1 μm (0.3–0.7 μm). Cells were placed subject to a minimum ECS gap constraint of 38–64 nm, consistent with experimental estimates of ECS width in living brain tissue (Syková & Nicholson, 2008).

Myelination was modelled using a multi-layer approach. Myelin thickness was calculated as

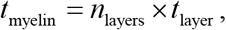

where *n*_layers_ is the number of myelin wraps and *t*_layer_ = 15 nm is the thickness contributed by one wrap, i.e. one compacted membrane bilayer pair (Sherman & Brophy, 2005). Two myelination conditions were compared: low myelination (MY−, 2 wraps for all axons) and high myelination (MY+, 16 wraps for pyramidal axons and 8 wraps for interneuron axons), the upper values corresponding to the g-ratios reported as near-optimal for rodent CNS fibres (Chomiak & Hu, 2009). In Sections 3 and 4 the wrap count was additionally varied across the full range (2–16 for pyramidal axons, 2–8 for interneuron axons) to resolve myelination gradients.

### 2.2 Computational Methods

#### 2.2.1 Lattice Boltzmann Method for Fluid Dynamics

ECS flow was simulated with the lattice Boltzmann method on a D2Q9 lattice (Chen & Doolen, 1998). The LBM recovers the incompressible Navier–Stokes equations through the evolution of particle distribution functions *f_i_* ( ***x***, *t*) :

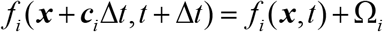

where ***c****_i_* are the nine discrete velocity vectors and and Ω*_i_* is the collision operator. Collisions were treated with the single-relaxation-time Bhatnagar–Gross–Krook (BGK) approximation (Bhatnagar et al., 1954):

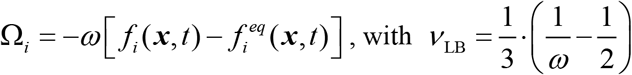

The relaxation parameter was initialised from a target creeping-flow Reynolds number of 0.01 via 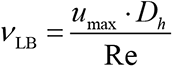 (with *u*_max_ = 0.05 lattice units and *D*_ref_ = 20 lattice nodes, twice the mean myelinated-profile radius) and 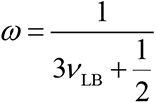, and was then constrained to the numerically stable interval [0.5, 1.95]. For the present geometries this lower constraint is active, so all production runs used *ω* = 0.5, corresponding to *ν*_LB_ = 0.5 in lattice units — the most strongly viscous, and therefore least inertial, setting available to the BGK scheme. That the resulting flow lies deep in the Stokes regime was verified a posteriori from the converged fields, where local Reynolds numbers computed with the physical kinematic viscosity of water are many orders of magnitude below unity (see Derived Flow-Field Diagnostics).

Solid cellular profiles and the upper and lower domain walls were treated as no-slip boundaries using the halfway bounce-back rule. A parabolic velocity profile was imposed as a Dirichlet condition at the inlet column (*x* = 0) to drive the flow (Zou & He, 1997),

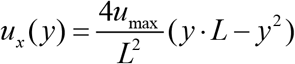

and a zero-gradient (first-order extrapolation) outflow condition was applied at the opposite column (*x* = *L*). The lateral boundaries were therefore of inlet/outlet type rather than periodic. Convergence to steady state required three criteria to be met simultaneously, and to remain met at three consecutive monitoring intervals: (i) the relative change of the ECS-averaged flow speed between monitoring intervals fell below 10⁻⁶; (ii) the normalised velocity-field residual of the MY+ fluid domain fell below 10⁻⁵; and (iii) the mean step-wise relative change of the MY+ velocity field, averaged over the last three monitoring intervals, fell below 10⁻⁵.

#### 2.2.2 Derived Flow-Field Diagnostics

From each converged velocity field, we computed a set of derived diagnostics. The local Péclet and Reynolds numbers were defined as

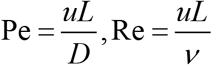

Where 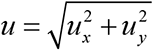 is the local flow speed converted to physical units, *L* is the local ECS channel width, *D* = 1 × 10⁻⁹ m² s⁻¹ is a representative small-molecule diffusivity and *ν* = 1 × 10⁻⁶ m² s⁻¹ is the kinematic viscosity of water at body temperature. Note that this *ν* is the physical viscosity used for diagnostic purposes and is distinct from the lattice viscosity *ν*_LB_ of the BGK solver. Because *D* and *ν* are both held fixed, Pe and Re differ only by the constant factor *ν*/*D* (a Schmidt number of ≈ 1000) throughout this study, and the two maps are therefore informationally equivalent; both are reported because they address different questions (advection versus inertia). The local ECS channel width was obtained from a Euclidean distance transform (EDT) of the pore space as twice the distance to the nearest membrane, *L* = 2 · EDT. Local vorticity was computed as

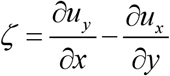

using second-order central differences in the interior of the domain. The kinetic-energy spectrum *E*(*k*) was obtained by two-dimensional Fourier transform of the velocity field followed by radial binning in wavenumber *k*; a power-law exponent was fitted to the log–log spectrum and compared with the *k*^−5/3^ scaling predicted for the inverse energy cascade of two-dimensional turbulence (Kraichnan, 1967) as a purely qualitative flow-regime check. The inlet column at *x* = 0, where the Dirichlet condition fixes the velocity, was excluded from all ECS-masked diagnostics (Pe, Re, vorticity), since it reflects the imposed boundary condition rather than the flow response of the geometry.

#### 2.2.3 Quandt-Davis Model for Myelin Capacitance and Conductance

The electrical effect of the myelin sheath was represented following the Quandt–Davis (Q–D) treatment of multi-lamellar internodal membrane (Quandt & Davis, 1992). Internodal segments were modelled as passive cable with wrap-dependent specific capacitance and leak conductance:

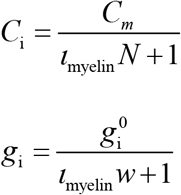

where *C*_m_ = 1.0 μF cm⁻² is the bare-membrane specific capacitance, 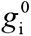 is the bare internodal leak conductance (3 × 10⁻⁴ S cm⁻² for pyramidal axons and 3 × 10⁻⁵ S cm⁻² for interneuron axons), *N* is the structural wrap count and *ι*_myelin_ is a dimensionless myelin index representing the electrical completeness of the sheath. The myelin index increased monotonically with wrap count, from *ι*_myelin_ = 0.125 at *N* = 2 to *ι*_myelin_ = 1.0 at the maximal wrap count for pyramidal axons (*N* = 16), and from *ι*_myelin_ = 0.0625 at *N* = 2 to *ι*_myelin_ = 1.0 at the maximal wrap count for interneuron axons (*N* = 8); it could also be overridden directly to impose a prescribed time course of demyelination (Sections 4 and 6). The surface-area factor 2π·*δ*_a_·*L* carried by the original Quandt–Davis expressions for total internodal capacitance and conductance is not applied explicitly, because NEURON scales specific membrane properties by the surface area it computes from the diameter and length of each segment.

#### 2.2.4 Electrophysiological Modelling with NEURON

Neuronal electrophysiology was simulated in NEURON (Carnevale & Hines, 2006; Hines & Carnevale, 1997) using detailed morphological reconstructions obtained from NeuroMorpho.Org. In the single-cell and single-synapse simulations of Sections 3 and 4 every neuron carried its full reconstructed morphology. In the 200-cell network simulations of Sections 5 and 6 a spatially graded compromise was required to keep the simulations tractable: neurons whose soma lay inside the 10 × 10 × 250 μm core volume surrounding the recording electrodes, which are the cells that dominate the extracellular signal, carried their full reconstructed morphology, whereas neurons outside that volume were reduced to an equivalent-cable representation whose somatic and axonal diameters were drawn from the statistics of the same reconstructions with independent jitter; glial cells retained their full morphology throughout. Imposing the full morphology on every neuron in a reduced twenty-cell configuration raised mean pyramidal firing rate approximately three-fold, in the same direction in all three network seeds tested (Supplementary Fig. 8f). In that reduced configuration the rate scaled monotonically with the proportion of pyramidal cells retaining their full morphology, so the three-fold figure brackets the magnitude of the effect rather than calibrating the offset of the production configuration, in which the proportion is both larger and more stable. The direction being consistent, morphological detail acts as an offset of the baseline rate; since every comparison reported in this work is either between groups simulated under an identical morphological configuration or between phases within one network, that offset does not enter any of the reported contrasts. Myelinated axons were built as alternating node-of-Ranvier and internodal sections. Nodes were represented by single segments (nseg = 1); internodes were discretised with an odd number of segments scaling with internodal length, nseg = max(3, 2⌊L/20 μm⌋ + 1), giving a segment length of ≤ 10 μm in passive internodal cable, which is short relative to the internodal electrotonic length constant.

Internodal length was set from axon calibre following Rushton’s proportionality between internode length and fibre diameter (Rushton, 1951; Waxman, 1980),

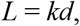

where *d* is the axon diameter and *k* = 80 for pyramidal neurons and *k* = 30 for interneurons; the resulting value was then rescaled by total fibre diameter (axon plus sheath) and capped at 500 μm.

Voltage-gated sodium and potassium conductances at nodes of Ranvier and at the axon initial segment (AIS) were described by Hodgkin–Huxley kinetics (Hodgkin & Huxley, 1952). Base nodal sodium conductance was 0.12 S cm⁻² for pyramidal axons and 0.75 S cm⁻² for interneuron axons. Consistent with the well-documented enrichment of Na⁺ channels at nodes and at the AIS relative to somatodendritic membrane (Caldwell et al., 2000; Kole et al., 2008), these base values were multiplied by an axonal enhancement factor of 2.5 (pyramidal) or 1.3 (interneuron), applied exclusively to axonal Hodgkin–Huxley compartments (nodes, AIS and unmyelinated axon) and never to somatic or dendritic membrane, yielding effective nodal densities of 0.30 and 0.975 S cm⁻² respectively. For heavily myelinated pyramidal axons (*N* ≥ 10) the nodal sodium conductance received a further 1.22-fold increase before axonal scaling. Nodal and AIS potassium conductance was 0.3 S cm⁻²; somatodendritic potassium conductance was 0.35 S cm⁻² in pyramidal cells and 0.3 S cm⁻² in interneurons. All simulations were run at 37 °C.

Synaptic transmission was implemented with two-state (dual-exponential) conductance-based synapses (Destexhe et al., 1994). Synaptic kinetics were assigned by presynaptic cell class: excitatory (pyramidal presynaptic) synapses had *τ*₁ = 5.0 ms, *τ*₂ = 12.5 ms and a reversal potential of 0 mV; inhibitory (interneuron presynaptic) synapses had *τ*₁ = 5.0 ms, *τ*₂ = 35.0 ms and a reversal potential of −80 mV. These time constants are deliberately slower than those of isolated AMPA-and GABA_A_-receptor currents; they represent the compound, spatially distributed synaptic conductance seen at the soma of a morphologically detailed cell and were chosen so that single-synapse somatic postsynaptic potentials (PSPs) were reliably measurable over the analysis window. Baseline peak synaptic conductances were connection-type specific: 0.02 μS for pyramidal→pyramidal, 0.002 μS for pyramidal→interneuron and 0.2 μS for interneuron→pyramidal and interneuron→interneuron connections (Bezaire & Soltesz, 2013). In addition, an ECS-coupling mechanism scaled the effective presynaptic weight as a function of presynaptic myelination: the scaling factor was 1.0 at maximal myelination (*N* = 16 pyramidal, *N* = 8 interneuron) and decreased with a power-law exponent of 1.4 towards floors of 0.28 (excitatory) and 0.70 (inhibitory) at minimal myelination.

### 2.3 Simulation Protocols

#### 2.3.1 ECS Flow Analysis (**Section 1**)

ECS flow was simulated using LBM in 2D domains (20 × 20 μm) representing individual CA1 layers. Simulations were performed under two myelination conditions: MY-(2 layers for all axons) and MY+ (16 layers for pyramidal neurons, 8 layers for interneurons). The same geometry (cell positions and sizes) was used for both conditions, differing only in myelin thickness, enabling paired comparison. Flow was driven by a parabolic velocity profile at the inlet with Reynolds number Re = 0.01. Simulations ran until convergence criteria were met: velocity residuals < 10^-6^ and mean ECS velocity change < 10^-5^ over 8 consecutive check intervals. Outputs included velocity fields, porosity (MY-vs. MY+), ECS volume fraction, and mean ECS velocity. Parameter Sensitivity Analysis (Section 1). To assess the robustness of the geometry-flow relationship independent of the demyelination manipulation, we examined how the pooled velocity-reduction effect scaled with cell packing density (linear regression) and how mean ECS flow velocity scaled with ECS volume fraction (power-law regression on log-transformed values) across all 32 simulations, pooling both myelination conditions and all four sublayers.

#### 2.3.2 Substance Transport Analysis (Section 2)

Molecular transport through the ECS was quantified by pore-scale homogenisation (Whitaker, 1999), applied to exactly the same 32 paired MY+/MY− binary ECS geometries used in Section 1 (8 per layer), giving 64 independent closure solutions. For each geometry and each Cartesian direction *d* ∈ {*x*, *y*} we solved the periodic diffusion closure ("cell") problem within the ECS pore space,

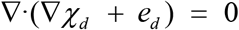

with zero normal flux at all cell-membrane interfaces and periodicity at the domain edges. The problem was discretised as a graph Laplacian over pore-space nodes — obstacle faces carry no conductance, which imposes the zero-flux membrane condition exactly — and solved by preconditioned conjugate gradients to a relative tolerance of 10⁻⁶ (a direct sparse solve was available as a cross-check). Direction-resolved geometric factors were recovered in energy form,

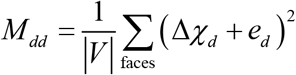

and combined with the ECS volume fraction *α* (the pore-pixel fraction) to give the direction-resolved effective diffusion coefficients and tortuosity components, tortuosity being defined as in the classical brain-ECS diffusion literature (Nicholson & Phillips, 1981; Syková & Nicholson, 2008)

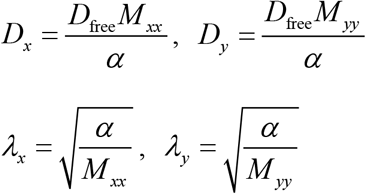

using a free glutamate diffusivity *D*_free_ = 5 × 10⁻¹⁰ m² s⁻¹ (Longsworth, 2002; Rusakov & Kullmann, 1998). The isotropic geometric factor 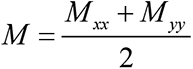 gives the overall 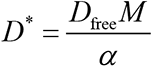 and 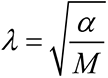 reported as the primary metrics in Fig. 2, while the anisotropy induced by the aligned myelinated-axon geometry was quantified by the diffusivity ratio *D_x_*/*D_y_* and by the geometric anisotropy index

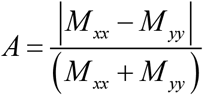

**Figure 2.**
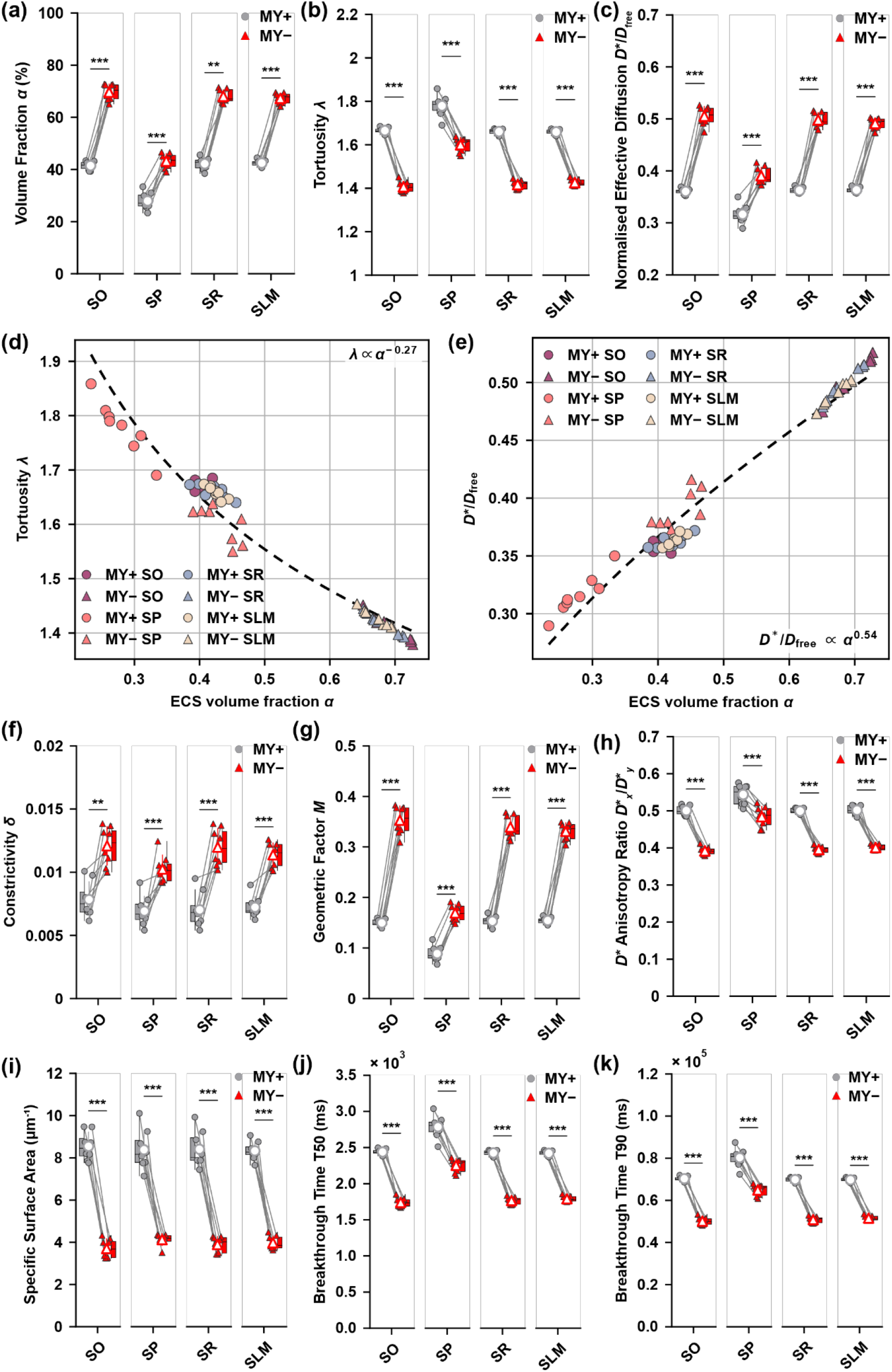
Demyelination-driven ECS expansion accelerates glutamate transport primarily, but not exclusively, through increased volume fraction. Pore-scale homogenization and advection-diffusion breakthrough simulations were performed on the same eight paired MY+/MY− geometries per sublayer as in Figure 1, modelling glutamate (*D*_free_ = 5 × 10⁻¹⁰ m²/s) as the diffusing species. Unless otherwise noted, all panels compare the four CA1 sublayers (SO, SP, SR, SLM) under myelinated (MY+, grey circle) and demyelinated (MY−, red triangle) conditions (*n* = 8 independent geometries per sublayer, paired *t*-test; \*\**p* < 0.01, \*\*\**p* < 0.001). (a) Homogenized ECS volume fraction *α*. (b) Homogenized tortuosity *λ*. (c) Normalized effective diffusion coefficient *D*\*/*D*_free_. (d) Tortuosity *λ* plotted against ECS volume fraction *α*, pooled across all sublayers and both myelination states (circle, MY+; triangle, MY−; colored by sublayer). A power-law fit (*λ* ∝ *α*^−0.27^, *R*² = 0.956) captures most of the variance, but an extended model including sublayer and myelination-state terms fit significantly better (*F*(5,57) = 57.3, *p* < 0.0001), indicating that *α* is the dominant but not sole determinant of tortuosity. (e) Normalized effective diffusion coefficient *D*\*/*D*_free_ plotted against *α*, same dataset and analysis as (d) (*D*\*/*D*_free_ ∝ *α*^0.54^, *R*² = 0.956; extended-model comparison *F*(5,57) = 57.3, *p* < 0.0001). (f) ECS constriction ratio *δ* = (P10/P90)² of the local channel-width distribution, where a higher *δ* indicates a more uniform, less constricted geometry. (g) Geometric factor M relating *D*\* to *α* 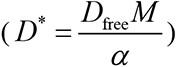. (h) Directional anisotropy of effective diffusion, *D*_x_/*D*_y_. (i) Specific surface area (SSA, μm⁻¹) of the ECS-membrane interface. (j) Time to 50% breakthrough (T50) from constant-source (step-concentration) breakthrough simulations across the domain width. (k) Time to 90% breakthrough (T90) from the same simulations as (j); T50 and T90 changed by an identical fold-ratio (0.75) between MY+ and MY−, consistent with both deriving from a single underlying diffusion time constant rather than representing independent measures. To test whether the ECS geometric expansion observed following demyelination (Figure 1) translates into faster molecular transport, we performed pore-scale homogenization and advection-diffusion breakthrough simulations on the same eight paired MY+/MY− geometries per sublayer used in Figure 1, modeling glutamate (*D*_free_ = 5 × 10⁻¹⁰ m²/s), the principal excitatory neurotransmitter in CA1, as the diffusing species.

Local ECS channel width was estimated at every pore pixel as twice the Euclidean distance to the nearest membrane obstacle; the coefficient of variation (CV), 10th percentile (P10) and 90th percentile (P90) of this per-pixel distribution were computed per geometry, and constrictivity was defined as δ = (P10/P90)^2^

Specific surface area (SSA) was computed as the length of the ECS–membrane interface normalised by ECS pore area. Point-to-point breakthrough times were obtained analytically from the isotropic effective diffusion coefficient using the one-dimensional semi-infinite-medium solution

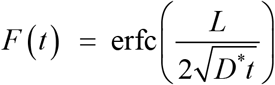

where *L* is the domain length along the transport direction; T10, T50 and T90 are the times at which F(t) = 0.1, 0.5 and 0.9, obtained by inverting the complementary error function. This analytic route was used in preference to direct particle tracking or transient concentration-field simulation because it is exact for the homogenised (REV-averaged) diffusion tensor and avoids the substantial cost of resolving pore-scale concentration fields over transport-relevant length and time scales. Accordingly, the breakthrough curves reported here should be read as the homogenised-medium transport response of each geometry, not as the concentration history of a localised point release.

Because this analysis reuses the Section 1 geometries and introduces no additional free or fitted parameter beyond the literature-based glutamate diffusivity (which enters purely as a known multiplicative scaling of *D*\* and hence of the breakthrough times), no separate parameter-sensitivity analysis was performed for this section; its robustness is inherited from the numerical and parametric checks already carried out on those geometries in Section 1, namely the grid-independence comparison and the solver-convergence test described for that section, together with the sensitivity of the converged flow to extracellular volume fraction and cell density across the thirty-two geometries (Supplementary Fig. 7a,b).

#### 2.3.3 Single-Synapse Postsynaptic Potential Analysis (Section 3)

Single-synapse PSPs were analysed for four connection types: pyramidal→pyramidal (P–P), pyramidal→interneuron (P–I), interneuron→pyramidal (I–P) and interneuron→interneuron (I–I). Each connection type was simulated across a gradient of presynaptic myelin wrap counts: 2, 4, 6, 8, 10, 12, 14 and 16 for pyramidal axons, and 2, 4, 6 and 8 for interneuron axons. Presynaptic action potentials were evoked by somatic current injection (duration 0.3 ms; amplitude 6.0 nA for pyramidal cells, 3.0 nA for interneurons) and PSPs were recorded at the postsynaptic soma. Each trial contained three stimulus epochs: an isolated single pulse at 50 ms, a pulse pair at 150 and 152 ms (inter-stimulus interval, ISI = 2 ms) used to quantify temporal summation, and an eight-pulse train beginning at 250 ms with a 2 ms inter-pulse interval used to probe high-frequency transmission reliability. Simulations ran for 500 ms with a fixed time step of 0.025 ms. Each condition was repeated 8 times with independent parameter randomisation (see **Biological Variability**). Layer-specific dynamic adjustment of resting and reversal potentials was applied on the basis of the ECS volume increase associated with demyelination, interpolated from the Section 1 results where available and otherwise falling back on layer-specific factors spanning +3 % (SP) to +35 % (SR). Reported outputs included PSP amplitude, rise time, decay time, half-width, duration, maximum rate of rise, spectral and energy descriptors, temporal-summation ratio, axonal conduction velocity, terminal action-potential amplitude, action-potential safety factor and synaptic transmission success.

##### Parameter sensitivity analysis (**Section 3**)

To confirm that the single-synapse results were not contingent on a knife-edge parameter choice, we performed a one-at-a-time (OAT) sensitivity sweep on a reduced ball-and-stick pyramidal cell receiving repetitive excitatory synaptic input. Seven parameters were each varied independently over five discrete levels around baseline: potassium conductance gK, 0.20–0.50 S cm⁻² (baseline 0.35); sodium conductance gNa, 0.8–1.6 S cm⁻² (baseline 1.2); synaptic weight, 0.01–0.08 μS (baseline 0.03); number of presynaptic input events, 2–12 (baseline 5); inter-event interval, 1–15 ms (baseline 2.0); synaptic decay time constant *τ*₂, 2–12 ms (baseline 5.0); and number of synapses, 1–8 (baseline 4). The resulting change in somatic action-potential count and peak amplitude relative to baseline was ranked by effect size. Discharge was governed almost entirely by the repolarising and depolarising conductances: the range of action-potential counts spanned by the potassium conductance was twice that spanned by the sodium conductance, and each of the synaptic parameters spanned less than a sixth of the potassium range, with the synaptic decay time constant and the number of input events having no detectable effect. The single-synapse results therefore do not rest on the particular synaptic settings adopted. We additionally verified the robustness of the transmission-success classification to the presynaptic release-threshold working point (−34 mV for pyramidal and −28 mV for interneuron presynaptic terminals) and to the integration time step Δ*t* (0.0125, 0.025 and 0.05 ms, tested for P–P and I–I connections), and computed Wilson score 95 % confidence intervals on the transmission success rate at each wrap count for all four connection types.

#### 2.3.4 PSP Temporal Evolution Analysis (Section 4)

The temporal evolution of PSPs was analysed with the same single-pulse and paired-pulse protocols as Section 3, but under a continuously varying myelination state comprising five consecutive phases (initial, degradation, minimum, recovery, final). Single-pulse stimulation assessed baseline PSP properties within each phase, while paired-pulse stimulation (ISI = 2 ms, matching Section 3) quantified temporal summation and short-term facilitation/depression. Presynaptic action potentials were again evoked by somatic current injection (0.3 ms; 6.0 nA pyramidal, 3.0 nA interneuron). Each phase occupied 500 ms and total simulation duration was 2500 ms with a time step of 0.025 ms. The same four connection types and wrap-count gradients as Section 3 were tested. The myelin index was driven from 1.0 (MY+) down to 0.125 (MY−, i.e. 2 wraps) and back to 1.0, with layer-specific dynamic adjustment of resting and reversal potentials tracking the associated ECS volume changes. Each condition was repeated 8 times. Interneuron morphologies were sampled from two reconstructions (TF35 axo-axonic and DL43j). The essential distinction from Section 3 is therefore temporal: Section 3 holds myelination fixed (static analysis), whereas Section 4 imposes a time-dependent degradation-and-recovery trajectory (dynamic analysis). Reported outputs included PSP amplitude and timing metrics, temporal-summation ratio, PSP spectral and energy descriptors, action-potential conduction delay and velocity, and phase-resolved summaries of all of the above.

#### 2.3.5 Static Network Extracellular Electrophysiology (Section 5)

Network-level extracellular electrophysiology (spikes and LFP) was computed for four experimental groups defined by the factorial combination of inflammatory state and myelination: normal composition with high myelination (N-High; MY+: 16 wraps pyramidal, 8 wraps interneuron), normal composition with low myelination (N-Low; MY−: 2 wraps for both classes), chronic inflammatory composition with MY+ (CI-High), and chronic inflammatory composition with MY− (CI-Low). Each network contained 200 cells (neurons plus glia) distributed across the four CA1 layers within a 20 × 20 × 500 μm domain. Cell composition differed between inflammatory states. The inflammatory census implements the reactive gliosis that is common to chronic neuroinflammatory conditions of the hippocampus, in which astrocytes and microglia expand while oligodendrocytes and neurons contract; the specific proportions were taken from quantitative hippocampal cell-density measurements made under sustained psychosocial stress(Czéh et al., 2006; Tynan et al., 2010), which we use here purely as a calibrated instance of that generic glial signature rather than as a model of any single disease aetiology. Normal networks comprised 55 % neurons, 24 % astrocytes, 16 % oligodendrocytes and 5 % microglia, whereas chronic inflammatory networks comprised 50 % neurons, 28 % astrocytes, 14 % oligodendrocytes and 8 % microglia. The inflammatory state acts on the network through two routes only: this census, and a shift of microglia to an activated phenotype (resting potential −70 to −65 mV, membrane time constant 50 to 30 ms, input resistance 500 to 300 MΩ, doubled calcium factor). The intrinsic parameters of pyramidal cells, interneurons, astrocytes and oligodendrocytes, the synaptic weights and the myelin parameters are identical in the two states, and the extracellular-space mechanism described below is driven by myelin wrap count and cortical layer rather than by cellular composition, so the inflammatory factor does not enter the neurotransmitter-dilution pathway. The inflammatory arm of the factorial design is therefore a test of whether the glial census alone reorganises network electrophysiology, and not a model of the signalling consequences of gliosis. Networks were simulated for 5000 ms with a time step of 0.025 ms at 37 °C. An absolute refractory period of 2 ms was enforced at spike detection to exclude non-physiological high-frequency discharge. Dynamic resting-and reversal-potential adjustment was applied according to myelin-dependent ECS volume changes as in Sections 3 and 4. Rather than injecting independent Poisson current into every cell, the network was driven by a shared Schaffer-collateral-like excitatory input at 5 Hz per pyramidal target, whose effective weight was modulated by the ECS mechanism described above. Recurrent connectivity used distance-dependent connection probability and weight, both decaying exponentially with inter-somatic distance (length constant 300 μm). Interneuron morphology was restricted to a single reconstruction (TF35 axo-axonic) to remove morphology as a confound. Extracellular potentials were computed with LFPy/LFPykit using the line-source approximation with an extracellular conductivity of *σ* = 0.3 S m⁻¹(Hagen et al., 2018; Holt & Koch, 1999; Lindén et al., 2014), recorded at four contacts arranged as a tetrode within stratum pyramidale at *z* = −80 μm. Eight independent repeats were run per group. Reported outputs included spike trains and firing-rate statistics, interspike-interval statistics, LFP time series and spatial distributions, power spectral density and band power, time–frequency representations, cross-frequency coupling, spike–LFP coupling, connectivity, nonlinear-dynamics descriptors and neural-manifold metrics.

#### 2.3.6 Dynamic Network Extracellular Electrophysiology (Section 6)

Dynamic network evolution was simulated as a single continuous trajectory through five phases: initial (intact myelination) → degradation → minimum (maximal demyelination) → recovery → final (restored myelination). The network contained 200 cells (neurons plus glia) in a 20 × 20 × 500 μm domain. Each phase lasted 5000 ms, for a total simulation duration of 25 000 ms with a time step of 0.025 ms at 37 °C. Cell composition was held at the chronic inflammatory census throughout (50 % neurons, 28 % astrocytes, 14 % oligodendrocytes, 8 % microglia; the CI composition defined in Section 5) so that the phase sequence isolates the effect of the myelination and ECS trajectory rather than confounding it with a simultaneous change in cellular census; the myelin index was driven dynamically from 1.0 (normal) to 0.125 (demyelination) and back to 1.0, with oligodendrocyte number tracking the myelin index. Layer-specific dynamic adjustment of resting and reversal potentials followed the ECS volume increase associated with demyelination. Spike detection (2 ms absolute refractory period), Schaffer-collateral-like 5 Hz drive, TF35-only interneuron morphology, and LFP computation (line-source, *σ* = 0.3 S m⁻¹, four contacts in stratum pyramidale) were identical to Section 5, and eight independent repeats were run. Reported outputs matched Section 5, with the addition of phase-resolved and phase-transition summaries.

### 2.4 Analysis and Validation

#### 2.4.1 Grid Independence Test

Grid independence of the LBM solution was assessed by repeating the flow simulation on the same geometry (identical cell positions and calibres) at two lattice resolutions that were carried to full convergence, 0.02 and 0.01 μm per lattice node. The number of iterations required to satisfy the three-part convergence criterion scaled from 365 300 to 690 500 steps, a factor of 1.89 for a two-fold refinement, which is the behaviour expected for diffusive relaxation of the velocity field and indicates that the solution is converging in a controlled manner rather than changing regime with resolution. A third, finer resolution of 0.005 μm was set up but was not carried through to convergence within the available compute budget, and is excluded from the comparison for that reason. Production runs used 0.01 μm per lattice node.

#### 2.4.2 Extracellular Signal Definition and Preprocessing

The extracellular potential computed by LFPy/LFPykit at each tetrode contact was stored at the integration rate of the simulation (40 kHz) as a single broadband trace. This trace is the model counterpart of the wideband voltage delivered by an in vivo headstage, and every extracellular measure reported here was derived from it through an explicit filter chain rather than read out directly, following standard extracellular-recording practice (Buzsáki et al., 2012). The LFP band was obtained with a sixth-order Butterworth low-pass at 300 Hz applied in the forward–backward (zero-phase) sense and then decimated to 1 kHz; the low-pass doubles as the anti-alias filter for the decimation, since 300 Hz lies well below the 500 Hz Nyquist frequency of the decimated signal. The spike band was obtained with a fourth-order zero-phase Butterworth band-pass at 300–3000 Hz and retained at the native 40 kHz, which resolves the biphasic extracellular waveform over the ±0.75 ms window used to display it in both sections. Zero-phase filtering was used throughout so that band-limited features are not shifted in time relative to the spike times, which matters for the spike-triggered measures below.

The first 200 ms of every trace was discarded before analysis. The network starts from a uniform initial condition, and the resulting settling of the extracellular baseline is a large low-frequency transient of numerical rather than physiological origin that would otherwise dominate spectral and entropy estimates over the early part of the record. In Section 5 this leaves 4800 ms per repeat; in Section 6 it removes the first 200 ms of the initial phase only, leaving the four subsequent phases intact.

Because the model contains no exogenous background activity, the simulated LFP lacks the ongoing broadband background of an *in vivo* recording and its absolute amplitudes are correspondingly small. The detectability of the phase effects against such a background was therefore tested explicitly rather than assumed, by adding a surrogate aperiodic 1/f background with superimposed theta and gamma peaks at a range of amplitudes relative to the in-band signal power and re-testing every band effect at each amplitude; this analysis, which is a worst-case bound because the surrogate is held constant across phases whereas the tissue contributing it in a real recording would itself be demyelinated, is reported in the supplementary material.

#### 2.4.3 Spike Train Analysis

Spike times were taken at the distal axonal recording site for pyramidal cells, which is the site that governs synaptic release, so that conduction failure along a demyelinated axon registers as a loss of output rather than being counted from a soma that still fires; interneurons were detected somatically throughout. Detection during the simulation used an upward crossing of −20 mV with a 15 mV hysteresis, so that an event is counted only once the compartment has repolarised below −35 mV, together with a 2 ms floor on the interval between counted events. At analysis time a single uniform convention merged events separated by less than 3 ms, retaining the first of each group.

The 3 ms window is a counting convention rather than an artefact-rejection step, and the distinction governs how the absolute rates should be read. The merged events are complete action potentials rather than detection artefacts: within depolarised episodes pyramidal cells discharge repetitively, each event showing a full upstroke from about −70 mV, an overshoot to +10 to +15 mV and complete repolarisation. The pyramidal firing rate is therefore a function of the counting window rather than an intrinsic property of the cell, and the rates reported here are to be read together with this convention. Interneurons, detected somatically, do not enter this regime and their rates are almost independent of the window, which provides an internal control. Because the fast discharge occurs only within short depolarised episodes, rates averaged over the analysis window remain physiological: at the 2 ms window the highest single-cell rate was 76.8 Hz, with a 99th percentile of 57 Hz.

The window does not carry any of the conclusions. Sweeping it from 2 to 20 ms leaves the main effect of myelination significant at every value, with the pyramidal partial *η*² rising monotonically from 0.468 to 0.985, and leaves the effect of inflammatory state non-significant at every value (*p* = 0.63 to 0.996); only the absolute scale of the rates changes. This sweep is reported in the supplementary material.

All firing rates, interspike-interval statistics and spike-triggered measures use the merged trains. Firing rate was computed per cell over the analysis window and summarised per repeat as the mean across cells of a given class. Interspike-interval distributions were characterised by their coefficient of variation, spike-count variability by the Fano factor over 100 ms windows (per cell and for the pooled population count), and temporal structure by the autocorrelogram, all computed separately for pyramidal cells and interneurons.

Extracellular spike waveforms were obtained from the spike band of the reference contact by averaging, for every cell contributing at least ten spikes, the segments centred on that cell’s spike times to form a single-cell template, and then averaging those templates with equal weight across cells and repeats. Averaging in two stages rather than pooling all spikes prevents the few most active cells from dominating the grand mean. Because the tetrode records a mixed population at a fixed location, a waveform recovered in this way is not guaranteed to be above the detection limit, so the amplitude of each grand mean was compared against a within-trace null: the peak-to-peak amplitude over the central 30 % of the window was divided by the peak-to-peak amplitude over the outer 40 %, where no spike-locked component should exist. A ratio near unity indicates that no waveform was resolved, and this ratio is reported alongside the waveforms rather than the waveforms being shown without qualification.

#### 2.4.4 LFP Spectral and Coupling Analysis

Frequency bands followed the convention used for the rodent hippocampus: delta 0.5–4 Hz, theta 4–12 Hz, beta 12–30 Hz and gamma 30–80 Hz. The theta band is wider than in human EEG conventions because hippocampal theta in the behaving rodent spans roughly 4–12 Hz and shifts with locomotor state (Buzsáki, 2002; Colgin, 2013).

Power spectral density was estimated with Thomson’s multitaper method (Thomson, 1982) using discrete prolate spheroidal sequences (Slepian, 1978) with time–bandwidth product NW = 6 and K = 2NW − 1 = 11 tapers, applied to the whole analysis window rather than to overlapping segments. On records of this length — 4.8 s in Section 5 and 5 s per phase in Section 6 — Welch’s method yields only about nine half-overlapping 1 s segments, so its variance is comparable in magnitude to the between-group differences of interest. The multitaper estimate has approximately 2K equivalent degrees of freedom and substantially lower spectral leakage, at the cost of a frequency resolution of 2NW/T, i.e. 2.5 Hz here, which is adequate for the broadband spectra observed. Band power was obtained as the Riemann sum of the spectral density over each band.

Time–frequency representations used the complex Morlet continuous wavelet transform (Torrence & Compo, 1998) with ω₀ = 6 over 96 logarithmically spaced frequencies from 0.5 to 120 Hz, reported as |*W*(*f*, *t*)|². Because this quantity is not calibrated to a spectral density, spectrograms are displayed on a linear power scale in arbitrary units rather than in decibels, so that the relative magnitude of band-limited power changes is preserved against the broadband baseline.

Spike–LFP coupling was quantified in four ways, each computed separately for pyramidal cells and interneurons with spikes pooled across cells of a class. The spike-triggered average was taken over a ±100 ms window around each spike. Spike-field coherence was computed as the ratio of the power spectrum of the spike-triggered average to the mean power spectrum of the individual spike-triggered segments (Fries et al., 2001), over ±128 ms windows and with a Hann taper; each segment was mean-subtracted before windowing, because at this window length the segments are otherwise dominated by the slow baseline they share by virtue of lying on the same population LFP trajectory, which inflates the ratio at low frequencies without reflecting spike-locked structure. Phase locking was assessed in the theta and gamma bands from the Hilbert phase of the band-passed LFP sampled at each spike time, and reported as the phase-locking value, as the pairwise phase consistency (Vinck et al., 2010) — which, unlike the phase-locking value, is not biased upwards as spike count falls — and as the preferred phase. Phase–amplitude coupling was quantified by the Tort modulation index (Tort et al., 2010) between theta phase and gamma amplitude over 18 phase bins. Where a cell class produced more than 8000 spikes in a repeat, a random subsample of that size was used for the two segment-based measures, which bounds the size of the segment matrix without materially affecting estimates that have already converged at that sample size. Where these coupling measures are compared across the five phases of Section 6, a further precaution is required, because firing rate itself falls during demyelination and both spike-field coherence and the phase-locking value are biased upwards as spike count decreases. The spike count entering each estimate was therefore equalised across phases within each repeat, by drawing a random subsample of the size of the sparsest phase for that cell class (1720– 3964 spikes for pyramidal cells and 148–468 for interneurons across the eight networks), so that any residual bias is common to all five phases. The adequacy of this correction was checked by recomputing every estimate without equalisation. Equalisation shifted the phase-locking value systematically upwards and most strongly where spikes were scarcest, by up to 18 % for interneuron theta, whereas its effect on the pairwise phase consistency was an order of magnitude smaller and of inconsistent sign, as expected of a measure whose expectation does not depend on spike count.

Interactions between recording sites were computed across the six pairs of tetrode contacts, on the LFP band decimated to 500 Hz, as magnitude-squared coherence, cross-correlation, and the phase lag index (Stam et al., 2007) together with its weighted variant (Vinck et al., 2011); the latter two are insensitive to the zero-lag component contributed by volume conduction.

#### 2.4.5 Nonlinear Dynamics and Neural Manifold Analysis

Complexity of the LFP band, decimated to 250 Hz, was characterised by thirteen descriptors computed over the whole analysis window: the Hurst exponent by rescaled-range analysis (Hurst, 1951; Mandelbrot & Wallis, 1995); wavelet entropy from the relative energies of a four-level db4 discrete wavelet decomposition (Rosso et al., 2001); permutation entropy of order 3 (Bandt & Pompe, 2002) and the corresponding permutation Lempel–Ziv complexity; Higuchi’s fractal dimension with *k*_max_ = 10 (Higuchi, 1988); the Shannon entropy of the amplitude histogram; approximate entropy (Pincus, 1991) and sample entropy (Richman & Moorman, 2000), both with m = 2 and r = 0.2 SD; Hilbert–Huang spectral entropy of the empirical-mode decomposition; Lempel–Ziv complexity of the median-binarised series (Lempel & Ziv, 1976); and the recurrence rate, determinism and laminarity of a recurrence quantification analysis (Marwan et al., 2025). The amplitude-histogram Shannon entropy, the permutation entropy, the wavelet entropy and the Hilbert–Huang spectral entropy were each normalised to the unit interval by their maximum attainable value (log₂ of the bin, pattern, subband and frequency-bin count, respectively). Approximate and sample entropy are differences of logarithmic correlation sums and have no finite upper bound, so they are reported in their native units, as are the Hurst exponent and the fractal dimension; the descriptors are therefore not all on a common scale, and cross-descriptor comparison is made on standardised change rather than on raw values (see Statistical Analysis). Two of the thirteen were found to be degenerate on these data and are reported only in the effect-size summary, not as panels: the recurrence rate is fixed at 5 % by construction of the recurrence threshold and therefore carries no measurement information, and the median-binarised Lempel–Ziv complexity takes only eight distinct values across the thirty-two runs. The remaining eleven are reported in full, each on its own native scale. For the composite figure panel the three descriptors whose phase dependence survives multiple-comparison correction — the Hurst exponent, approximate entropy and permutation entropy — are drawn together as the standardised paired change *d_z_* relative to the initial phase, which is what allows measures carried on incommensurable native scales (the Hurst exponent on the unit interval, approximate entropy as an unbounded log-ratio near 1.7, permutation entropy just below its normalised ceiling) to be read on one axis.

Population activity was projected onto a low-dimensional manifold from the pyramidal spike trains. Spike counts were binned at 50 ms and converted to rate, then smoothed across bins with a Gaussian kernel of *σ* = 2 bins, an effective width of about 100 ms; the first 800 ms and the last 300 ms of the smoothed series were discarded, the former because the network starts from rest and has not yet reached steady discharge, the latter because the zero-padded convolution systematically depresses values within 3*σ* of either end. Because the number of active pyramidal cells differs between groups (87 to 101), per-cell rate vectors are not comparable across groups, so each time bin was instead reduced to seven descriptors of the instantaneous population rate distribution that are independent of cell count and cell identity: the across-cell mean, the standard deviation, the fraction of cells firing above 0.5 Hz, and the 25th, 50th, 75th and 90th percentiles. These descriptors from all groups and all repeats were pooled, standardised and submitted to a single principal component analysis, yielding one common three-dimensional basis (69.4, 26.1 and 3.0 % of variance) in which every trajectory was expressed. Per-group principal component analysis followed by Procrustes superimposition was considered and rejected, because Procrustes normalises both configurations to unit Frobenius norm and matches them by optimal rotation and scaling, thereby removing the scale and shape differences that carry the group effect.

Four measures were computed per repeat in this common space. All four are distances, or ratios of distances, within a shared low-dimensional state space; this is the standard framing of representational geometry, in which conditions are compared through the distances between their population response patterns (Kriegeskorte et al., 2008; Kriegeskorte & Wei, 2021), and the low-dimensional projection on which it rests is established practice for population recordings (Cunningham & Yu, 2014). The displacement of the population state was the Euclidean distance from the trajectory centroid to the centroid of the healthy reference condition (normal, MY+); for the reference condition itself a leave-one-out centroid of the remaining repeats was used, since comparing a repeat against a mean that contains it otherwise fabricates an apparent effect of inflammation. Reproducibility was the distance from each repeat’s centroid to the leave-one-out centroid of the remaining repeats of the same condition, a quantity that for the reference condition coincides with the displacement measure by construction. State-space speed was the mean step length between consecutive time bins, and the net displacement ratio was the endpoint separation divided by the total path length, retained as a purely geometric control. Condition identity was decoded from the same seven population descriptors with a support vector machine (*C* = 1) under leave-one-repeat-out cross-validation, in which all time bins of one repeat are held out and the remaining seven repeats form the training set, with feature standardisation fitted on the training folds only.

Linear and radial-basis-function kernels (*γ* = 1/(*n*_features_ × variance)) were run over identical folds. Three problems were posed: myelination (two classes, chance 1/2), inflammatory state (two classes, chance 1/2) and condition (four classes, chance 1/4). Stratified *k*-fold cross-validation over pooled time bins was deliberately not used, because the rate matrices are smoothed over approximately 100 ms, so neighbouring bins are strongly correlated and random splitting places correlated samples on both sides of the partition, which inflates apparent accuracy.

Trajectory shape was examined separately, over the whole 5 s window at 100 ms bins with a Gaussian kernel of 1.5 bins. Four descriptors defined per repeat, the mean Frenet curvature, the mean absolute torsion, a path deviation measure (root-mean-square orthogonal distance of the trajectory from the straight chord joining its endpoints, in the units of the principal-component scores) and the net displacement ratio, were tested with the same 2 × 2 factorial analysis of variance used for the measures above, and three alignment-based summaries, the residual Procrustes disparity relative to a reference repeat, the mean pairwise length-normalised dynamic time warping distance and the cross-repeat variance of the aligned trajectories, were computed as one aggregate value per group and therefore not tested inferentially. No pairwise contrast survived correction for any of these descriptors, and the two that returned a main effect of myelination are not interpreted: the path deviation measure carries the units of the principal-component scores and, because its bins span the simulation from t = 0, scales with the overall extent of the trajectory rather than describing its shape, while curvature is numerically fragile because its denominator is the cube of the tangent norm and diverges where the trajectory momentarily stalls. The values are reported in Results. On these data the manifold signature of demyelination is a displacement of the population state, not a change in the geometry of its trajectory.

In Section 6, where the five phases are traversed within each network rather than compared between groups, the same population data support two further analyses. Pyramidal spike counts were binned at 100 ms and smoothed with a Gaussian kernel of *σ* = 1.5 bins, and each repeat’s rate matrix was projected onto its own first three principal components; for the pooled three-dimensional display the eight trajectories were brought into a common frame by Procrustes alignment to a reference repeat, which is a display convention and carries no quantitative claim. Three phase-resolved measures were then computed within each repeat and divided by that repeat’s root-mean-square radius about its global centroid, which renders them dimensionless and comparable across networks: the displacement of each phase centroid from the initial-phase centroid, the mean distance of time points to their own phase centroid, and the mean step length between consecutive bins. Because a centroid estimated from a finite number of bins fluctuates even when the underlying state does not, the reference point for the displacement measure is not zero but the distance between the centroids of two random halves of the initial phase, taken as the median over 200 splits; the four subsequent phases are tested against that null rather than against a fixed origin. These measures are invariant to the choice of principal-component basis, since distances within a repeat are unaffected by an orthogonal transformation of its own coordinates, and they involve no comparison of coordinates between repeats.

Decoding of phase identity from the instantaneous population state used the seven cell-count-independent descriptors defined above rather than principal-component coordinates, and no alignment. Per-repeat principal components are unsuitable for this purpose because each repeat obtains its own arbitrary basis, and they are comparable across networks only after alignment; Procrustes alignment, however, matches the ***i***th time bin of one repeat to the ***i***th time bin of the reference, and phase identity is a deterministic function of time bin, so the alignment would supply precisely the quantity the classifier is asked to recover. The population descriptors require no alignment, being directly comparable across networks by construction. Support vector classification used a leave-one-repeat-out outer loop, so that no time bin from a test network appears in training, with feature scaling fitted on the training folds only and hyperparameters selected by a nested grouped search over the training repeats alone. Chance level is 1/5. Linear and radial-basis kernels are reported side by side; they perform equivalently here, and the confusion matrix is assembled from the out-of-fold predictions of the outer loop so that its diagonal mean equals the reported accuracy by construction.

#### 2.4.6 Biological Variability

Inter-individual variability within the rat population was represented by parameter randomisation across repeats (Markram et al., 2015; Van Geit et al., 2008). Sodium, potassium and leak conductances, internodal leak conductance, membrane capacitance (somatic, dendritic and axonal), axial resistance, node length and passive dendritic parameters were each drawn independently from normal distributions with a coefficient of variation of 5 %. The afferent drive, by contrast, was not randomised across repeats: in Section 6 the Schaffer-collateral Poisson input is generated from a counter-based stream seeded by fibre index alone, so that fibres are mutually independent within a network but every repeat receives the identical input realisation. The design is therefore one in which a single afferent spike pattern is delivered to networks that differ in their intrinsic parameters, analogous to presenting the same stimulus to different animals, and the repeats are not independent realisations of the input. Because the paired phase-versus-initial differences consequently share a component determined by the common input realisation, they are positively correlated across repeats and the standard error of their mean is underestimated, so the post-hoc p-values reported for Section 6 should be read as anti-conservative. The effects that carry the Section 6 conclusions are large relative to this concern, but the borderline ones would require replication with per-repeat input seeds. Synaptic peak conductances were **not** randomised; they were set deterministically by connection type and by the ECS-coupling scaling described above. Synaptic time constants were likewise held fixed. Each parameter combination was repeated 8 times.

#### 2.4.7 Statistical Analysis

Statistical testing followed the design of each section. For Sections 1 and 2, where MY− and MY+ were solved on identical geometries, comparisons were paired: normality of the paired differences was assessed with the Shapiro–Wilk test (SHAPIRO & WILK, 1965) and the paired *t*-test or the Wilcoxon signed-rank test (Wilcoxon, 1992) was applied accordingly. For Section 3, where myelination conditions correspond to different parameter combinations rather than matched geometries, independent-sample comparisons used Welch’s unequal-variance *t*-test (Welch, 1947) or the Mann–Whitney *U* test(Mann & Whitney, 1947). For Section 4, phase-resolved metrics were compared against the initial phase within each connection type using paired t-tests.

For Section 5 the primary model was a 2 × 2 factorial ANOVA on myelination and inflammatory state treating the four groups as independent samples, which yields the two main effects and their interaction. Four pre-specified contrasts were then tested — the effect of inflammation at each myelination level and the effect of myelination in each inflammatory state — using Welch’s *t*-test with Holm–Bonferroni correction over that family of four (Holm, 1979); the two diagonal comparisons, which change both factors simultaneously, are not interpretable and were not tested. Effect sizes are reported as Hedges’ *g* with the small-sample correction (Hedges, 1981). Because the eight repeats of each group share the same Section 1 extracellular-space input and are therefore not strictly independent across groups, the same data were additionally analysed as a randomised complete block design with repeat as the blocking factor; the block effect was not significant for any reported metric and the two analyses agree, so the independent-samples model is reported as the more conservative of the two and the blocked analysis is tabulated as a robustness check.

For Section 6 the five phases are repeated measures on the same eight simulated networks, so the analysis was specified accordingly. Two omnibus tests are reported for every metric: the Friedman test with Kendall’s W (Friedman, 1937) as the primary, distribution-free test, and a one-way repeated-measures ANOVA with Greenhouse–Geisser correction (Greenhouse & Geisser, 1959) as a parametric counterpart. Sphericity was not pre-tested, since Mauchly’s test has very low power at *n* = 8; the Greenhouse–Geisser correction was applied unconditionally instead. Post-hoc comparisons of each phase against the initial phase used the paired t-test as the pre-specified primary test, with Holm–Bonferroni correction applied within each metric over the family of four phase-versus-initial contrasts; normality of the paired differences was verified with the Shapiro–Wilk test, and the Wilcoxon signed-rank test is computed alongside as a distribution-free robustness check. The *t*-test rather than the Wilcoxon test was chosen as primary because at *n* = 8 the two-sided Wilcoxon *p*-value cannot fall below 2/2⁸ = 0.0078, so with a family of four comparisons the smallest attainable Holm-corrected *p* is 0.031 — a resolution limit of the rank statistic at this sample size rather than a statement about the data. Where the paired differences fail the Shapiro–Wilk pre-check, as they do for firing rate in both cell classes, the Wilcoxon test is reported as primary for that metric and the t-test as the robustness check, with the same family of four. All significance annotations on phase-resolved panels are Holm-corrected; where the scientific question concerns a single pre-specified contrast the family was restricted accordingly rather than the test being changed after the fact. Effect sizes for these repeated-measures contrasts are reported as *d_z_*, the mean of the paired differences divided by their standard deviation (Lakens, 2013), which is the effect-size measure matched to the paired test and allows descriptors carried on different native scales to be compared on a common axis.

The significance level was 0.05 throughout, and significance is annotated as * *p* < 0.05, ** *p* < 0.01, *** *p* < 0.001.

#### 2.4.8 Implementation

All simulations and analyses were implemented in Python 3.10.11. The LBM fluid-dynamics solver (Sections 1–2), the homogenisation solver (Section 2), all analysis pipelines and all figure generation were written in Python using NumPy 2.2.6, SciPy 1.15.1, scikit-learn 1.6.1, pandas 2.2.3 and Matplotlib 3.10.0; the nonlinear-dynamics descriptors additionally used PyWavelets 1.8.0 for the wavelet decomposition, EMD-signal 1.6.4 for the empirical-mode decomposition and nolds 0.6.1 for the rescaled-range estimate of the Hurst exponent.

Compartmental electrophysiology (Sections 3–6) used NEURON 9.0 through its Python interface (Carnevale & Hines, 2006; Hines & Carnevale, 1997), with extracellular potentials computed via LFPy 2.3.5 and LFPykit 0.5.1 (Hagen et al., 2018; Lindén et al., 2013).

Simulations were executed on multi-core workstation and cloud compute nodes, with independent repeats run in parallel as separate processes and with the same package versions verified on every node. Simulated durations were: Sections 1–2, run to the convergence criterion rather than a fixed duration; Section 3, 500 ms; Section 4, 2500 ms (5 × 500 ms phases); Section 5, 5000 ms; Section 6, 25 000 ms (5 × 5000 ms phases).

## 3 Results

### 3.1 Demyelination Increases ECS Flow Velocity through Expanded Extracellular Volume

To directly test the geometric prediction that demyelination expands the extracellular space (ECS) and thereby facilitates interstitial bulk flow, we performed lattice-Boltzmann fluid simulations on eight independently generated CA1 microenvironment geometries, each evaluated in a fully myelinated (MY+) and demyelinated (MY−) configuration, and repeated across the four CA1 sublayers (stratum oriens, SO; stratum pyramidale, SP; stratum radiatum, SR; stratum lacunosum-moleculare, SLM; Figure 1a–c). Cell process geometry and packing were calibrated to match literature-reported ECS porosity targets across all sublayers (Figure 1d). All 64 flow simulations (8 independent geometries × 4 sublayers × 2 myelination conditions) reached a converged steady state, with both the mean-velocity trace and its iteration-to-iteration relative change falling below the pre-registered convergence thresholds (velocity residual < 10⁻⁶; relative change < 10⁻⁵ over 8 consecutive check intervals) in every condition (Supplementary Figure 1a,b; representative stratum radiatum trace shown for MY+ and MY−). The kinetic-energy spectrum of the converged velocity field followed a power law with exponent *α* = −1.31, markedly shallower than the *k*^−5/3^ Kolmogorov inertial-range scaling expected for developed turbulence, confirming that the simulated flow remained in the laminar, diffusion-dominated (Stokes) regime throughout (Supplementary Figure 1c), consistent with the imposed Re = 0.01 and the measured local Reynolds numbers of order 10⁻¹⁰–10⁻⁹ (see below).

Demyelination increased mean interstitial fluid velocity by 5.84-fold across all sublayers pooled (MY+: 0.052 ± 0.021 μm/s; MY−: 0.304 ± 0.076 μm/s; paired *t*-test, *t*(31) = 19.78, *p* < 0.0001; Figure 1g), an effect that was consistent in direction across every individual sublayer (SO: 8.11-fold, *p* < 0.0001; SP: 3.87-fold, *p* = 0.0002; SR: 6.64-fold, *p* < 0.0001; SLM: 6.06-fold, *p* < 0.0001), despite the four sublayers differing substantially in myelination ratio (Figure 1e) and baseline geometry (Figure 1c-d). Mean local kinetic energy showed a parallel 5.86-fold increase (*p* < 0.0001; Figure 1h), consistent with the velocity field becoming both faster and spatially more variable following loss of the myelin sheath (Figure 1f).

Notably, this velocity increase was not accompanied by a proportional increase in net directional transport: mean energy flux across the domain was reduced to 0.66-fold of the myelinated condition (*p* < 0.0001; Figure 1i), and mean viscous energy dissipation was reduced to 0.60-fold (*p* < 0.0001; Figure 1j). We interpret this apparent discrepancy as reflecting a qualitative reorganization of the flow field rather than a simple uniform acceleration: expansion of the ECS lumen lowers hydraulic resistance and permits locally faster, more disordered (vortical) flow, which raises instantaneous velocity and kinetic energy without correspondingly raising net directional flux or velocity-gradient-driven dissipation (Figure 1f). This interpretation is directly supported by voxel-wise flow-field statistics extracted from the same simulations (Supplementary Figure 1d–i). Representative per-pixel maps (stratum oriens) show that both the local Péclet number and vorticity magnitude are spatially concentrated into a sparse network of high-advection "flow channels" threading between packed cell processes, and that this channel network is visibly more extensive under demyelination (Supplementary Figure 1d,f,h). Quantitatively, demyelination significantly increased the mean local Péclet number by 5.13-fold pooled across sublayers (MY+: (1.56 ± 0.49) × 10⁻⁷; MY−: (8.00 ± 2.3) × 10⁻⁷; paired *t*(31) = 16.16, *p* < 0.0001; 3.5-to 7.2-fold within every individual sublayer, all *p* ≤ 0.0001; Supplementary Figure 1e). Because the Reynolds number in this model shares the same characteristic velocity and a fixed viscosity-to-diffusivity ratio (Schmidt number ≈ 1000), Re scales identically with Pe (Re = 10⁻³ × Pe) and yields the same fold-changes and significance levels (Supplementary Figure 1f,g).

Critically, mean local vorticity magnitude also increased significantly by 3.96-fold pooled (MY+: (3.75 ± 0.55) × 10⁻⁵; MY−: (1.48 ± 0.24) × 10⁻⁴ lattice units; paired *t*(31) = 23.60, *p* < 0.0001; 2.9-to 4.8-fold within every individual sublayer, all *p* < 0.0001; results confirmed by Wilcoxon signed-rank test in sublayers where the paired differences failed the Shapiro-Wilk normality pre-check; Supplementary Figure 1h,i), directly confirming that the expanded ECS lumen after demyelination supports more locally rotational flow rather than a uniform acceleration of directed transport. This distinction is addressed further in the Discussion and motivates the direct transport measurements presented in Figure 2. To confirm that these effects reflect the imposed demyelination-driven ECS widening rather than an idiosyncratic feature of a particular geometry realization, we examined how baseline geometric parameters modulate flow across all 32 simulations. Velocity reduction scaled approximately linearly with cell packing density (slope = 0.061 %/cell, *R*² = 0.468, *n* = 32) and mean ECS flow velocity followed an inverse power-law relationship with ECS volume fraction (*v* ∝ *φ*^−0.78^, *R*²(log) = 0.285, *n* = 32; Supplementary Figure 7a,b). Both relationships held in the direction predicted by Poiseuille-type ECS transport theory (denser packing and lower porosity both independently constrain interstitial flow), indicating that the demyelination effect reported above (Figure 1g) sits on this same geometry-velocity relationship rather than arising from a non-representative subset of geometries.

### 3.2 Demyelination Accelerates Molecular Transport through Expanded ECS

Demyelination increased the local ECS volume fraction α by 1.61-fold pooled across sublayers (MY+: 0.385 ± 0.066; MY−: 0.621 ± 0.114; paired *t*-test, *t*(31) = 24.6, *p* < 0.0001; Figure 2a), consistent with Figure 1. This geometric expansion was accompanied by a significant reduction in tortuosity *λ* (MY+: 1.691 ± 0.058; MY−: 1.463 ± 0.084; 0.87-fold, *p* < 0.0001; Figure 2b) and a corresponding 1.34-fold increase in the normalized effective diffusion coefficient *D*\*/*D*_free_ (MY+: 0.351 ± 0.022; MY−: 0.472 ± 0.050; *p* < 0.0001; Figure 2c), directly demonstrating accelerated molecular transport through the expanded ECS. This isotropic enhancement was consistent when diffusion was resolved separately along the two principal directions: demyelination reduced tortuosity along both *x* (*λ*_x_, 0.927-fold pooled; MY+: 2.056 ± 0.043, MY−: 1.906 ± 0.047; *p* < 0.0001) and *y* (*λy*, 0.838-fold pooled; MY+: 1.470 ± 0.062, MY−: 1.233 ± 0.090; *p* < 0.0001), and correspondingly increased *D_x_* (1.16-fold pooled, *p* < 0.0001) and *D_y_* (1.44-fold pooled, *p* < 0.0001), significant in every individual sublayer (Supplementary Figure 2b,c,e,f).

To determine whether this transport enhancement was fully explained by the change in ECS volume fraction, we examined the structure-transport scaling relationships across all sublayers and both myelination states pooled (Figure 2d-e). Both *λ* and *D*\*/*D*_free_ followed classical Archie’s-law-type power-law scaling with *α* (*λ* ∝ *α*^−0.27^; *D*\*/*D*_free_ ∝ *α*^0.54^), with MY+ and MY− data points from all four sublayers falling along common scaling curves. This indicates that demyelination does not introduce a qualitatively distinct transport regime, but rather moves the system along a single, geometry-determined structure-transport relationship by increasing *α*.

This enhanced transport was further supported by a 1.575-fold increase in the ECS constriction ratio *δ* = (P10/P90)² of the local width distribution (MY+: 0.0073 ± 0.0012; MY−: 0.0114 ± 0.0014; *p* < 0.0001; Figure 2f; higher *δ* indicates a more uniform, less constricted channel geometry) and a 2.18-fold increase in the geometric factor *M* (MY+: 0.137 ± 0.030; MY−: 0.298 ± 0.079; *p* < 0.0001; Figure 2g), both consistent with widened, less obstructed diffusion pathways following myelin loss. The directional anisotropy of effective diffusion (*D_x_*_/_*D_y_*) was significantly reduced following demyelination (MY+: 0.511 ± 0.025; MY−: 0.418 ± 0.042; 0.82-fold, *p* < 0.0001; Figure 2h), suggesting that loss of the highly oriented myelinated axon geometry renders the diffusion medium more isotropic.

Specific surface area (SSA) was correspondingly reduced by 0.47-fold (MY+: 8.42 ± 0.72 μm⁻¹; MY−: 3.93 ± 0.38 μm⁻¹; *p* < 0.0001; Figure 2i), reflecting the lower membrane surface density of the expanded extracellular compartment. Because the reduction in tortuosity was larger along *y* than along *x*, the tortuosity anisotropy ratio *λ_x_*/*λ_y_* increased by 1.27-fold following demyelination (MY+: 0.324 ± 0.021, MY−: 0.411 ± 0.040; *p* < 0.0001; Supplementary Figure 2d), the geometric counterpart of the reduced diffusion anisotropy *D_x_*/*D_y_* (Figure 2h). This was accompanied by systematic widening of the local ECS channel-width distribution: its coefficient of variation decreased by 0.83-fold pooled (MY+: 0.822 ± 0.022, MY−: 0.684 ± 0.018; *p* < 0.0001), while both P10 (1.73-fold; MY+: 39.3 ± 2.9 nm, MY−: 67.9 ± 7.8 nm; *p* < 0.0001) and P90 (1.37-fold; MY+: 464 ± 39 nm, MY−: 635 ± 39 nm; *p* < 0.0001) increased, all significant in every sublayer (Supplementary Figure 2g-i). The disproportionately larger increase at P10 than P90 shows that demyelination preferentially widens the narrowest bottleneck channels, consistent with the reduced CV and the increased constriction ratio *δ* reported above.

Finally, direct breakthrough-curve simulations using a sustained, constant-concentration glutamate source at one domain boundary (modelling how a locally elevated extracellular glutamate level would propagate across the ECS microenvironment, rather than a single transient synaptic release event) confirmed accelerated transport at the level of transport kinetics: both the time to 50% breakthrough (T50) and 90% breakthrough (T90) were reduced to 0.75-fold of the myelinated condition (T50: MY+ 2518 ± 176 ms vs. MY− 1888 ± 224 ms; T90: MY+ 72.5 ± 5.1 s vs. MY− 54.4 ± 6.5 s; both *p* < 0.0001; Figure 2j-k), indicating that glutamate reaches a given fraction of its equilibrium distribution substantially faster in the demyelinated ECS. This acceleration was already evident at early breakthrough: the time to 10% breakthrough (T10) was reduced to the same 0.75-fold ratio as T50 and T90 (MY+: 423 ± 30 ms, MY−: 317 ± 38 ms; *p* < 0.0001; Supplementary Figure 2a,j), reinforcing that all three breakthrough milestones are governed by a single underlying effective diffusion coefficient rather than kinetically distinct transport phases. Together, these structural and kinetic measures converge on a single conclusion: demyelination-driven ECS expansion accelerates molecular transport through a unified, geometry-determined mechanism rather than through independent or compensatory transport pathways.

### 3.3 Demyelination Impairs Synaptic Transmission through Neurotransmitter Dilution

Having established that demyelination expands the extracellular space and accelerates solute diffusion through it (Figs. 1–2), we next asked whether this structural change has a direct electrophysiological consequence at the single-synapse level. Recent in vivo evidence shows that synchronized neuronal firing itself generates large-amplitude ionic waves in the interstitial fluid, sufficient to drive measurable cerebrospinal fluid–interstitial fluid perfusion (Jiang-Xie et al., 2024), establishing that the ECS ionic/molecular environment is a dynamic, functionally consequential medium rather than a passive, fixed-composition background. This raised the possibility that a purely structural perturbation of the ECS — independent of any change in firing pattern — could similarly reshape conditions at the synapse. We therefore simulated single presynaptic–postsynaptic pairs across all four canonical connection types (P– P, P–I, I–P, I–I) with the number of intact myelin wraps swept continuously from 2 to 16 (pyramidal-presynaptic) or 2 to 8 (interneuron-presynaptic) layers, and examined how axonal conduction and postsynaptic potentials scale with myelin thickness.

Demyelination first degraded the fidelity of action-potential propagation along the axon in every connection type. Conduction velocity fell monotonically with fewer myelin wraps (P– P: 3.96 → 0.69 m/s from 16 to 2 layers, Spearman’s ρ = 0.99, p = 1.9 × 10⁻⁵⁴; P–I: 3.89 → 0.42 m/s, ρ = 0.99, p = 1.1 × 10⁻⁵⁷; I–P: 2.94 → 0 m/s from 8 to 2 layers, ρ = 0.98, p = 1.7 × 10⁻²¹; I–I: 2.87 → 0.13 m/s, ρ = 0.97, p = 7.8 × 10⁻²¹), and terminal axonal AP amplitude collapsed in parallel (P–P: −2.1 → −39.0 mV; P–I: −1.4 → −39.6 mV; I–P: −4.9 → −44.8 mV; I–I: −5.6 → −43.3 mV; all ρ ≈ 0.97, p < 10⁻¹⁸). Notably, interneuron-presynaptic axons (I–P) lost propagation outright at the thinnest myelin tested (2 layers, mean conduction velocity = 0 m/s), a more severe failure mode than seen in any pyramidal-presynaptic connection at the same nominal layer count, indicating that interneuron axons in this model are intrinsically more vulnerable to demyelination-induced conduction block. Consistent with this collapse in axonal fidelity, synaptic transmission success rate fell from 100% at 6 or more myelin layers (all four connection types) to markedly reduced levels at thinner myelin (4 layers: 38% for P–P, 25% for P–I, 0% for I–P, 38% for I–I; 2 layers: 0–12% across all types), with the shortfall attributable to a mixture of outright AP-propagation failure and synaptic release failure downstream of a successfully propagated but severely attenuated AP (Fig. 3g). Critically, restricting the analysis to trials with successful transmission — thereby isolating the amplitude effect from outright conduction failure — postsynaptic potentials remained substantially and monotonically smaller under demyelination in every connection type. Single-PSP amplitude, compared between the lowest layer count with 100% transmission success (layer 6) and the maximum layer tested, increased by 1.96-fold in P–P (2.37 → 4.65 mV, p = 1.8 × 10⁻²²), 1.94-fold in P–I (2.07 → 4.02 mV, p = 7.2 × 10⁻²²), 1.32-fold in I–P (3.70 → 4.89 mV, p = 1.7 × 10⁻⁴), and 1.24-fold in I–I (5.39 → 6.68 mV, p = 4.0 × 10⁻³) as myelin thickness increased (Fig. 3h,i). The consistently larger effect size in excitatory (P–P, P–I) than inhibitory (I–P, I–I) connections mirrors the corresponding difference in conduction-velocity fold-change (5.8–9.2-fold for EPSP-forming axons vs. 21.5-fold, but from a near-zero baseline, for I–I), suggesting that pyramidal presynaptic axons operate over a wider dynamic range of myelination-dependent conduction fidelity than interneuron axons in this model. Representative postsynaptic traces (Fig. 3f) illustrate this amplitude attenuation directly, without a qualitative change in waveform shape (further quantified in Supplementary Fig. 3).

**Figure 3.**
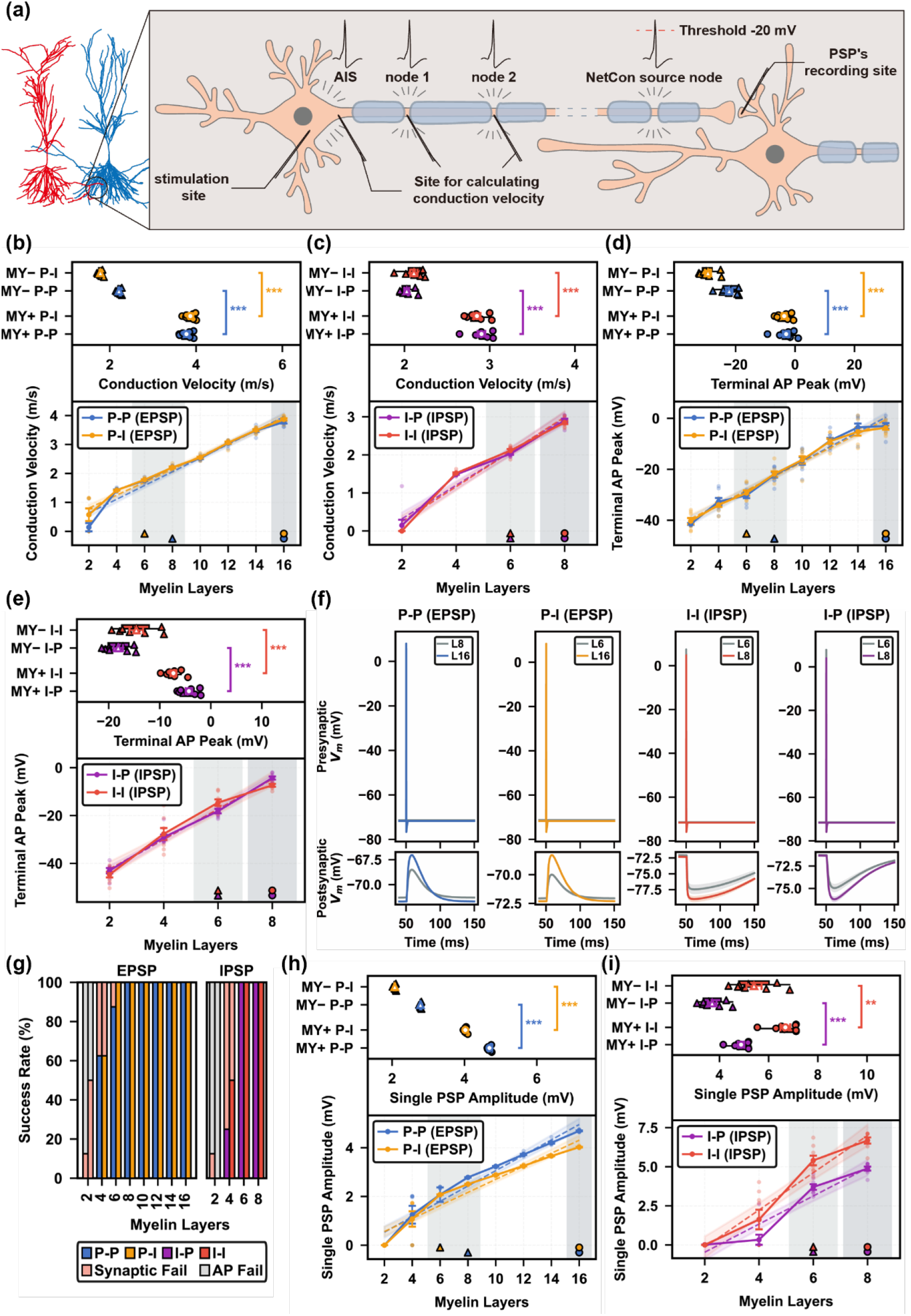
A continuous myelin-thickness gradient reveals graded, connection-type-specific degradation of both axonal conduction and synaptic transmission strength following demyelination. Single presynaptic–postsynaptic pair simulations across four connection types (pyramidal→pyramidal, P–P; pyramidal→interneuron, P–I; interneuron→pyramidal, I–P; interneuron→interneuron, I–I), with the number of intact myelin wraps swept from 2 to 16 layers for pyramidal-presynaptic connections (P–P, P–I) and 2 to 8 layers for interneuron-presynaptic connections (I–P, I–I), 8 independent geometric/ECS repeats per layer (incorporating the same per-repeat ECS jitter as Figs. 1–2 plus biological jitter in axon geometry, node spacing and synaptic kinetics). Top sub-panels in (b)–(e), (h), (i) compare the lowest layer count at which synaptic transmission was 100% successful across repeats (MY−, triangles; layer 6 for all four connection types) against the maximum layer count tested (MY+, circles; layer 16 for P–P/P–I, layer 8 for I–P/I–I), Welch’s *t*-test or Mann–Whitney *U* as appropriate (\*\*\**p* < 0.001, \*\**p* < 0.01, \**p* < 0.05, n.s. not significant); bottom sub-panels show the full dose–response trend across all layers (mean ± s.e.m., Spearman’s ρ and p reported in the text/legend). (a) Schematic of the simulated synapse: presynaptic axon wrapped by a variable number of myelin layers, the distal node from which the NetCon spike is triggered, and the postsynaptic recording site. (b) Conduction velocity for excitatory (EPSP-forming) connections, P–P and P–I. (c) Conduction velocity for inhibitory (IPSP-forming) connections, I–P and I–I. (d) Terminal axonal action-potential peak amplitude for EPSP connections. (e) Terminal axonal action-potential peak amplitude for IPSP connections. (f) Representative postsynaptic membrane-potential waveforms at low (MY−, layer 6) versus high (MY+, maximum layer) myelin thickness, one example per connection type. (g) Synaptic transmission success rate across all layers and connection types (stacked bars: successful transmission, synaptic failure, outright AP-propagation failure). (h) Single EPSP amplitude (successful-transmission subset only), P–P and P–I. (i) Single IPSP amplitude (successful-transmission subset only), I–P and I–I.

### 3.4 Progressive Demyelination Leads to Cumulative Neurotransmitter Dilution and PSP Decline

Single presynaptic–postsynaptic pair simulations (P–P, P–I, I–P, I–I; 8 independent repeats per connection type, same per-repeat ECS/biological jitter as Figs. 1–3) were driven through a continuous, progressively evolving myelin trajectory divided into five equal-duration phases — Initial, Degradation, Minimum, Recovery, Final — in which the myelin index decays exponentially from baseline (1.0) to a connection-type-specific floor and then recovers exponentially back to baseline, rather than the two discrete MY+/MY− states used in Fig. 3. The same ECS-mediated synaptic reversal-potential dilution mechanism as Fig. 3 (driven by the Section 1 extracellular volume-fraction data) operates continuously throughout, so that the degree of neurotransmitter/ion dilution at any instant tracks the instantaneous, cumulative degree of demyelination rather than switching between two fixed levels. Unless noted, bar/line panels show mean ± s.e.m. across 8 repeats with individual data points overlaid; paired t-test or Wilcoxon signed-rank test (matched by repeat), \*\*\**p* < 0.001, \*\**p* < 0.01, \**p* < 0.05, n.s. not significant.

(a) Schematic of the five-phase paradigm: the stimulation site, axon initial segment (AIS), the distal node-of-Ranvier NetCon source used for spike detection (threshold = −20 mV) and the postsynaptic PSP recording site, illustrated at each of the five phases. (b) Representative continuous extracellular LFP evoked by periodic PSP events (P–P, repeat 1), recorded at 16 linearly spaced virtual electrodes along each of the X, Y and Z axes centred on the neuron, showing that the ECS-dilution-driven PSP decline of panels (d)-(i) is also visible as a shrinking spatial LFP footprint — the population-level readout examined in Sections 5–6. (c) Mean ± s.d. (8 repeats) postsynaptic voltage waveforms time-locked to the single-pulse stimulus, for each connection type (P–P, P–I: EPSP; I–P, I–I: IPSP), overlaid across all five phases (colour-coded). (d) Single-pulse PSP amplitude across the five-phase cycle. Consistent with cumulative, progressive dilution rather than a step change, the decline was already significant by Degradation — before myelin integrity reached its minimum — in all four connection types (Initial vs Degradation, all p < 0.001), and PSP amplitude was statistically indistinguishable from zero at Minimum (one-sample test vs. 0, p ≥ 0.25), confirming complete loss of transmission at maximal dilution. (e) Single-pulse PSP energy across the five-phase cycle, mirroring (d). (f) Axonal AP conduction margin (mean terminal-node potential minus the −20 mV NetCon threshold; left axis, solid lines) and synaptic transmission success rate (right axis, dotted line, identical across connection types). The margin crosses zero exactly at Minimum, coincident with a drop from 100% to 0% transmission success: at this most severe point of demyelination, outright axonal conduction failure — not merely amplitude dilution — additionally silences the synapse. (g) Single-pulse PSP amplitude plotted against the instantaneous myelin index (pooled across repeats and phases, n = 40 per connection type), with OLS linear fit and 95% CI. All four connection types show a strong, near-identical positive correlation (Spearman’s ρ = 0.945–0.958, all p < 10⁻¹⁹; R² = 0.93–0.96), directly demonstrating that PSP amplitude tracks the cumulative, continuous degree of demyelination rather than exhibiting a discrete threshold — the central prediction of the neurotransmitter-dilution mechanism when tested under a continuously varying, rather than two-state, myelin trajectory. (h) Paired-pulse (2 ms inter-stimulus interval) maximum summated PSP amplitude across the five-phase cycle, showing the same graded collapse-and-recovery pattern as (d). (i) Paired-pulse total PSP energy across the five-phase cycle. Notably, the two interneuron-sourced connections (I–P, I–I) show a distinctly larger fold-reduction from Initial to Degradation (∼21-fold) than the two pyramidal-sourced connections (P–P, P–I; ∼3-fold), indicating that while the dilution mechanism operates uniformly across connection types, its downstream impact on temporally summated signalling energy is disproportionately larger for interneuron-sourced transmission (see Discussion).

Section 3 established, using a discrete two-state (MY+/MY−) design, that demyelination impairs single-synapse transmission through a common, quantitative mechanism: expansion of the ECS dilutes the local neurotransmitter/ion concentration available to the synapse, shifting its reversal potential away from resting potential and thereby reducing driving force (Fig. 3). Because that design necessarily compared only two fixed myelination states, it could not address whether this dilution effect is a genuinely graded, cumulative process that tracks demyelination continuously, or an artefact of comparing two arbitrarily chosen endpoints. We therefore tested the same neurotransmitter-dilution mechanism — using the identical ECS-mediated reversal-potential shift derived from the Section 1 extracellular volume-fraction data — under a fundamentally different, continuous modelling paradigm: a five-phase myelin trajectory (Initial, Degradation, Minimum, Recovery, Final) in which the myelin index of each of the four connection types (P–P, P–I, I–P, I–I) decayed exponentially from baseline to a connection-type-specific floor and then recovered exponentially back to baseline, rather than jumping between two fixed states (Fig. 4a). If neurotransmitter dilution is a genuinely cumulative process, PSP amplitude should decline progressively as demyelination deepens, and this decline should already be measurable before myelin integrity reaches its minimum — not appear only as a discontinuous jump at complete failure. This is exactly what we observed: single-pulse PSP amplitude fell significantly from Initial to Degradation in all four connection types (P–P: 5.38 → 4.22 mV; P–I: 4.03 → 3.09 mV; I–P: 4.67 → 2.67 mV; I–I: 6.67 → 4.12 mV; all paired comparisons *p* < 0.001), well before reaching the Minimum phase, where amplitude was statistically indistinguishable from zero (one-sample test vs. 0, *p* ≥ 0.25 for all connection types) and axonal AP conduction margin (terminal-node potential relative to the −20 mV spike-detection threshold) had crossed from strongly positive (20.7– 21.7 mV at Initial) to strongly negative (−11.2 to −4.1 mV), driving synaptic transmission success from 100% to 0% (Fig. 4f). Most directly, single-pulse PSP amplitude correlated strongly with the instantaneous myelin index when pooled across all repeats and phases (Spearman’s *ρ* = 0.945–0.958, all *p* < 10⁻¹⁹; *R*² = 0.93–0.96 for all four connection types; Fig. 4g) — a continuous, near-linear dose–response relationship rather than a step function, directly confirming that the magnitude of neurotransmitter dilution, and hence PSP attenuation, scales with the cumulative degree of demyelination at any given moment. This relationship was also symmetric: PSP amplitude and axonal conduction margin fully recovered by the Final phase in step with remyelination (Recovery-vs-Final paired comparisons, all *p* < 0.001), showing that the dilution effect is a reversible function of the instantaneous ECS/myelin state rather than a hysteretic or cumulative-damage process — consistent with a genuine dilution mechanism rather than an artefact specific to the discrete Section 3 design.

**Figure 4.**
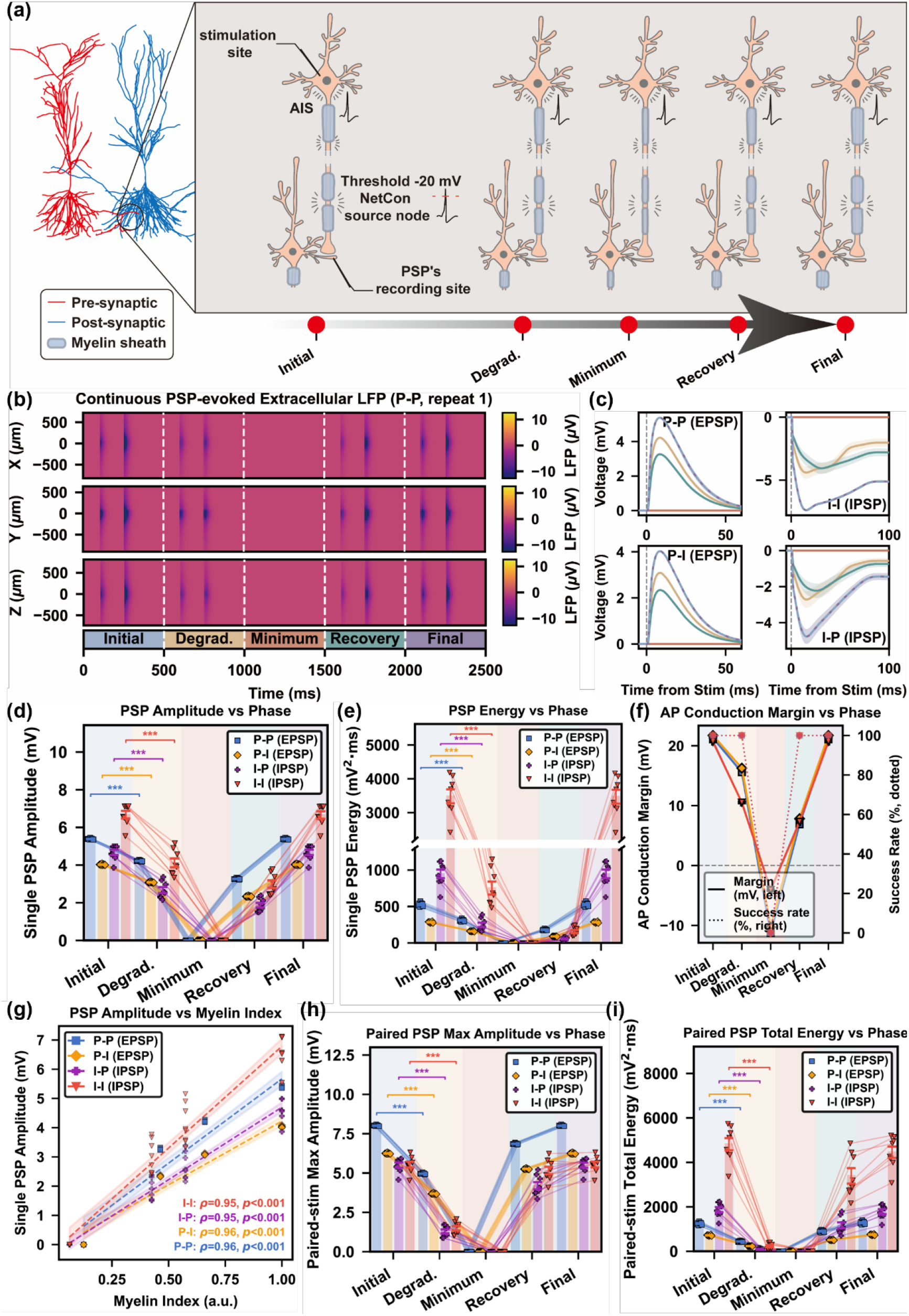
An independent, continuous demyelination–remyelination model converges on **the same neurotransmitter-dilution mechanism identified in** Fig. 3**: PSP amplitude and energy decline cumulatively and reversibly as demyelination progresses, tracking the instantaneous myelin index.**

This continuous framework further allowed us to test whether the dilution mechanism affects all connection types uniformly. It largely did: PSP energy fell 1.70–1.77-fold from Initial to Degradation in the two pyramidal-sourced connections (P–P, P–I) and paired-pulse total energy fell 2.82–3.06-fold in the same connections (Fig. 4e,i). However, the two interneuron-sourced connections (I–P, I–I) showed a substantially larger effect in the energy domain — 3.84–4.63-fold for single-pulse energy and a strikingly convergent ∼21-fold for paired-pulse total energy (I–P: 20.96-fold; I–I: 21.01-fold; Fig. 4i). Because I–P and I–I share an identical presynaptic interneuron axon/myelin trajectory, differing only in the postsynaptic target identity, the presynaptic drive reaching the synapse is essentially the same signal for both; this shared presynaptic pathway, propagated through an otherwise identical synaptic transduction mechanism, most parsimoniously explains why the two converge on nearly the same fold-change, and identifies temporally summated energy delivered by interneuron axons as a downstream readout that is disproportionately sensitive to the same underlying dilution mechanism, without requiring a distinct mechanism of its own.

Finally, as in Section 3, the intrinsic kinetic properties of the PSP waveform, including time-to-peak, rise time, and 90% spectral bandwidth (Supplementary Fig. 4a–c), did not change by an amount comparable to the amplitude and energy effects described above. The small residual changes in waveform shape, such as a shift toward a more leptokurtic IPSP under demyelination in I–P/I–I synapses (Supplementary Fig. 4i), covaried with the same amplitude/energy collapse and did not indicate an independent kinetic mechanism. This specificity check confirms that the dilution-driven amplitude/energy attenuation identified in Fig. 3 and reproduced here (Fig. 4d,e,h,i) is not confounded by a change in the synapse’s fixed conductance kinetics.

Collectively, these results indicate that two mechanistically independent modelling approaches converge on the same conclusion: a discrete two-state myelin-thickness gradient (Section 3) and a continuous demyelination–remyelination trajectory (Section 4).

Demyelination-driven expansion of the extracellular space dilutes the local neurotransmitter/ion environment and thereby weakens postsynaptic potentials in proportion to the instantaneous degree of demyelination, in a progressive and reversible manner. This cross-model convergence argues against a discretization artefact and supports neurotransmitter dilution as the mechanism linking ECS expansion (Figs. 1–2) to synaptic dysfunction. The extracellular LFP footprint anticipated in Fig. 4b then provides the basis for the population-level electrophysiological analyses in Sections 5–6.

### 3.5 Demyelination Reorganises Network-level Electrophysiology through Extracellular-Space-Mediated Neurotransmitter Dilution

Four network conditions were generated by crossing myelination with inflammatory cell composition. Myelination was set either to MY+ (16 wraps on pyramidal axons, 8 on interneuron axons) or to MY− (2 wraps on both classes), and composition either to the normal census (55 % neurons, 24 % astrocytes, 16 % oligodendrocytes, 5 % microglia) or to a chronic inflammatory census with the reactive-gliosis profile common to demyelinating and neuroinflammatory conditions (50 / 28 / 14 / 8 %). Eight networks were instantiated per condition and 4.8 s of steady-state activity was analysed in each. Figure 5a shows the layer-resolved anatomy and the resulting connection graph of one representative network, in which pyramidal somata occupy the pyramidal layer and their apical dendrites and the interneuron axons span the radiatum and lacunosum-moleculare.

**Figure 5.**
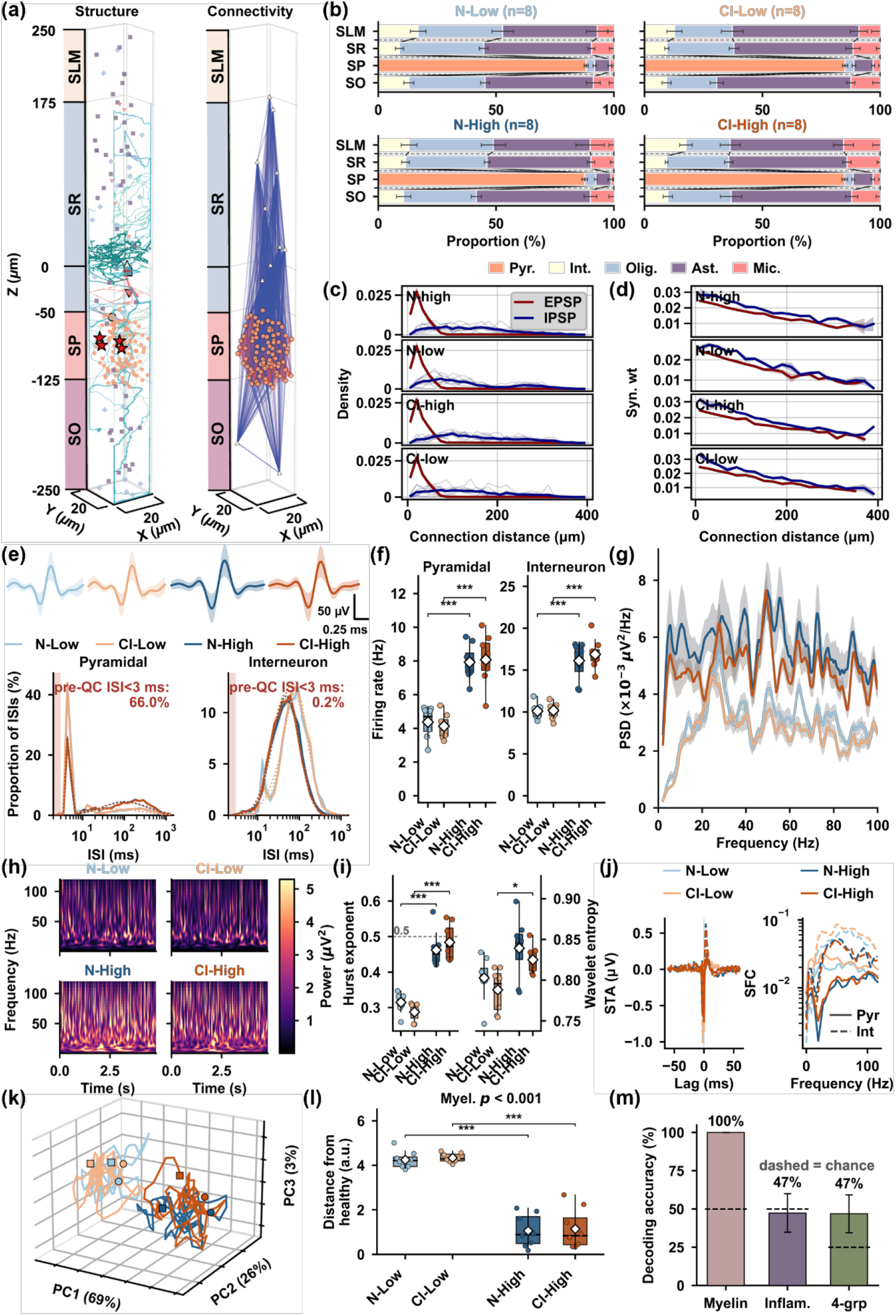
Myelination, rather than the glial census, determines CA1 network electrophysiology.

Four conditions were compared in a 2 × 2 design: myelination MY− (2 wraps on all axons) or MY+ (16 wraps pyramidal, 8 interneuron), crossed with the normal (N) or chronic inflammatory (CI) cell census; *n* = 8 independently instantiated networks per condition, 4.8 s of steady-state activity each. Throughout, light blue = N-Low (normal, MY−), light orange = CI-Low (inflamed, MY−), dark blue = N-High (normal, MY+), dark orange = CI-High (inflamed, MY+).

(a) Anatomy and connectivity of one representative network (N-High, run 01). Left, somata and reconstructed processes positioned within the four CA1 strata (SO, stratum oriens; SP, pyramidale; SR, radiatum; SLM, lacunosum-moleculare) in a 20 × 20 × 500 µm domain; red stars mark the tetrode recording sites. Right, the realised synaptic connection graph of the same network, glia excluded. (b) Cell-type composition of each stratum, mean ± SD over the eight networks of each condition. Pyr., pyramidal cells; Int., interneurons; Olig., oligodendrocytes; Ast., astrocytes; Mic., microglia. (c) Distribution of connection lengths for excitatory (EPSP, red) and inhibitory (IPSP, blue) connections. Thin lines, individual networks; thick lines, condition mean. One row per condition. (d) Mean synaptic weight as a function of connection length, same colours and layout as (c); shading is SEM across the eight networks. (e) Top, mean ± SD extracellular waveform of pyramidal spikes in each condition, sampled at 40 kHz over a fixed 1.5 ms window; scale bars 50 µV and 0.25 ms apply to all four. Bottom, interspike-interval distributions of pyramidal cells and interneurons on a logarithmic interval axis, pooled over cells and networks. The annotation gives the percentage of raw intervals shorter than the 3 ms quality-control criterion, averaged over the four conditions; these intervals are excluded from all subsequent analyses. (f) Mean firing rate of pyramidal cells (Pyr) and interneurons (Int). Boxes span the interquartile range with the median as a line and the mean as a white diamond; dots are the eight individual networks. (g) Power spectral density of the local field potential, multitaper estimate (NW = 6, 11 tapers) over the full 4.8 s window, smoothed over 13 frequency bins; line is the condition mean and shading the SEM across networks. The residual ripple is reproducible across networks and reflects the shared afferent drive realisation rather than estimation variance (see Methods). (h) Morlet wavelet spectrograms (ω₀ = 6), averaged across the eight networks of each condition and displayed on a common linear power scale. (i) Hurst exponent and wavelet entropy of the field potential. The dashed line at 0.5 marks the uncorrelated case; values below it indicate anti-persistence. Titles give the myelination main effect of the 2 × 2 ANOVA; inflammation and interaction were not significant for either descriptor (wavelet entropy p = 0.15 and 0.98; Hurst *p* = 0.84 and 0.088). Box conventions as in (f). (j) Spike-triggered average of the field potential (left) and spike-field coherence (right), computed separately for pyramidal cells (solid) and interneurons (dashed). (k) Trajectories of the instantaneous population state in the three leading principal components of a single analysis pooled over all conditions and networks (69, 26 and 3 % of variance). One representative network per condition; squares mark trajectory start and circles trajectory end. (l) Displacement of each network’s trajectory centroid from the healthy reference centroid (normal, MY+), in the units of the shared principal-component space; the reference condition itself is scored leave-one-network-out.

Boxes span the interquartile range with the median as the horizontal line and whiskers at 1.5 times the interquartile range, white diamonds are the mean with a vertical line spanning mean ± SD, and dots are the eight individual networks. Brackets give Holm-corrected Welch contrasts; the title gives the myelination main effect. The reference condition sits at 1.06 rather than 0 because the distance between two noisy centroids is positively biased; the offset is common to all four conditions. (m) Accuracy of a radial-basis support vector machine decoding myelination state (two classes), inflammatory state (two classes) and condition (four classes) from the seven population descriptors, under leave-one-network-out cross-validation. Bars are the mean over the eight folds, error bars SD, and dashed lines the corresponding chance level (50, 50 and 25 %). Myelination was decoded perfectly in every fold, hence no error bar. Statistics throughout: 2 × 2 factorial ANOVA for main effects and interaction, with four pre-specified contrasts tested by Welch’s *t*-test and Holm–Bonferroni correction within each measure. \**p* < 0.05, \*\**p* < 0.01, \*\*\**p* < 0.001.

The four conditions were anatomically matched apart from the two manipulated variables. Layer-resolved composition reproduced the intended census in every layer (Fig. 5b); across the four layer-by-neuron-class compartments that contain neurons, the only one in which the inflammatory manipulation was resolvable was the pyramidal fraction of the pyramidal layer (88.3 % and 87.6 % in normal networks against 85.1 % and 84.9 % in inflamed networks, *p* = 0.012), and no compartment differed by myelination (*p* ≥ 0.36). Wiring was likewise matched. Excitatory connections were short-range, with a density peaking near 20 µm and a mean length of 30.8–32.4 µm, whereas inhibitory connections reached four to five times further (mean 132–154 µm; Fig. 5c). Synaptic weight fell off with distance along an essentially identical gradient in all four conditions (Fig. 5d; slope −0.0609 to −0.0632 per µm, myelination *p* = 0.99, inflammation *p* = 0.43), and mean excitatory weight was unchanged (23.1 against 23.0 nS, *p* = 0.11). The one structural consequence of the inflammatory manipulation followed from its smaller neuronal fraction: excitatory synapse number fell from 8050 to 6380 at MY− and from 7274 to 6311 at MY+ (inflammation *p* = 3.3 × 10⁻⁵, myelination *p* = 0.12), a loss of 13–21 %, while inhibitory synapse number was not significantly affected (416 → 331 and 398 → 334, *p* = 0.15). The inflammatory manipulation was therefore not inert at the level of the wiring diagram, which is the relevant baseline against which its absence from the functional measures should be read.

Two properties of the simulated recordings constrain what can be measured and are reported before the functional results. First, the amplitude of the spike-locked field depended strongly on myelination: the ratio of centre to flank amplitude of the mean pyramidal waveform was 5.7 and 4.6 at MY− against 20.3 and 17.0 at MY+ (Supplementary Fig. 5h), so pyramidal spikes are far better resolved in the myelinated networks. For interneurons the ratio was indistinguishable from 1 in all four conditions, that is, the simulated field contains no interneuron spike-locked component, and every interneuron measure reported below therefore derives from spike times rather than from waveform detection. Second, raw pyramidal discharge was strongly burst-like: 66.0 % of detected pyramidal intervals fell below the 3 ms quality-control criterion, against 0.2 % for interneurons, and that fraction itself depended on myelination (77.3 % and 78.0 % at MY− against 54.5 % and 54.2 % at MY+, *p* = 5.5 × 10⁻⁹). After quality control the pyramidal interval distribution remained bimodal, with a burst mode near 5 ms and a second mode near 200 ms, whereas interneuron intervals were unimodal (Fig. 5e). The two modes are occupied very differently in the two myelination states: median pyramidal interval was 5.0 ms at MY− and 39.6 ms at MY+ (*p* = 1.0 × 10⁻⁴), while median interneuron interval shortened from 77.1 to 49.6 ms (*p* = 1.9 × 10⁻¹¹).

Both cell classes discharged roughly twice as fast in the myelinated networks (Fig. 5f).

Pyramidal rate rose from 4.38 ± 0.92 Hz (normal, MY−) and 4.14 ± 0.71 Hz (inflamed, MY−) to 7.93 ± 1.03 and 8.08 ± 1.49 Hz at MY+ (myelination *p* = 1.4 × 10⁻¹⁰, *η*² = 0.78; Hedges’ *g* = 3.43 in normal and 3.20 in inflamed networks, Holm-corrected *p* = 1.8 × 10⁻⁵ and 1.5 × 10⁻⁴). Interneuron rate rose from 10.12 ± 0.91 and 10.20 ± 0.94 Hz to 16.14 ± 2.27 and 16.90 ± 1.79 Hz (*p* = 5.6 × 10⁻¹², *η*² = 0.82; *g* = 3.29 and 4.43). Neither class showed a main effect of inflammation (*p* = 0.90 and 0.45) or an interaction (*p* = 0.61 and 0.55). The additional spikes were not simply added to the same discharge pattern: pyramidal firing became markedly more regular even as it accelerated, the coefficient of variation of the interspike interval falling from 2.17 ± 0.18 and 2.18 ± 0.12 to 1.21 ± 0.11 and 1.22 ± 0.14 (*p* = 1.0 × 10⁻¹⁷, *η*² = 0.93; Supplementary Fig. 5a), and the Fano factor of pyramidal spike counts falling from 2.99 ± 0.30 and 2.86 ± 0.29 to 0.95 ± 0.04 and 0.94 ± 0.04 (p = 2.6 × 10⁻²¹, η² = 0.96, |g| = 8.6–9.0).

Interneuron irregularity, by contrast, was unaffected by either factor (0.71, 0.69, 0.72 and 0.71; myelination *p* = 0.47; Supplementary Fig. 5b). Demyelination thus does not simply silence the network: it converts a moderately regular, faster discharge into a slower one dominated by bursts.

The absolute rates depend on the convention used to collapse repetitive discharge into a single counted event, but the separation between myelination states does not: the two low-myelination groups lie below the two high-myelination groups at every collapse window from 2 to 20 ms, with no crossing anywhere in that range (Supplementary Fig. 8g). Re-detecting spikes in the same simulations under a 3 ms rather than a 2 ms convention lowers pyramidal rate by 2.29 ± 0.74 Hz while leaving the transmission of spikes between cells untouched (+0.01 ± 0.06 Hz), so this sensitivity is a counting convention applied to genuine high-frequency discharge rather than a change in network behaviour (Supplementary Fig. 8a, b). Scaling the two recurrent weights that set the excitation-inhibition balance moved rates in the expected directions and in the same direction in every network without altering any conclusion: stronger inhibitory-to-pyramidal feedback reduced pyramidal rate by up to 1.02 Hz, and weakening pyramidal-to-interneuron drive reduced interneuron rate by up to 2.69 Hz (Supplementary Fig. 8d, e).

The local field potential was elevated at MY+ across the whole 2–100 Hz range (Fig. 5g), and the time–frequency representations show that in all four conditions the activity took the form of recurrent broadband transients on a low-frequency background, stronger and more frequent in the myelinated networks (Fig. 5h). Band power increased by factors of 1.4 to 8.7 (Supplementary Fig. 5c–f): delta from 1.21 ± 0.29 and 1.05 ± 0.39 to 10.5 ± 5.6 and 8.72 ± 3.23 (× 10⁻⁹ a.u.; p = 4.8 × 10⁻⁸), theta from 12.7 ± 2.6 and 10.9 ± 1.8 to 48.4 ± 24.2 and 38.7 ± 11.6 (p = 3.1 × 10⁻⁷), beta from 67.2 ± 7.4 and 62.6 ± 10.8 to 106 ± 33 and 89.2 ± 14.0 (p = 3.6 × 10⁻⁵), and gamma from 164 ± 23 and 141 ± 30 to 296 ± 93 and 259 ± 38 (*p* = 3.8 × 10⁻⁷). No band showed an effect of inflammation (*p* = 0.12 to 0.42) or an interaction (*p* = 0.37 to 0.71). The gain was not uniform across frequency, delta growing 8.7-fold while beta grew only 1.6-fold, so the spectrum of the myelinated network is not a scaled copy of the demyelinated one but is tilted towards low frequencies.

The same asymmetry between the two factors held for the nonlinear structure of the field potential. Of thirteen complexity descriptors, eleven were usable; the recurrence rate is fixed at 5 % by construction of the recurrence threshold and the median-binarised Lempel–Ziv complexity took only eight distinct values across the thirty-two runs, and both are shown with zero effect in the effect-size summary rather than suppressed (Supplementary Fig. 5i). Four descriptors survived Benjamini–Hochberg correction across the thirteen. The largest by a wide margin was the Hurst exponent (Fig. 5i, left), which rose from 0.315 ± 0.027 and 0.287 ± 0.024 at MY− to 0.462 ± 0.049 and 0.484 ± 0.050 at MY+ (*p* = 7.0 × 10⁻¹³, *q* = 9.1 × 10⁻¹², *η*² = 0.85, *g* = −4.19). All four values lie below 0.5, so the field potential is anti-persistent in every condition, but demyelination moves it substantially further from the uncorrelated case: successive fluctuations reverse more readily, consistent with the loss of the long-timescale structure seen in the spectra. Wavelet entropy rose from 0.803 ± 0.026 and 0.788 ± 0.026 to 0.839 ± 0.037 and 0.825 ± 0.017 (*p* = 7.5 × 10⁻⁴, *q* = 4.9 × 10⁻³, *g* = −1.30; Fig. 5i, right), that is, spectral energy is more evenly dispersed across scales in the myelinated network. The remaining two significant descriptors, permutation entropy (0.9988 → 0.9965, *p* = 3.1 × 10⁻³, *q* = 0.013) and Higuchi’s fractal dimension (2.004 → 1.990, *p* = 0.015, *q* = 0.049), both sit against their theoretical ceilings of 1 and 2 and change by less than 0.3 % and 0.7 % respectively, so they are reported only in the supplement. No descriptor showed a main effect of inflammation (smallest *p* = 0.079) or an interaction (smallest *p* = 0.086). These conclusions do not depend on the parametric model: Mann–Whitney *U* tests agreed with Welch’s *t*-test on every one of the fifty-two descriptor-by-contrast cells, with no reversal of significance.

Spike–field coupling separated the two cell classes (Fig. 5j). The pyramidal spike-triggered average deepened with myelination, its peak-to-trough amplitude rising from 0.41 ± 0.12 and 0.41 ± 0.08 µV to 0.62 ± 0.14 and 0.59 ± 0.14 µV (*p* = 1.2 × 10⁻⁴, *η*² = 0.42; Holm-corrected *p* = 0.025 and 0.030), while the interneuron spike-triggered average was unchanged (1.18–1.31 µV, *p* = 0.44). Spike-field coherence moved in the opposite direction and only for interneurons: pyramidal coherence was flat (*p* = 0.14), whereas mean interneuron coherence fell from 0.045 ± 0.011 and 0.057 ± 0.018 to 0.031 ± 0.004 and 0.029 ± 0.008 (*p* = 1.9 × 10⁻⁵, *η*² = 0.49; Holm-corrected *p* = 0.030 and 0.011). The same loosening of interneuron coupling appeared in gamma-band phase locking (0.457 ± 0.042 and 0.506 ± 0.056 at MY− against 0.400 ± 0.039 and 0.385 ± 0.048 at MY+; *p* = 9.5 × 10⁻⁶, *η*² = 0.51). Interneurons in the myelinated network therefore fire faster but are less tightly locked to the field, which is what would be expected if their discharge is driven increasingly by fast local input rather than entrained by the population rhythm.

The single place where the inflammatory manipulation left a functional trace was theta-band phase locking of pyramidal cells, and only in the demyelinated network. Pairwise phase consistency, which unlike the phase-locking value is not biased upwards by low spike counts, doubled from 0.0078 ± 0.0048 to 0.0158 ± 0.0066 when inflammation was added at MY− (inflammation main effect *p* = 0.012; Holm-corrected contrast *p* = 0.047), with no corresponding change at MY+ (0.0054 ± 0.0044 against 0.0071 ± 0.0042, *p* = 0.62) and no significant interaction (*p* = 0.090; Supplementary Fig. 5g, upper). The phase-locking value gave the same picture on the same data (0.088 ± 0.025 → 0.126 ± 0.025 at MY−, inflammation p = 0.0094, Holm-corrected contrast *p* = 0.028), and because firing rate did not differ between inflammatory states this agreement is not attributable to the spike-count bias of the phase-locking value. Myelination also acted on pyramidal consistency, and in fact more strongly than inflammation did (main effect *p* = 0.004), so this measure is not specific to the inflammatory factor; what is specific is that it is the one measure in which inflammation registered at all. Interneuron theta phase consistency responded to myelination only (0.0052 and 0.0059 at MY− against 0.0150 and 0.0155 at MY+, main effect *p* = 0.012, inflammation *p* = 0.87), although neither pre-specified contrast survived correction (both *p* = 0.30; Supplementary Fig. 5g, lower). Theta band power was itself unaffected by inflammation (*p* = 0.24), so this is a change in the timing of pyramidal spikes relative to an unchanged rhythm rather than a change in the rhythm.

Finally, the four conditions were compared as trajectories through a common population state space. Seven cell-count-independent descriptors of the instantaneous pyramidal rate distribution were pooled across all conditions and repeats and reduced by a single principal component analysis, whose first three components captured 69.4, 26.1 and 3.0 % of the variance. In that shared basis the two myelination states occupied disjoint regions, while normal and inflamed trajectories of the same myelination state were interleaved (Fig. 5k).

Quantified as the distance of each network’s trajectory centroid from the healthy reference centroid, the displacement was 4.25 ± 0.39 and 4.34 ± 0.21 at MY− against 1.06 ± 0.73 and 1.15 ± 0.90 at MY+ (mean ± SD; myelination *p* = 1.2 × 10⁻¹⁴, *η*² = 0.88, *g* = −5.18 and −4.64; inflammation *p* = 0.69, interaction *p* = 1.00; Fig. 5l). The reference condition does not sit at zero but at 1.06 because the Euclidean distance between two noisy centroids is positively biased; that offset is common to all four conditions and is differenced out by the factorial comparison, but the absolute values should not be read as true state-space distances. No geometric property of the trajectories separated the conditions. Cross-repeat reproducibility (*p* = 0.27) and the net displacement ratio (*p* = 0.45) were flat; trajectory speed and a path deviation measure each produced a myelination main effect (*p* = 0.010 and *p* = 0.012) but no pre-specified contrast survived correction for either (all *p* ≥ 0.23); mean Frenet curvature and mean absolute torsion showed no effect of either factor (all *p* ≥ 0.20); and three alignment-based summaries that yield a single value per group differed little between groups (residual Procrustes disparity 0.64 to 0.72, mean pairwise length-normalised dynamic time warping distance 0.070 to 0.077, and cross-repeat variance traces that were fully interleaved). The manifold signature of demyelination on these data is therefore a displacement of the population state rather than a change in the geometry of its path.

Decoding made the asymmetry quantitative. A support vector machine with a radial-basis kernel, cross-validated by holding out all time bins of one network at a time, identified myelination state from the instantaneous population descriptors with 100.0 ± 0.0 % accuracy in all eight folds against a chance level of 50 %, but identified inflammatory state at 47.4 ± 12.6 %, that is, at chance (Fig. 5m). Four-way condition identity was decoded at 46.9 ± 12.4 % against a chance level of 25 %; this is close to the 50 % that perfect myelination discrimination combined with chance-level inflammation discrimination would predict, so the above-chance four-way accuracy carries no information about inflammation. A linear kernel over identical folds gave the same result (100.0, 44.4 and 47.5 %).

Taken together, across roughly thirty measures spanning discharge rate and regularity, spectral power, nonlinear complexity, spike–field coupling and population geometry, the myelination manipulation produced large and mutually consistent effects, whereas the inflammatory manipulation produced none that survived correction except the doubling of pyramidal theta phase consistency in the already demyelinated network, and this despite having removed 13– 21 % of the excitatory synapses by construction. Where the two factors were tested for interaction, none was found (smallest *p* = 0.064 across all reported measures), so inflammation did not detectably modify the consequences of demyelination either. Three caveats bound this negative result. The inflammatory manipulation was a change of cellular census and of microglial phenotype; the intrinsic properties of neurons, the synaptic weights and the extracellular-space mechanism were identical in the two states, the last being driven by myelin wrap count rather than by composition. What is tested here is therefore whether the glial census alone reorganises network electrophysiology, and the answer bears on the signalling consequences of gliosis only to the extent that those act through the census. The eight networks of each condition share one realisation of the afferent Schaffer-collateral drive, so the reported p-values are anti-conservative for between-network comparisons (see Methods); this makes the significant myelination effects more secure than their nominal p-values only in sign, and makes the null inflammation effects, if anything, easier rather than harder to detect. And with n = 8 per condition the design has limited power for small effects: an effect of inflammation smaller than about g = 1.5 would not reliably be resolved here.

### 3.6 Progressive Demyelination Reversibly Disrupts Network Dynamics beyond a Threshold in Myelin Integrity

Sections 3 to 5 established the neurotransmitter-dilution mechanism at the single synapse and its consequences for a network held in one of two fixed myelination states. Neither design can say whether a network passes smoothly through intermediate states or whether function collapses only past a threshold, and neither can say whether the collapse reverses when myelin is restored. We therefore simulated a single continuous 25 s trajectory in which the myelin index decayed exponentially from 1.0 to 0.125, held at that floor, and then recovered exponentially back to 1.0, giving five consecutive 5000 ms phases (Initial, Degradation, Minimum, Recovery, Final). Cell composition was held at the chronic inflammatory census (50 % neurons, 28 % astrocytes, 14 % oligodendrocytes, 8 % microglia) for the entire run, so that the phase sequence isolates the myelination and ECS trajectory rather than confounding it with a change in cellular composition. Eight networks were run independently, and because all five phases are recorded from the same network, every comparison below is paired. An important consequence of the design should be stated at the outset: the mean myelin index was 0.996 in Initial, 0.125 in Minimum and 0.997 in Final, but 0.299 in Degradation and 0.711 in Recovery with the instantaneous value sweeping across almost the whole range within each of those two phases (0.13 to 0.99 and 0.13 to 1.00 respectively). Initial, Minimum and Final are therefore quasi-steady states and are the phases on which the quantitative claims below rest, whereas Degradation and Recovery are averages over a transition and are reported as such.

Raster and per-neuron rate displays of a representative network show the intended structure, namely dense discharge in Initial that thins over Degradation, reaches its sparsest point in Minimum, and refills over Recovery to a Final phase visually indistinguishable from Initial (Fig. 6a). Spike quality was verified before any functional analysis. Applying the uniform 3 ms counting convention reduced the raw pyramidal event count by 28.2 % to 38.9 % across the eight networks, affecting 22 to 44 of the 79 to 105 pyramidal cells per network, against only 0.1 % to 0.6 % of interneuron events, the asymmetry expected from a convention that acts on repetitive discharge within depolarised episodes at the distal axonal site and not on somatically detected interneuron spikes. After it, the proportion of intervals violating a 2 ms refractory period was 0.00 % in both cell classes, and the mean extracellular waveform retained the biphasic shape expected of a well-isolated extracellular spike (Fig. 6b). All spike-derived measures below use the merged trains.

**Figure 6.**
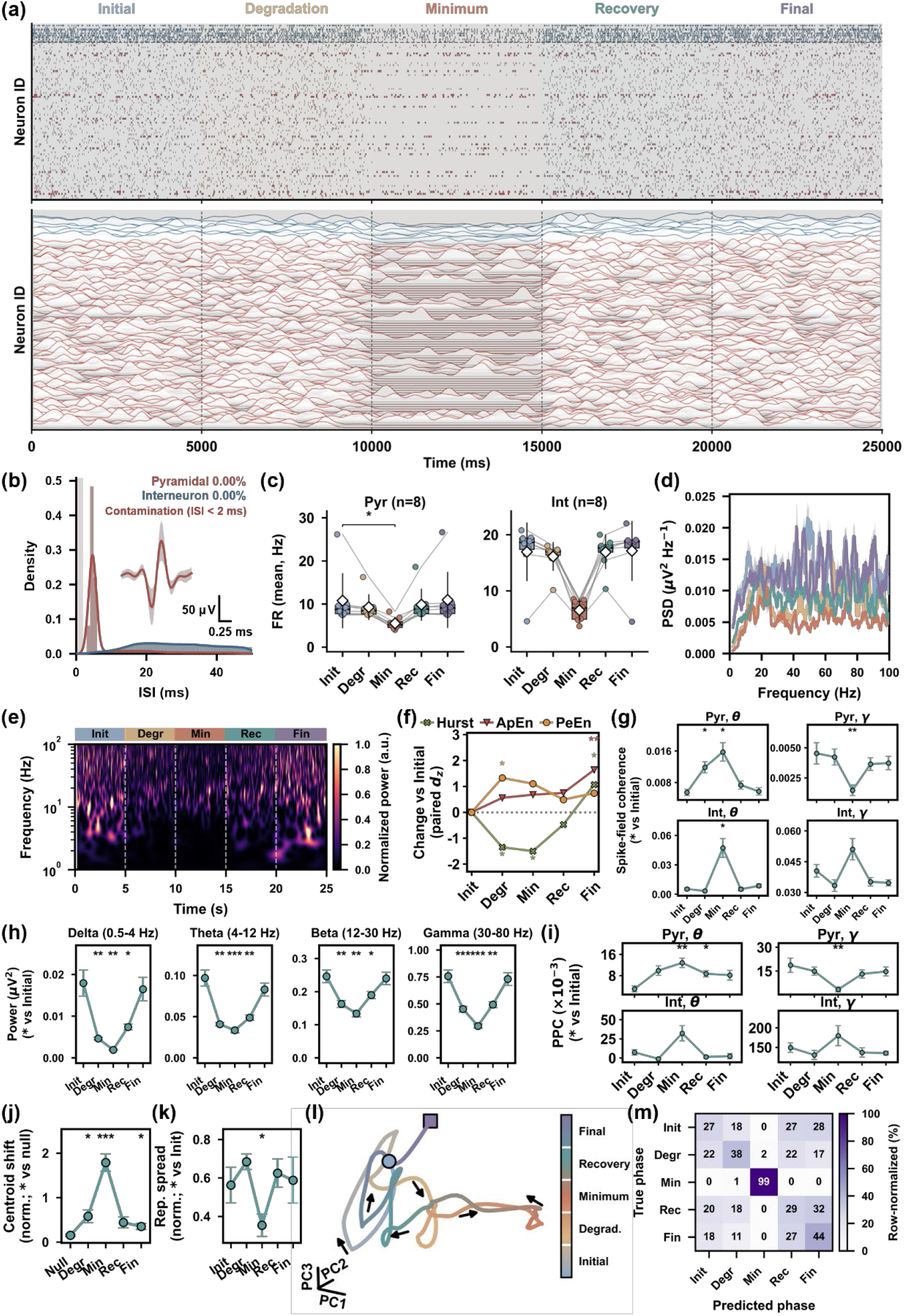
A continuous demyelination and remyelination trajectory reveals a threshold in myelin integrity beyond which network dynamics collapse and from which they largely, but not completely, recover.

Firing rate fell in both cell classes and then returned to baseline (Fig. 6c). Pyramidal rate went from 10.79 ± 2.23 Hz in Initial to 5.52 ± 0.50 Hz in Minimum and back to 10.99 ± 2.27 Hz in Final, a 49 % loss at the myelin floor that was consistent in direction in all eight networks (Wilcoxon with Holm correction, *p* = 0.031, *d_z_* = −1.01) and completely reversed by Final (*p* = 0.33, *d_z_* = +0.68). Interneuron rate fell further in relative terms, from 16.97 ± 1.83 Hz to 6.54 ± 0.65 Hz and back to 17.13 ± 1.88 Hz, a 61 % loss with a large paired effect size (*d_z_* = −2.25), but this comparison did not reach significance under the rank test (*p* = 0.0625) even though the parametric test on the same data gives *p* = 0.0015. The discrepancy is fully attributable to one network. Seven of the eight networks lost 10.4 to 14.0 Hz of interneuron rate at Minimum, whereas network 7 gained 0.27 Hz, and that network is atypical at baseline in exactly the way that would produce such an exception, with the highest pyramidal (26.2 Hz) and by far the lowest interneuron (4.58 Hz) Initial rate of the eight. With *n* = 8 the two-sided signed-rank statistic cannot fall below 0.0156 once a single pair is discordant, and Holm correction across four phases then places it above 0.05. We report the rank test as primary because the paired differences fail the Shapiro-Wilk pre-check in both cell classes (all *p* ≤ 0.005 for the Minimum contrast), and we note the parametric result and the effect size alongside it rather than resolving the discrepancy by test selection. The reduction is also visible without any pairing across phases: pooled over all 20 ms bins of all eight networks, instantaneous pyramidal rate correlated positively with the instantaneous myelin index (Spearman *ρ* = 0.270, *p* = 3.0 × 10⁻¹⁶⁶, *n* = 9968 bins; Supplementary Fig. 6b). Within-phase variability was close to Poisson at the single-cell level: in a representative network the median spike-count Fano factor was 0.98 for pyramidal cells and 0.81 for interneurons, whereas the Fano factor of the summed population count was 6.91 and 2.05 respectively, locating the excess variance in shared fluctuations rather than in individual cells.

A single 25 s simulation was divided into five consecutive 5000 ms phases: Initial (Init), Degradation (Degr), Minimum (Min), Recovery (Rec) and Final (Fin), during which the myelin index decayed exponentially from 1.0 to 0.125 and recovered back to 1.0 (mean index per phase 0.996, 0.299, 0.125, 0.711 and 0.997). Cell composition was held at the chronic inflammatory census throughout, so the trajectory isolates myelination and the associated extracellular-space changes. *n* = 8 independently instantiated networks, each contributing all five phases, so all comparisons are paired within network.

(a) Activity of one representative network over the full 25 s. Top, spike raster with interneurons at the top (blue) and pyramidal cells below (red); bottom, per-neuron smoothed firing-rate traces for the same network on the same time axis. Shaded bands mark the five phases. (b) Interspike-interval distributions of pyramidal cells (red) and interneurons (blue), pooled over cells and networks, with the mean ± SD extracellular waveform inset (scale bars 50 µV and 0.25 ms). The annotation gives the proportion of intervals shorter than the 2 ms refractory criterion after the 3 ms counting convention, which is 0.00 % in both classes. (c) Mean firing rate of pyramidal cells (Pyr) and interneurons (Int) by phase. Grey lines join the five phases of each network; boxes span the interquartile range with the median as a line and the mean as a white diamond. Statistics are Wilcoxon signed-rank tests against Initial with Holm–Bonferroni correction across the four comparisons, chosen as primary because the paired differences fail a Shapiro-Wilk pre-check. The interneuron loss at Minimum is large in effect size (*d_z_* = −2.25) but does not reach corrected significance (*p* = 0.0625) because one of the eight networks moves in the opposite direction; the corresponding paired *t*-test gives *p* = 0.0015. (d) Power spectral density of the local field potential by phase, multitaper estimate over each 5000 ms phase window, averaged across networks. (e) Wavelet spectrogram of one representative network over the full 25 s, power normalised to its maximum and displayed on a linear scale; the frequency axis is logarithmic and dashed lines mark the phase boundaries.

(f) Change from Initial in the three nonlinear descriptors that carry the effects, as the paired effect size *dz*: Hurst exponent, approximate entropy (ApEn) and permutation entropy (PeEn). Points are the mean over networks and error bars the SEM of the paired difference. Asterisks give Holm-corrected paired t-tests against Initial and are colour-matched to the descriptor; all starred cells except the Hurst exponent at Final also survive Benjamini-Hochberg correction across the thirteen descriptors computed. The remaining descriptors are shown individually in Supplementary Fig. 6d. (g) Spike-field coherence between spikes and the local field potential, in the theta (4 to 12 Hz) and gamma (30 to 80 Hz) bands, for pyramidal cells and interneurons separately. Spike counts were matched across phases within each network before computation, so that no phase difference can arise from the concurrent change in firing rate.

(h) Band power of the local field potential in the delta, theta, beta and gamma bands, as the integral of the multitaper spectral density over each band. (i) Pairwise phase consistency between spikes and field phase in the same two bands and the same two cell classes as (g), on the same spike-count-matched trains. Pairwise phase consistency is reported in preference to the phase-locking value because its expectation does not depend on spike count; the phase-locking value on the same data is shown in Supplementary Fig. 6c. (j) Displacement of each phase’s population-state centroid, normalised and compared against a split-half null baseline that measures the displacement expected by chance between two halves of the same phase. The reference category on the x-axis is therefore Null rather than Init. (k) Normalised spread of the population state within each phase, compared against Initial. (l) Population-state trajectory of one representative network in the three leading principal components, coloured by phase. The square marks the start and arrows the direction of travel. (m) Row-normalised confusion matrix of a linear support vector machine decoding phase identity from the seven population descriptors under leave-one-network-out cross-validation with nested hyperparameter tuning. The title gives overall accuracy and the chance level. Minimum is separated almost perfectly while the other four phases are confused predominantly with one another. Statistics throughout, except where stated in (c): paired *t*-test against Initial with Holm–Bonferroni correction across the four phase comparisons of each measure, normality of the paired differences verified by Shapiro-Wilk, and the Wilcoxon signed-rank test computed as a robustness check. \**p* < 0.05, \*\**p* < 0.01, \*\*\**p* < 0.001.

The relationship between rate and myelin integrity was strongly nonlinear. When phase-mean rate is plotted against phase-mean myelin index (Supplementary Fig. 6a), the descending and ascending limbs do not coincide: Degradation sits at a mean index of 0.30 with a rate of 9.29 ± 1.03 Hz, whereas Recovery sits at a 2.4-fold higher mean index of 0.71 with a rate only 6 % higher at 9.87 ± 1.29 Hz. Both transition phases therefore lie on a plateau, and almost the entire rate deficit is contributed by the interval between an index of about 0.3 and the floor of 0.125. This knee corresponds to between four and five intact wraps out of sixteen, which is precisely the range in which Section 3 found single-synapse transmission success to fall from 100 % at six wraps to 25 % to 38 % at four wraps, so the network-level threshold identified here and the synaptic threshold identified there are the same threshold. Because Degradation and Recovery average over different distributions of instantaneous myelin index, the non-coincidence of the two limbs in Supplementary Fig. 6a should be read as a description of the phase averages rather than as evidence of genuine path dependence; the direct test of path dependence is Initial against Final, which is null for both cell classes (*p* = 0.33 and 0.84).

Spectral power tracked the same trajectory and recovered completely. Power spectral density was highest in Initial and Final, lowest in Minimum, and intermediate in the two transition phases across the entire 2 to 100 Hz range (Fig. 6d), and the time-frequency representation shows that the activity took the form of recurrent broadband transients whose density and amplitude were reduced but not abolished at the myelin floor (Fig. 6e). Quantified by band (Fig. 6h), power at Minimum fell to 0.107 of its Initial value in delta (0.0179 ± 0.0031 to 0.00191 ± 0.00018 µV², *p* = 0.0047, *d_z_* = −1.86), 0.344 in theta (0.0969 ± 0.0098 to 0.0334 ± 0.0029 µV², *p* = 0.00054, *d_z_* = −2.66), 0.542 in beta (0.246 ± 0.019 to 0.134 ± 0.010 µV², *p* = 0.0035, *d_z_* = −1.95) and 0.390 in gamma (0.757 ± 0.058 to 0.295 ± 0.020 µV², *p* = 0.00046, *d_z_* = −2.73). The 9.4-fold loss in delta against a 1.8-fold loss in beta means that demyelination does not scale the spectrum uniformly but tilts it away from low frequencies, the mirror image of the low-frequency enrichment that accompanied myelination in the static design of Section 5. Every band was also significantly below Initial during Recovery (*p* = 0.018 to 0.0023), consistent with the plateau described above, and every band was statistically indistinguishable from Initial by Final (*p* = 0.115 to 0.659, all |*d_z_*| ≤ 0.64, recovering to 86 % to 98 % of the Initial value). Spectral power is therefore fully reversible on this timescale. These reductions are measured on a field potential that carries no background activity from outside the modelled volume, and they degrade if such background is imposed: adding synthetic broadband noise leaves delta and theta non-significant once the background matches the signal in power, while beta, gamma and total power hold to three times the signal and no band survives at ten times (Supplementary Fig. 8c). That is an adverse bound rather than an estimate of in vivo conditions, since the added background is uncorrelated with the signal at every frequency.

The nonlinear structure of the field potential behaved differently, and this is where the trajectory failed to close. Thirteen complexity descriptors were computed per phase; twelve are shown individually in Supplementary Fig. 6d and the three that carry the effects are summarised as paired effect sizes in Fig. 6f. Because Holm correction controls only the four phase comparisons within a descriptor and not the selection across thirteen descriptors, we additionally applied Benjamini-Hochberg correction across the thirteen within each phase, and report only cells that survive it. The Hurst exponent fell from 0.416 ± 0.018 in Initial to 0.358 ± 0.008 in Degradation (Holm *p* = 0.019, *q* = 0.046, *d_z_* = −1.35) and 0.354 ± 0.015 in Minimum (Holm *p* = 0.015, *q* = 0.049, *d_z_* = −1.50). All values lie below 0.5, so the field potential is anti-persistent throughout, exactly as in Section 5, and demyelination moves it further from the uncorrelated case. Permutation entropy rose from 0.9784 ± 0.0012 to 0.9845 ± 0.0009 in Degradation (Holm *p* = 0.029, *q* = 0.046, *d_z_* = +1.33). Both of these reversed by Final. What did not reverse was approximate entropy, which rose from 1.651 ± 0.012 in Initial to 1.683 ± 0.008 in Final (Holm *p* = 0.0096, *q* = 0.031, *d_z_* = +1.64) without ever reaching significance at Minimum. The Hurst exponent shows the same pattern in the opposite direction, overshooting its baseline in Final (0.447 ± 0.013, Holm *p* = 0.039), although this cell does not survive correction across the thirteen descriptors (*q* = 0.095) and is reported as suggestive only. No other descriptor survived Benjamini-Hochberg correction in any phase.

Restoring myelin therefore returns firing rate and spectral power to baseline while leaving the temporal complexity of the field potential measurably altered.

Spike-field coupling showed a frequency-specific dissociation that was the largest and most reproducible functional signature of the myelin floor. Because the Minimum phase contains the fewest spikes of the five, and because coherence and phase-locking estimators are biased upward at low spike counts, spike counts were matched across phases within each network before any coupling measure was computed (pyramidal counts subsampled to 2482 from a range of 2482 to 5014, interneuron counts to 297 from 299 to 782). Every coupling result below therefore compares equal numbers of spikes, and the phase with the fewest spikes is the reference to which the others were reduced. Pyramidal spike-field coherence in the theta band rose 2.9-fold from 0.0055 ± 0.0008 in Initial to 0.0158 ± 0.0024 in Minimum (*p* = 0.012, *d_z_* = +1.57), while pyramidal coherence in the gamma band fell 3.1-fold from 0.0045 ± 0.0009 to 0.0015 ± 0.0004 (*p* = 0.0036, *d_z_* = −1.95), both fully recovering by Final (*p* = 0.84 and 0.53; Fig. 6g). Interneuron theta coherence rose 8.9-fold at Minimum (0.0053 ± 0.0012 to 0.0472 ± 0.0097, *p* = 0.013, *d_z_* = +1.56) with no significant change in the gamma band (all *p* ≥ 0.087). Pairwise phase consistency, whose expectation does not depend on spike count at all, reproduced the pyramidal dissociation independently: theta consistency rose 4.3-fold from 0.0030 ± 0.0011 to 0.0128 ± 0.0018 at Minimum (*p* = 0.0019, *d_z_* = +2.17) and remained elevated through Recovery (*p* = 0.011, *d_z_* = +1.52), while gamma consistency fell 5.5-fold from 0.0187 ± 0.0045 to 0.0034 ± 0.0015 (*p* = 0.0083, *d_z_* = −1.68; Fig. 6i). The phase-locking value gave the same answer on the same spikes (theta 2.1-fold up, *p* = 0.0019; gamma 2.3-fold down, *p* = 0.0015; Supplementary Fig. 6c), and the two estimators diverged only for interneuron theta during Degradation, where the phase-locking value reports a decrease (*p* = 0.035) that pairwise phase consistency does not (*p* = 0.076); we treat the unbiased estimator as authoritative there. The consistent picture across three estimators is that at the myelin floor pyramidal spikes become more tightly locked to the surviving slow rhythm and less tightly locked to gamma, that is, the network does not simply couple less, it shifts the frequency at which it couples. This shift is not a consequence of a change in the rhythm’s amplitude, since theta power at Minimum was itself reduced 2.9-fold, so the same or fewer spikes are being timed more precisely against a weaker rhythm.

As a check on what inter-electrode coupling can be interpreted in this model, we also computed volume-conduction-immune connectivity between the four tetrode contacts of one network across the whole 25 s recording (Supplementary Fig. 6e). The weighted phase lag index was near its ceiling in the delta band for every one of the six contact pairs (0.942 to 1.000) but collapsed above it (theta 0.037 to 0.317, beta 0.007 to 0.071, gamma 0.016 to 0.121), and the unweighted phase lag index behaved likewise (delta 0.060 to 0.808, gamma 0.002 to 0.088). Only the delta band therefore carries a consistent non-zero-lag phase relationship across the tetrode; the high inter-contact coherence at higher frequencies reflects volume conduction from a shared source rather than lagged coupling between distinct generators. This is a within-tetrode measurement over tens of micrometres and does not license any statement about long-range connectivity, which is why all coupling claims in this section are made between spikes and the local field rather than between electrodes.

The population state traversed and returned along the manifold, but not exactly to its starting point. Instantaneous population activity was reduced to a shared three-dimensional basis, in which the trajectory leaves the Initial region, excurses furthest during Minimum, and loops back through Recovery towards Final (Fig. 6l). Quantified against a split-half null baseline that measures how far apart two halves of the same phase fall by chance, the displacement of the phase centroid was 11.9-fold above null at Minimum (1.786 ± 0.195 against 0.151 ± 0.024, *p* = 0.00053, *d_z_* = +2.67) and 3.8-fold above null already during Degradation (0.579 ± 0.141, *p* = 0.046, *d_z_* = +1.03). It remained 2.3-fold above null at Final (0.350 ± 0.070, *p* = 0.021, *d_z_* = +1.34; Fig. 6j), the same incomplete closure seen in approximate entropy. The state-space volume the network occupied contracted at the floor, the normalised spread falling to 0.63 of its Initial value (0.563 ± 0.092 to 0.354 ± 0.058, *p* = 0.032, *d_z_* = −1.29; Fig. 6k), so demyelination narrows the network’s dynamic repertoire in addition to displacing it.

Trajectory speed showed the same qualitative dip at Minimum (0.245 ± 0.041 to 0.178 ± 0.030) but did not reach significance (*p* = 0.129; Supplementary Fig. 6f), so we make no claim about the rate at which the state moves.

Decoding quantified how distinguishable the five phases actually are. A support vector machine trained on the seven population descriptors and cross-validated by holding out all time bins of one network at a time, with hyperparameters tuned in an inner cross-validation restricted to the training networks, identified phase at 47.5 ± 2.7 % against a chance level of 20 % (Fig. 6m and Supplementary Fig. 6g). A radial-basis kernel gave the same result (47.6 ± 3.8 %), as did an untuned linear kernel (48.0 ± 3.1 %). The descriptors were used directly, with no intervening embedding. Replacing them by the three leading principal components of each network taken separately dropped accuracy to 21.1 ± 10.4 %, that is, to chance, because each network’s principal-component basis is arbitrary and its coordinates are therefore not comparable across networks; bringing the trajectories into a common frame by Procrustes alignment first recovers 40.5 ± 11.5 %, still short of the descriptors themselves. The information thus resides in the population descriptors, and the low-dimensional display of Fig. 6l retains part of it only after alignment. The structure of the errors is more informative than the overall accuracy. Minimum was identified correctly in 99.0 % of its bins and was almost never predicted for any other phase (0.0 % to 1.8 % of their bins), whereas the remaining four phases were recovered at only 27.3 % (Initial), 37.8 % (Degradation), 29.0 % (Recovery) and 44.2 % (Final) and were confused predominantly with one another (Fig. 6m). The near-perfect separability of a single phase against near-chance discrimination among the other four is the decoding-domain statement of the threshold identified above: the population state is essentially binary with respect to whether myelin has fallen past the knee, and carries little information about where along the rest of the trajectory a given moment sits. The choice of embedding for that aligned low-dimensional view makes no difference either: a two-dimensional t-distributed stochastic neighbour embedding and the three-dimensional principal component analysis decode at 38.7 % to 40.8 % under either kernel, a spread of 2 % (Supplementary Fig. 8h).

Taken together, a continuously varying myelination trajectory produces three findings that the two-state designs of Sections 3 to 5 could not. First, the network-level response to demyelination is not graded but threshold-like, with almost the entire deficit in firing rate accumulating below a myelin index of about 0.3, and that threshold coincides with the four-to-six-wrap threshold for single-synapse transmission established in Section 3. Second, most of the disruption is reversible: firing rate and power in all four bands return to values statistically indistinguishable from baseline within one 5000 ms phase of myelin restoration. Third, reversal is nonetheless incomplete in a specific and consistent way, with approximate entropy elevated (*q* = 0.031) and the population centroid still displaced 2.3-fold above its null baseline (*p* = 0.021) at Final, while the Hurst exponent overshoots its own baseline (Holm *p* = 0.039, *q* = 0.095). Three limitations bound these conclusions. The eight networks share a single realisation of the afferent Schaffer-collateral drive, so between-network standard errors are underestimated and the reported p-values are anti-conservative; this matters least for the large paired effects (|*d_z_*| ≥ 1.9 for band power at Minimum) and most for the borderline ones, of which the Final-phase results are examples and should be treated as requiring independent confirmation. The five phases come from one continuous run, so they are not independent samples and the pairing is a design feature rather than a statistical convenience. And a 5000 ms recovery phase is short relative to biological remyelination, so the incomplete closure reported here bounds what recovers quickly rather than what recovers eventually.

## 4 Discussion

We set out to ask whether myelin acts on network function only as an insulator of axons, or also as a structural determinant of the space between cells, and whether the second route is strong enough to matter. Six linked models, from the lattice-Boltzmann flow field of a reconstructed neuropil geometry to a two hundred cell network of morphologically detailed neurons, give a consistent answer. Removing myelin enlarges the extracellular space, straightens the diffusive path through it, dilutes the neurotransmitter released into it, weakens and eventually abolishes synaptic transmission, and halves the discharge rate of the network, which loses power across the whole spectrum and moves to a displaced region of its state space. Restoring myelin reverses almost all of it. Figure 7 summarises the chain and the direction of each link.

**Figure 7.**
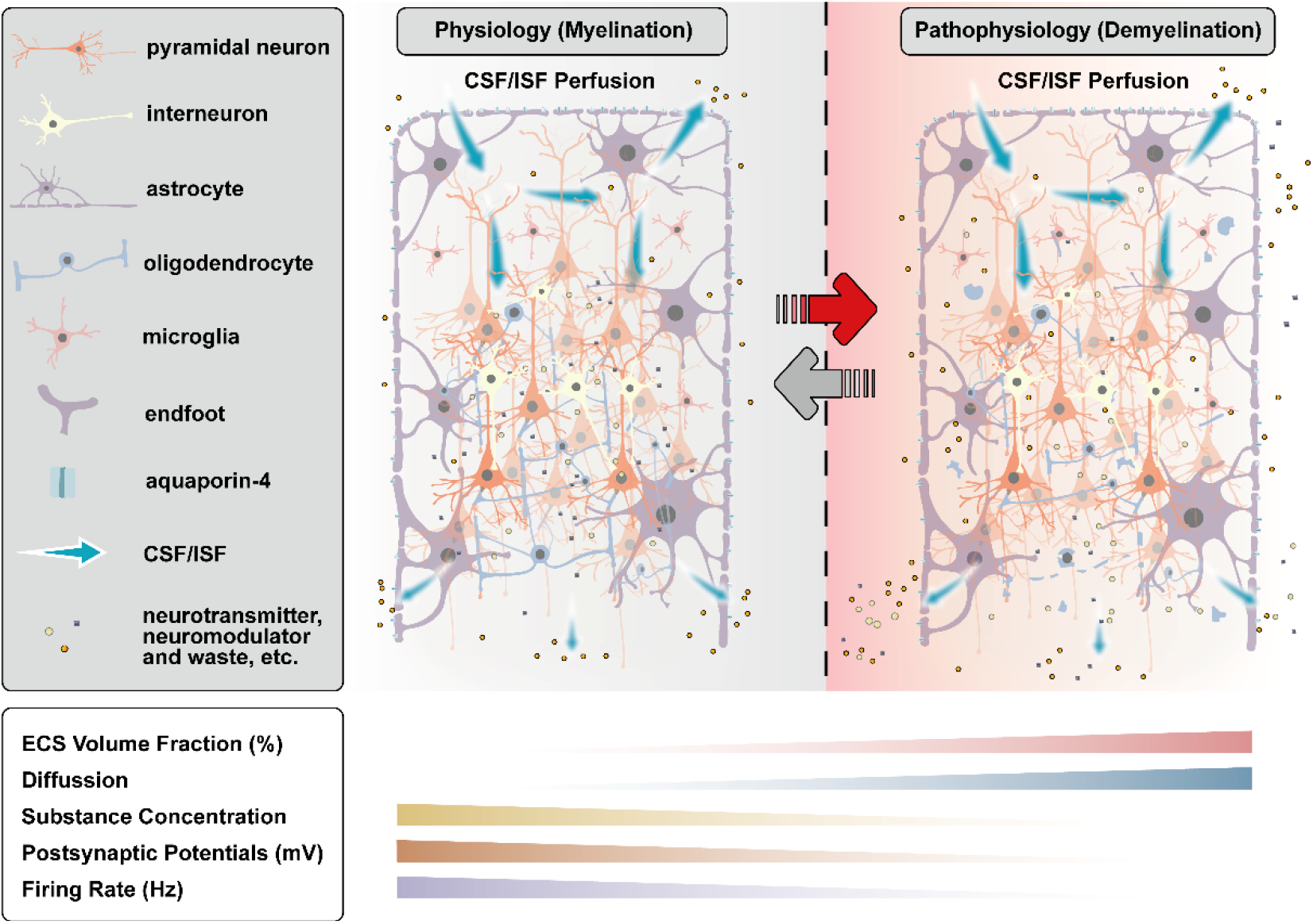
Myelin shapes the extracellular space, and through it both the transmission and the clearance functions that the space supports.

The first step is purely geometric and is the one most easily overlooked. Myelin is a space-occupying structure: sixteen wraps at 15 nm each add about a quarter of a micrometre to the radius of every pyramidal axon, and in a neuropil packed to a 38 to 64 nm minimum gap that volume has to come from somewhere. Stripping it away raised the extracellular volume fraction from roughly 28 to 43 per cent in the pyramidal layer and from roughly 42 to 69 per cent in stratum oriens (Fig. 2a), with tortuosity falling from about 1.7 to about 1.4 (Fig. 2b) and the normalised effective diffusion coefficient rising from about 0.35 to about 0.50 of its free-solution value (Fig. 2c). The two quantities follow the volume fraction with exponents of −0.27 and +0.54 respectively (Fig. 2d, e); these are not two independent observations, since the effective diffusion coefficient is defined as the inverse square of the tortuosity, and the agreement of the fitted exponents to within rounding is a consistency check on the extraction rather than a second result. Two caveats attach to the absolute numbers. The flow and transport fields were computed on two dimensional sections, so the volume fractions reported here are cross-sectional areas rather than three dimensional porosities and exceed the value of about 0.2 obtained by real-time iontophoresis in living tissue (Nicholson & Hrabětová, 2017; Nicholson & Phillips, 1981; Syková & Nicholson, 2008); and the tortuosity of a two dimensional section understates that of the connected pore network it samples. What should be compared with experiment is therefore the direction and the scaling, not the level. The direction deserves emphasis because it runs against the intuition that pathology narrows the extracellular space. In this model demyelination opens it, and the transport consequences are favourable: specific surface area halves, and the times for a solute front to cross the domain fall by roughly a quarter to a third (Fig. 2i to k).

The flow field behaves less simply than the diffusion field, and the distinction matters for how Figure 7 should be read. Mean flow speed in the extracellular space was several fold higher after demyelination in every layer (Fig. 1g), as the wider channels present less resistance, yet mean energy flux and mean viscous dissipation both fell (Fig. 1i, j). Faster fluid in a more open geometry does less work on the tissue, because the shear rates that dominate dissipation are set by the gradient across the gap rather than by the bulk speed. Perfusion and transport therefore move in the same direction here while the energetics move in the other, and any statement that demyelination "improves" extracellular flow needs to specify which of the three is meant.

The second step converts the geometric change into a synaptic one. In the present implementation the local concentration of released transmitter scales inversely with the extracellular volume available to it, and that dilution enters the synaptic model as a shift of the reversal potential, so that a given conductance change produces a smaller driving force. The consequence at a single synapse is steeply nonlinear rather than graded. Postsynaptic potential amplitude grew monotonically with wrap count, from complete failure at two wraps to 4.69 mV at sixteen for the pyramidal to pyramidal connection, and transmission success went from zero at two wraps, through 25 to 63 per cent at four, to complete from six wraps upwards (Supplementary Fig. 7d, g). The continuous dynamic model of Section 4, which shares the ionic and synaptic mechanisms but drives myelin as a continuous index rather than as a discrete wrap count, reproduced both the direction and the location of that threshold (Supplementary Fig. 7e, h). Two independently constructed representations of the same physical variable agreeing on where transmission fails is stronger evidence than either alone, and it is what licenses treating the wrap count and the myelin index as interchangeable descriptions in the network sections. The magnitude of the dilution effect is also consistent with what is known about the extrasynaptic fate of glutamate, where the concentration reached at a given distance from the release site depends directly on the volume fraction and tortuosity of the intervening space (Rusakov & Kullmann, 1998).

At the network level the same threshold reappears. In the static factorial design of Section 5, where myelination was fixed at either two or sixteen wraps, demyelination roughly halved the discharge of both cell classes (pyramidal 4.4 against 7.9 Hz, interneuron 10.2 against 16.1 Hz), and did so while making pyramidal firing markedly more irregular, with the coefficient of variation of the interspike interval rising from 1.21 to 2.17 and the Fano factor from 0.95 to 2.99. Demyelination does not simply turn the network down; it converts a moderately regular discharge into a slower one dominated by bursts. Band power fell by factors of 1.4 to 8.7, and the loss was largest at low frequencies, so the spectrum tilts rather than scaling. In the continuous trajectory of Section 6 the same network passed through the intermediate states, and the rate deficit was not distributed evenly along the way: phase mean rate against phase mean myelin index lies on a plateau down to an index of about 0.3, with almost the entire deficit accumulating between 0.3 and the floor of 0.125. That knee corresponds to four to five intact wraps out of sixteen, which is the range in which single synapse transmission collapsed in Section 3. The network threshold and the synaptic threshold are the same threshold, observed twice at two levels of description.

Schematic summary of the mechanism established in Figures 1 to 6. The left panel shows the myelinated state (MY+, 16 wraps on pyramidal axons and 8 on interneuron axons) and the right panel the demyelinated state (MY−, 2 wraps on all axons); the arrow between them gives the direction of the process, from physiology to pathophysiology, and denotes the continuous loss of myelin simulated as a trajectory in Section 6 rather than a comparison between two unrelated tissues. Each panel shows one field of hippocampal CA1 neuropil containing pyramidal neurons, interneurons, astrocytes with their perivascular endfeet and aquaporin-4 water channels, oligodendrocytes and microglia, in the cell-type proportions used in the simulations; the census is identical on the two sides, since only myelination differs between the states being contrasted. Teal arrows mark the movement of cerebrospinal and interstitial fluid through the extracellular space, and the small circles and squares mark the solutes it carries, namely the neurotransmitter and neuromodulator released at synapses together with metabolic waste. Loss of myelin returns the volume formerly occupied by the sheath to the extracellular space, drawn on the right as wider clearances between cellular processes.

The five wedges below give the direction in which each quantity moves along that progression, their width indicating relative magnitude rather than any numerical scale. Extracellular volume fraction and diffusion both increase (Fig. 2a, c): the space opens and the diffusive path through it straightens. The concentration reached by a given quantity of released transmitter therefore falls, being diluted into a larger volume, and with it the postsynaptic potential (Supplementary Fig. 7g, h) and the discharge rate of the network (Figs. 5f and 6c). The third wedge refers to the synaptic transmitter specifically: the same dilution that weakens transmission accelerates the clearance of waste, so the two classes of solute drawn in the panels above move in opposite directions in terms of what the change means functionally. The teal arrows are not scaled to the simulated flow, which rises in speed while falling in viscous dissipation on demyelination (Fig. 1g, j). No element of this figure is drawn to scale and none is simulation output.

The trajectory also shows that most of the disruption is reversible. Firing rate returned to within 2 per cent of baseline and every frequency band to between 86 and 98 per cent of it, none differing significantly from Initial. The exception is instructive: approximate entropy of the field potential was higher at the end of the trajectory than at its start (1.651 to 1.683, *q* = 0.031) although it was never significantly altered at the myelin floor itself, and the Hurst exponent overshot its baseline in the same direction, though that cell does not survive correction across the thirteen descriptors and is reported as suggestive only. One descriptor out of thirteen is a slender basis for a claim about hysteresis, and we do not make one. What can be said is that the measures that recover are those that summarise how much the network fires and at what frequencies, while the measure that does not is one that summarises the temporal predictability of the field, and that the two need not recover together.

The population geometry gives the same picture from a different angle. A three dimensional projection of the population state separates the myelination conditions cleanly in Section 5, with a support vector machine identifying myelination at 100 per cent accuracy under leave one network out validation, and the separation is a displacement of the state rather than a change in the shape of the trajectory: no descriptor of curvature, torsion or path deviation survived correction, whereas the distance of each condition from the healthy state did. In Section 6 the five phases were decodable at 47.5 per cent against a chance level of 20 per cent, with the Minimum phase almost perfectly identified and the two transition phases confusable with their neighbours, which is the error structure expected of a trajectory sampled at five points rather than of five distinct states. Reading these as a representational displacement rather than as a change in dynamical repertoire follows the logic of representational geometry, in which the relevant quantity is the arrangement of states rather than the coordinates of any one of them (Cunningham & Yu, 2014; Kriegeskorte et al., 2008; Kriegeskorte & Wei, 2021).

Two features of the design bound what these results can claim. The inflammatory arm of the Section 5 factorial varied the cellular census and the microglial phenotype, and it removed 13 to 21 per cent of the excitatory synapses by construction, yet it left every functional measure unchanged except pyramidal theta phase consistency. That null is informative about the census specifically, and it is not evidence that neuroinflammation is functionally silent, because the intrinsic properties of neurons, the synaptic weights and the extracellular mechanism itself were identical in the two states: the signalling consequences of gliosis are simply not represented in the model. The eight networks of each condition also share one realisation of the afferent Schaffer collateral drive, so they are not fully independent and the reported p values are anti-conservative for comparisons made across networks. This makes the significant myelination effects more secure in sign than their nominal values suggest, and makes the null inflammation effects, if anything, easier rather than harder to detect.

Other limitations follow from the modelling choices. Beyond the dimensionality of the flow domain noted above, the geometries are static: cells neither swell nor move, and no regulation of extracellular volume by astrocytic transport is represented, so the model describes the space that myelin vacates and not the space that glia would subsequently reclaim. The dilution mechanism enters the synaptic model as a shift of the reversal potential rather than as an explicit diffusion of transmitter in the reconstructed pore space, which captures the direction and the threshold but not the time course of the concentration transient. In the network simulations only the neurons within the core volume surrounding the electrodes carried their full reconstructed morphology, which shifts the absolute firing rate without entering any of the reported contrasts (Supplementary Fig. 8f). And the model contains no feedback from activity to myelination, so it can describe what a given degree of myelin loss does to the network but not how the network would respond to it over the timescale on which oligodendrocytes remodel.

Within those bounds the result is a specific and testable one. Myelin loss acts on the network through the space it vacates as well as through the axons it uncovers, and the two routes have opposite signs for the two functions that the extracellular space serves: clearance improves while transmission fails. The threshold at which the second effect appears is sharp, sits near a quarter of the normal wrap count, and is the same whether it is measured at one synapse or in the discharge of a whole network. If it holds in tissue, it implies that partial remyelination need not be proportionally partial in its benefit, since recovering the first few wraps past the threshold would restore transmission that additional wraps beyond it would not further improve, and it identifies the extracellular volume fraction, which is measurable in vivo, as the quantity that reports on the state of the network rather than on the state of the sheath alone.

## 5 Highlights

- Coupled lattice Boltzmann and NEURON models link ECS geometry to CA1 dynamics
- Demyelination widens extracellular space and raises effective diffusion
- Widened ECS dilutes transmitter, cutting PSP amplitude and network firing
- Myelination, not glial census, determines CA1 network electrophysiology
- Demyelination cuts firing and band power; remyelination restores both

## 6 Declaration

### 6.1 Ethics approval and consent to participate

Not applicable.

### 6.2 Consent for publication

Not applicable.

### 6.3 Availability of data and materials

The code and the derived data supporting the findings of this study are available from the corresponding author on reasonable request. The neuronal morphological reconstructions used are publicly available from the NeuroMorpho.Org archive.

### 6.6 Authors’ contributions

Xin Jia: Conceptualisation, Methodology, Software, Formal analysis, Investigation, Data curation, Visualisation, Writing – original draft. Cuidie Zeng: Validation, Writing – review & editing. Jing Bi, Lingchen Liu, Tengao Gao and Zhiming Zeng: Writing – review & editing. Lei Liang, Chenxiang Qi and FuYuan Lin: Writing – review & editing. Fangxiao Cheng: Conceptualisation, Supervision, Resources, Funding acquisition, Project administration, Writing – review & editing. All authors read and approved the final manuscript.

### 6.7 Acknowledgements

Not applicable.

### 6.8 Declaration of generative AI and AI-assisted technologies in the manuscript preparation process

During the preparation of this work, the author(s) used [NAME OF TOOL/SERVICE] for [REASON]. The author(s) reviewed and edited the output as needed and take full responsibility for the content of the published article.

---

**Supplementary Figure 1.**
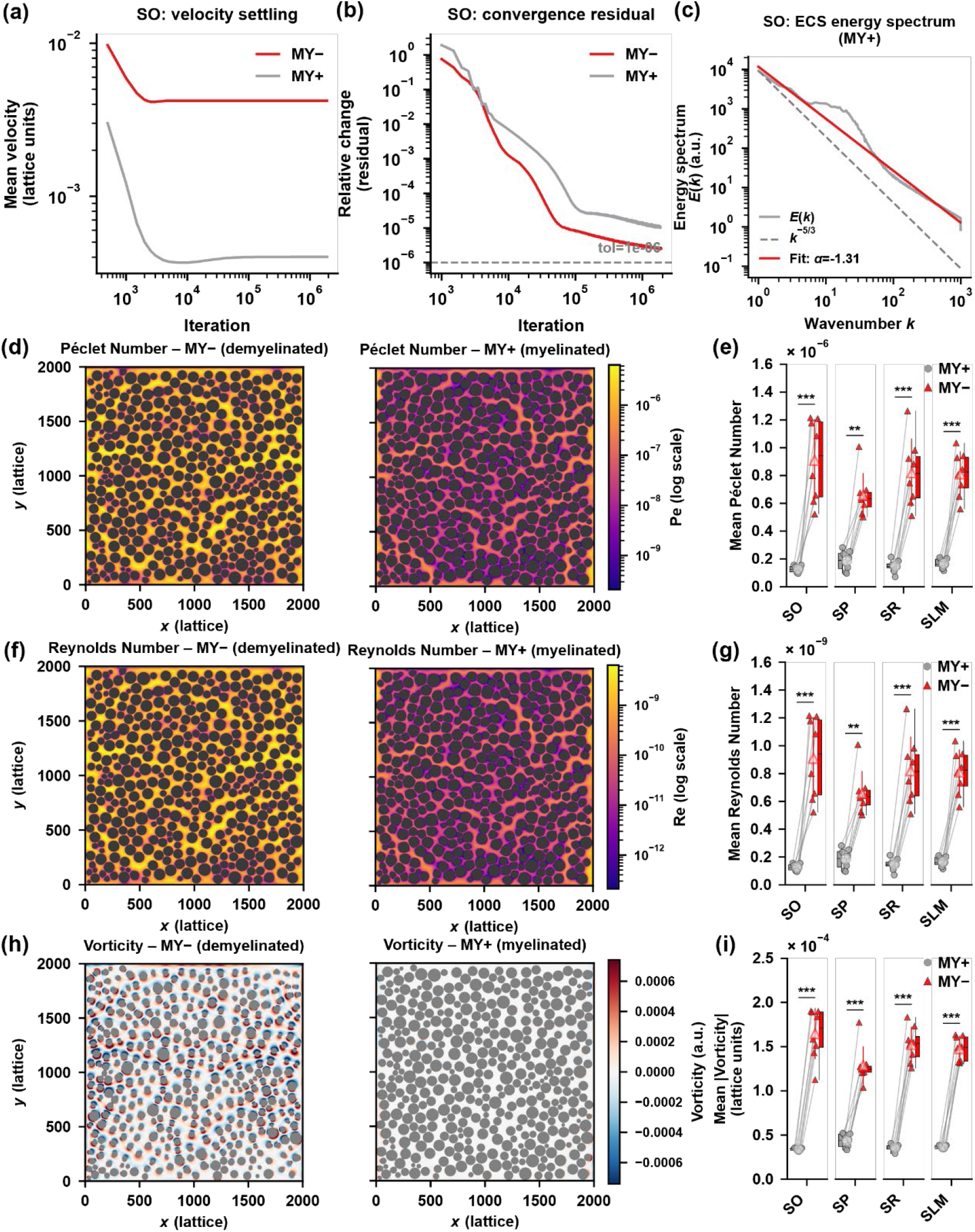
Numerical convergence, flow-field statistics, and parameter sensitivity of the LBM extracellular-space flow simulations (Section 1). (a) Mean flow velocity vs. iteration (representative: stratum oriens, MY+ and MY−), showing convergence to a stable plateau. (b) Relative iteration-to-iteration change in mean velocity on a log scale, both conditions falling below the 10⁻⁵ convergence threshold (dashed line). (c) Kinetic-energy spectrum E(k) of the converged velocity field (log-log), with a fitted power-law exponent (α = −1.31) markedly shallower than the k^−5/3 scaling expected for a 2D turbulent inverse cascade (dashed reference), confirming the flow remains in the laminar, diffusion-dominated regime. (d) Representative per-pixel Péclet number map (stratum oriens, log color scale), illustrating a sparse network of high-Pe advective "flow channels" between packed cell processes. (e) Mean local Péclet number across the four CA1 sublayers under MY+ and MY− (paired comparison, n = 8 geometries/sublayer; **/***p < 0.01/0.001). (f) Representative per-pixel Reynolds number map, same simulation as (d); because Re and Pe share the same characteristic velocity and a fixed viscosity-to-diffusivity ratio in this model, Re = 10⁻³ × Pe pixel-wise. (g) Mean local Reynolds number across sublayers, MY+ vs MY− (statistically identical to (e) up to the constant scale factor). (h) Representative per-pixel vorticity magnitude map, same simulation as (d). (i) Mean local vorticity magnitude across sublayers, MY+ vs MY−.

**Supplementary Figure 2.**
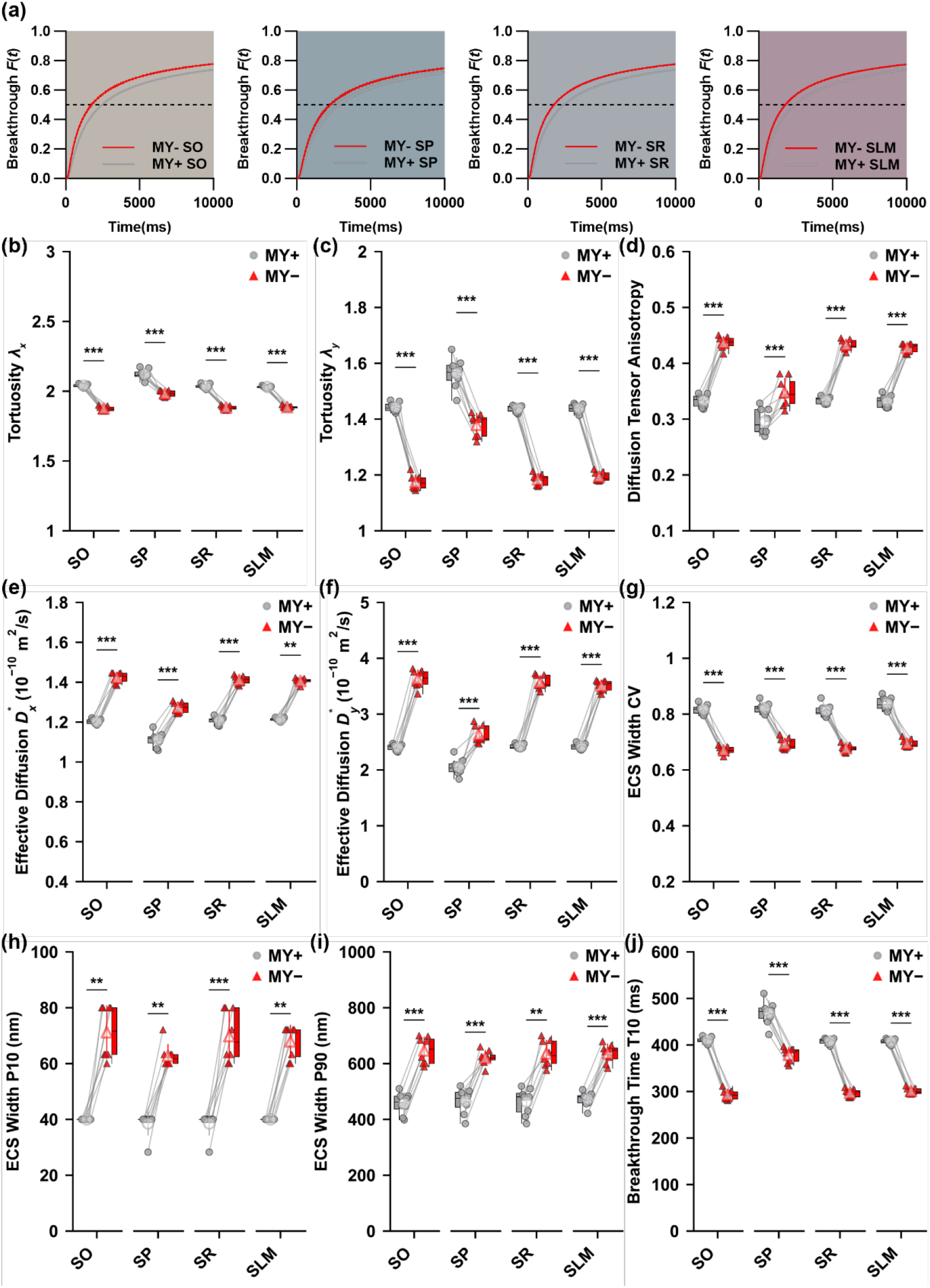
Directional and structural decomposition of the ECS diffusion geometry underlying Figure 2, using the same eight paired MY+/MY− geometries per sublayer. Unless otherwise noted, all panels compare the four CA1 sublayers (SO, SP, SR, SLM) under myelinated (MY+, grey circle) and demyelinated (MY−, red triangle) conditions (*n* = 8 independent geometries per sublayer, paired t-test; \*\*\**p* < 0.001). (a) Glutamate breakthrough curves (concentration fraction of the sustained source’s steady-state value vs. time) for all *n* = 8 repeats per sublayer (SO/SP/SR/SLM), MY+ (grey) vs. MY− (red), using a constant-concentration boundary condition at one domain edge; the consistent leftward (faster) shift of all eight MY− curves relative to their paired MY+ curves within each sublayer illustrates the transport acceleration underlying the T10/T50/T90 statistics in (j) and Figure 2j-k. (b) Tortuosity component *λ_x_* (diffusion direction parallel to the dominant axon orientation). (c) Tortuosity component *λ_y_* (perpendicular direction). (d) Diffusion-tensor anisotropy index, defined as 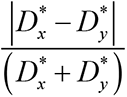 from the homogenization closure-problem solution (bounded 0-1; a normalized companion measure to the raw ratio D_x/D_y reported in Figure 2h). (e) Direction-resolved effective diffusion coefficient D*_x. (f) Direction-resolved effective diffusion coefficient D*_y. (g) Coefficient of variation (CV) of the local ECS channel-width distribution. (h) 10th percentile (P10) of the local ECS channel-width distribution. (i) 90th percentile (P90) of the local ECS channel-width distribution. (j) Time to 10% breakthrough (T10) from the same constant-source breakthrough simulations as Figure 2j-k.

**Supplementary Figure 3.**
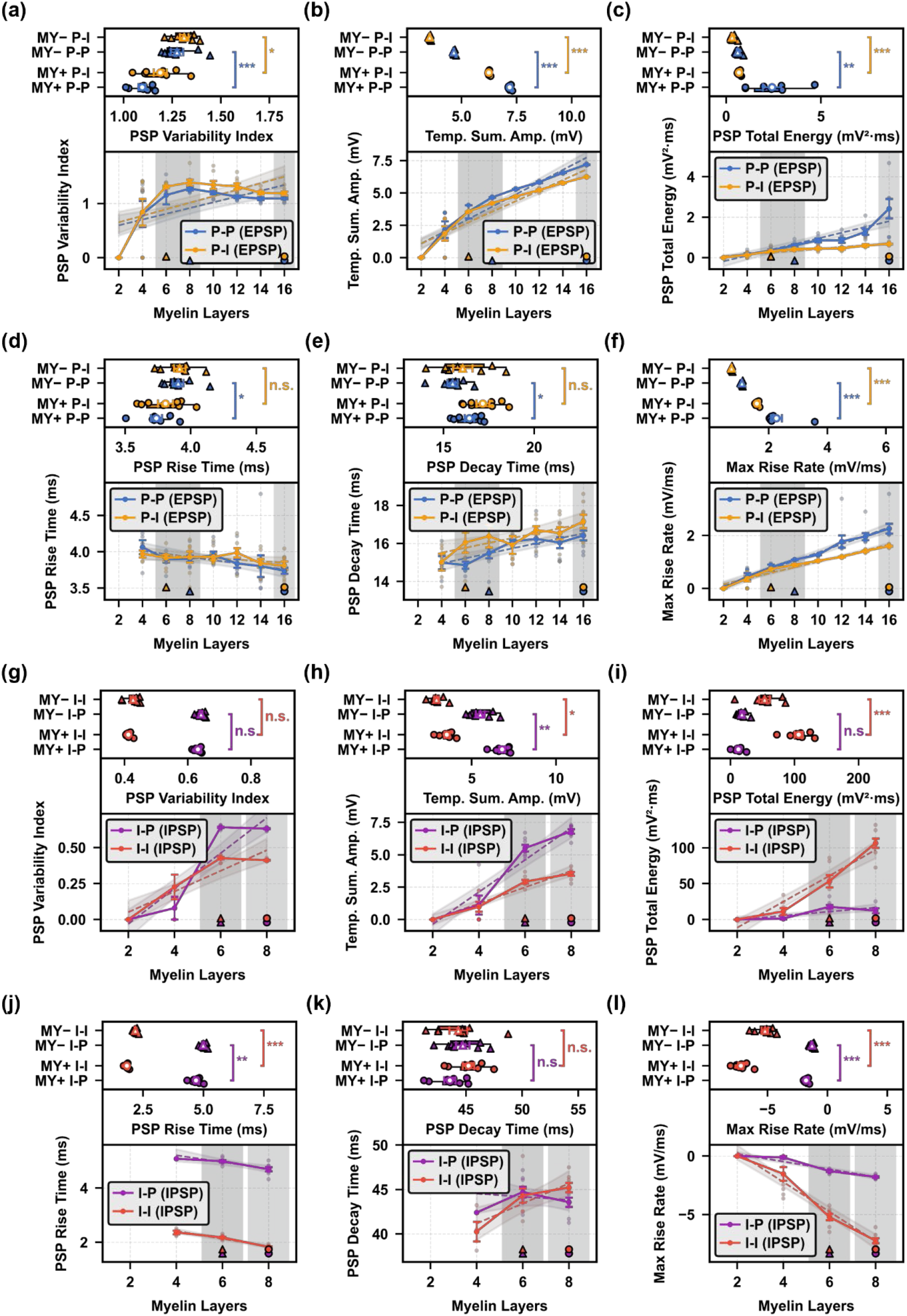
PSP waveform-shape, temporal-summation and energetic properties across the myelin-thickness gradient, for the same successful-transmission trials underlying Figure 3h,i. All panels compare layer 6 (MY−, lowest layer with 100% transmission success across repeats) against the maximum layer tested (MY+: layer 16 for P–P/P–I, layer 8 for I–P/I–I), Welch’s *t*-test or Mann–Whitney *U* (\*\*\**p* < 0.001, \*\**p* < 0.01, \**p* < 0.05, n.s. not significant), with the full dose–response trend across all layers shown below each comparison (mean ± s.e.m.). (a)–(f) EPSP-forming connections (P–P, blue; P–I, orange): (a) PSP variability index. (b) Temporal-summation amplitude (paired-pulse peak amplitude). (c) Total PSP energy. (d) PSP rise time. (e) PSP decay time. (f) Maximum PSP rise rate (peak d*V*/d*t*). (g)–(l) IPSP-forming connections (I–P, purple; I–I, red), same six metrics as (a)–(f).

**Supplementary Figure 4.**
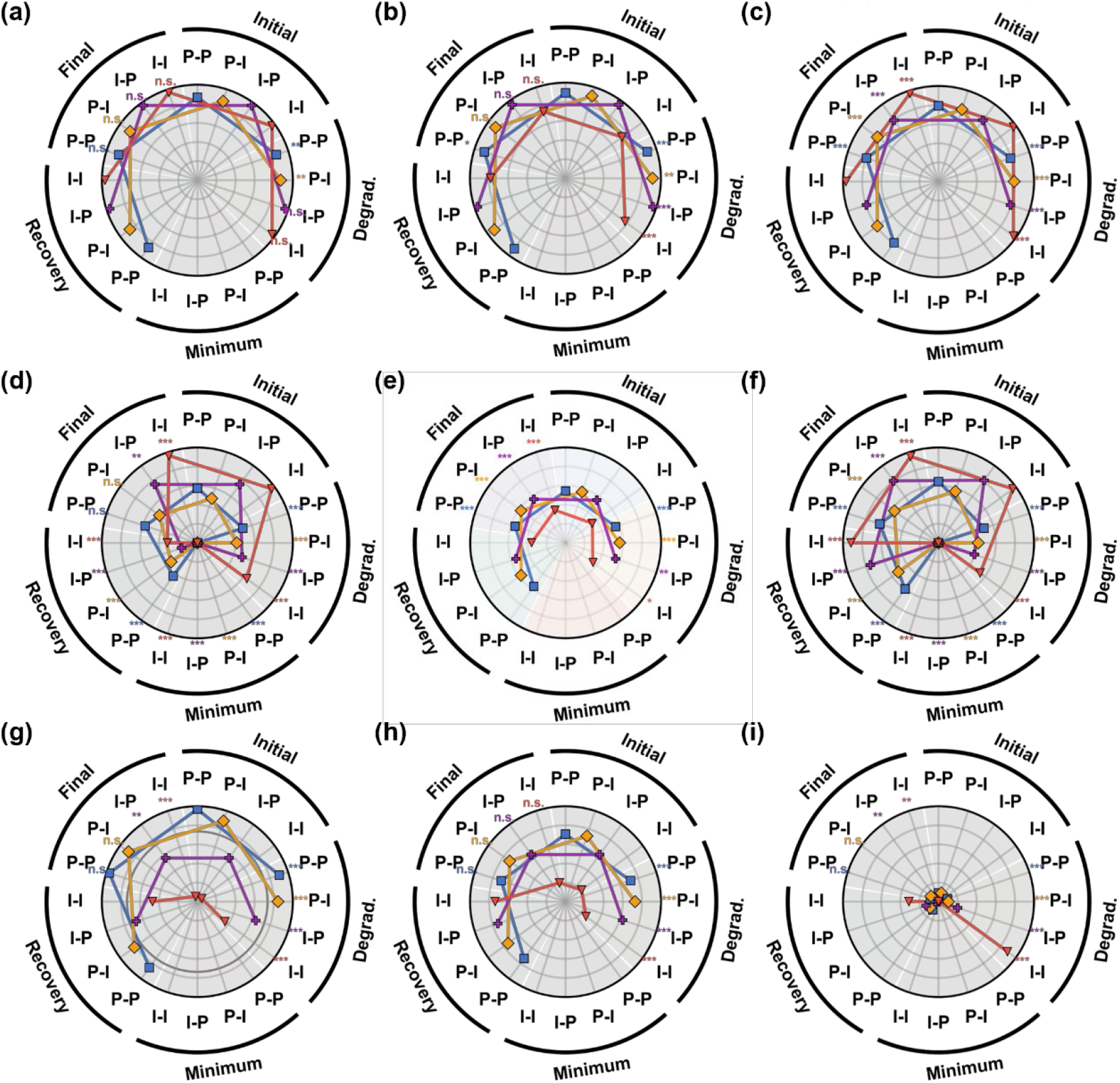
PSP kinetics, waveform-shape and temporal-summation metrics across the continuous five-phase demyelination–remyelination cycle, for all four connection types. Grouped radar (spider) plots: each of the five phases (Initial, Degradation, Minimum, Recovery, Final) is represented as one radial axis; each connection type (P–P, blue square; P– I, orange diamond; I–P, purple cross; I–I, red triangle) is drawn as a single open polyline connecting its mean value across the five phase-axes (no fill; 8 repeats per point). Missing vertices at Minimum reflect the fact that no PSP is generated at that phase (0% transmission success; see Fig. 4f), not a plotting artefact. Significance markers next to each phase label give the paired comparison against the Initial-phase baseline for that connection type (paired t-test or Wilcoxon signed-rank, matched by repeat; \*\*\**p* < 0.001, \*\**p* < 0.01, \**p* < 0.05, n.s. not significant); Minimum is excluded from statistical comparison for the reason above. All axes use a linear radial scale; panels are otherwise independent (each metric has its own radial range). (a) PSP time-to-peak. Effectively unchanged across the four transmitting phases for all connection types (e.g. P–P: 8.36 → 8.41 → 8.48 → 8.36 ms; Initial vs Final n.s. for all connection types), confirming that the intrinsic rise kinetics of the synaptic conductance are not confounded with the axon’s myelin state (positive control; see Discussion). (b) PSP rise time. Likewise near-invariant (P–P/P–I: ∼7.2 ms; I–P/I–I: ∼11.1–11.4 ms); Initial-vs-Final comparisons are non-significant for 3 of 4 connection types. (c) PSP 90%-bandwidth. Stable within each connection type (P–P/P–I/I–P: ∼0.625; I–I: ∼1.25) but with small, statistically significant Initial-vs-Degradation and Initial-vs-Final shifts (*p* < 0.001 for all four connection types), indicating a subtle, fully reversible spectral narrowing that accompanies — without reflecting the dominant driver of — the amplitude/energy collapse in Fig. 4d,e. (d) PSP area-under-curve. Tracks the amplitude/energy collapse-and-recovery pattern of Fig. 4d,e (Initial-vs-Degradation and Recovery-vs-Final paired comparisons, both *p* < 0.001 for all four connection types; one-sample tests confirm non-zero AUC at all phases except Minimum for P–P, n.s./marginal for the other three). (e) Paired-pulse summation (peak amplitude of the temporally summated response). The same underlying peak-amplitude metric as Fig. 4h, re-plotted here in the grouped-radar layout for direct at-a-glance comparison across all four connection types (Initial-vs-Degradation and Recovery-vs-Final, both p < 0.001 for all connection types except one *p* = 0.022 case). (f) Paired-pulse total AUC. Parallels the ∼21-fold vs ∼3-fold asymmetry seen for paired total energy (Fig. 4i) between interneuron-and pyramidal-sourced connections (Initial-vs-Degradation, *p* < 0.001 for all four connection types). (g) PSP initial slope (rising-phase mV/ms). Declines from Initial to Degradation in all four connection types (*p* < 0.001) with sign set by EPSP (positive, P–P/P–I) versus IPSP (negative, I–P/I–I) polarity; largely recovers by Final (n.s. for P–P/P–I, but not for I–P/I–I, *p* ≤ 0.002). (h) PSP symmetry index. Significant Initial-vs-Degradation shift for all four connection types (*p* < 0.001); fully reversible by Final for P–P/P–I (identical values, no variability) and I–I (n.s.), but not for I–P (*p* = 0.195, n.s., trending). (i) PSP kurtosis. The most strongly affected waveform-shape metric: significant Initial-vs-Degradation change for all four connection types (*p* < 0.001), including a qualitative sign change in the inhibitory connections (I–P: −1.24 → 4.80; I–I: −1.21 → 24.5), indicating a pronounced platykurtic-to-leptokurtic transition of the IPSP waveform under demyelination that is not seen for the excitatory connections.

**Supplementary Figure 5.**
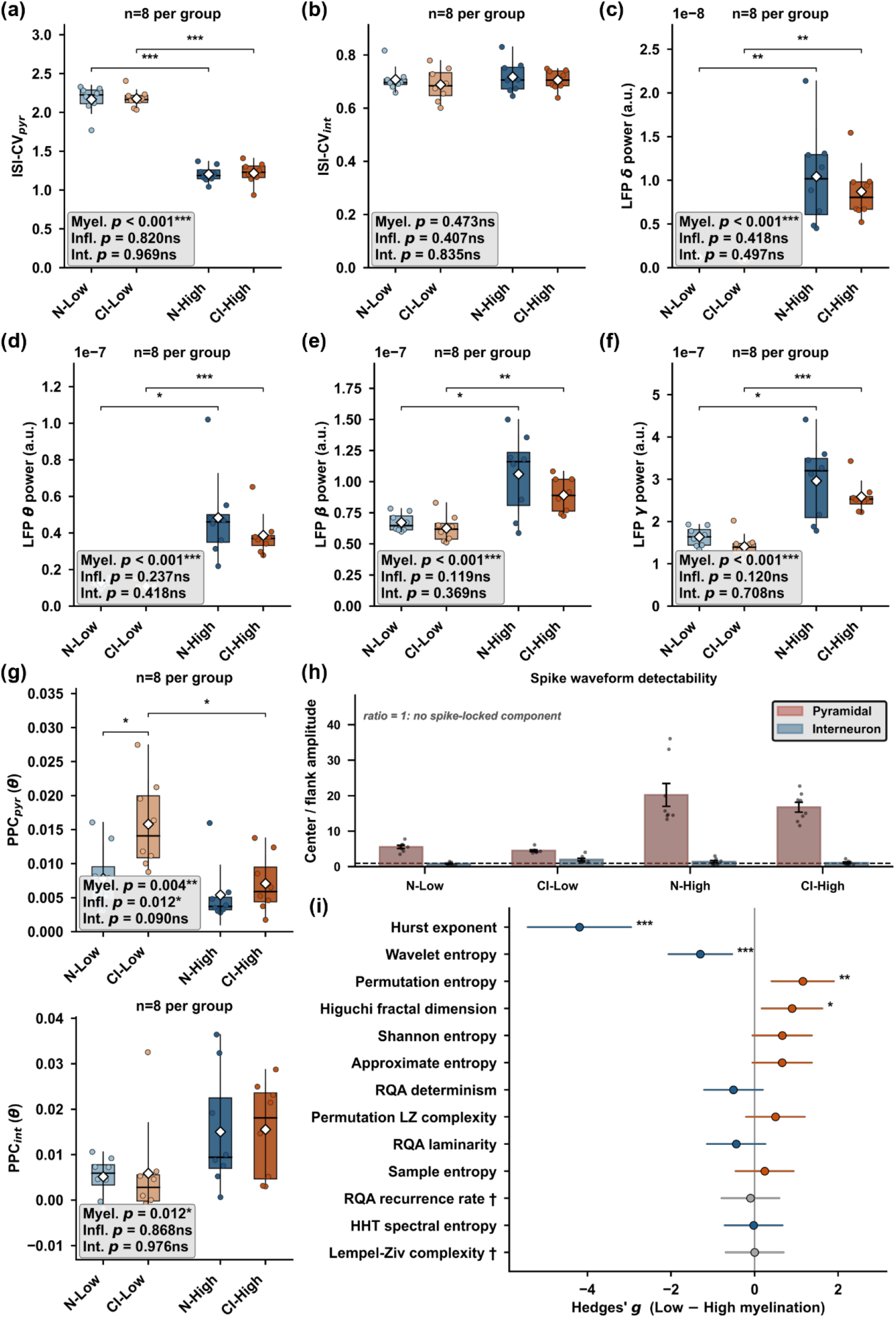
Discharge regularity, band power, spike–field phase consistency and effect sizes across the 2 × 2 design. Conditions, colours, *n* and box conventions as in Figure 5. Each boxed inset gives the three terms of the 2 × 2 factorial ANOVA: Myel., myelination main effect; Infl., inflammation main effect; Int., interaction. Brackets show the four pre-specified contrasts after Holm–Bonferroni correction within the measure. (a, b) Coefficient of variation of the interspike interval for pyramidal cells and interneurons. Pyramidal discharge is far more irregular at MY− (CV ≈ 2.2, burst-dominated) than at MY+ (CV ≈ 1.2, near-Poisson), whereas interneuron irregularity is unaffected by either factor. (c–f) Band power of the local field potential in the delta (0.5–4 Hz), theta (4–12 Hz), beta (12–30 Hz) and gamma (30–80 Hz) bands, obtained as the integral of the multitaper spectral density over each band and reported in arbitrary units. All four bands are elevated at MY+, by 8.7-fold in delta down to 1.6-fold in beta; none differs by inflammatory state. (g) Pairwise phase consistency between spikes and theta-band field phase, for pyramidal cells (upper) and interneurons (lower). Pairwise phase consistency is used in preference to the phase-locking value because its expectation does not depend on spike count. The upper panel is the only measure in this study in which the inflammatory manipulation produced a significant effect: consistency roughly doubles when inflammation is added to the demyelinated network (inflammation main effect *p* = 0.012, and the N-Low against CI-Low contrast survives Holm correction at *p* = 0.047), with no equivalent change at MY+ and no significant interaction (*p* = 0.090). The same panel also carries a myelination main effect (p = 0.004); the second bracket marks the corresponding fall in consistency from the inflamed demyelinated networks to their myelinated counterparts. The lower panel shows a myelination main effect (p = 0.012, inflammation p = 0.868) for which neither pre-specified contrast survived correction. (h) Detectability of spike waveforms in the simulated field, quantified as the ratio of the centre amplitude of the mean waveform to its flank amplitude; a ratio of 1 (dashed line) means no spike-locked component is present. Bars are the condition mean, error bars SEM, dots individual networks. Pyramidal spikes are three to four times better resolved at MY+ than at MY−, and interneuron spikes are not resolvable in any condition, which is why all interneuron measures in this study are derived from spike times rather than from waveforms. (i) Effect of myelination on all thirteen nonlinear descriptors of the field potential, as Hedges’ *g* for MY− minus MY+ with 95 % confidence interval. Blue, lower at MY−; orange, higher at MY−. Asterisks give the myelination main effect of the 2 × 2 ANOVA (\**p* < 0.05, \*\**p* < 0.01, \*\*\**p* < 0.001); the four descriptors that reach significance also survive Benjamini–Hochberg correction across the thirteen (*q* = 9.1 × 10⁻¹², 4.9 × 10⁻³, 0.013 and 0.049). Daggers mark the two descriptors that are degenerate on these data, the recurrence rate being fixed at 5 % by construction of the recurrence threshold and the median-binarised Lempel–Ziv complexity taking only eight distinct values across the thirty-two runs; both are shown with their zero effect rather than omitted.

**Supplementary Figure 6.**
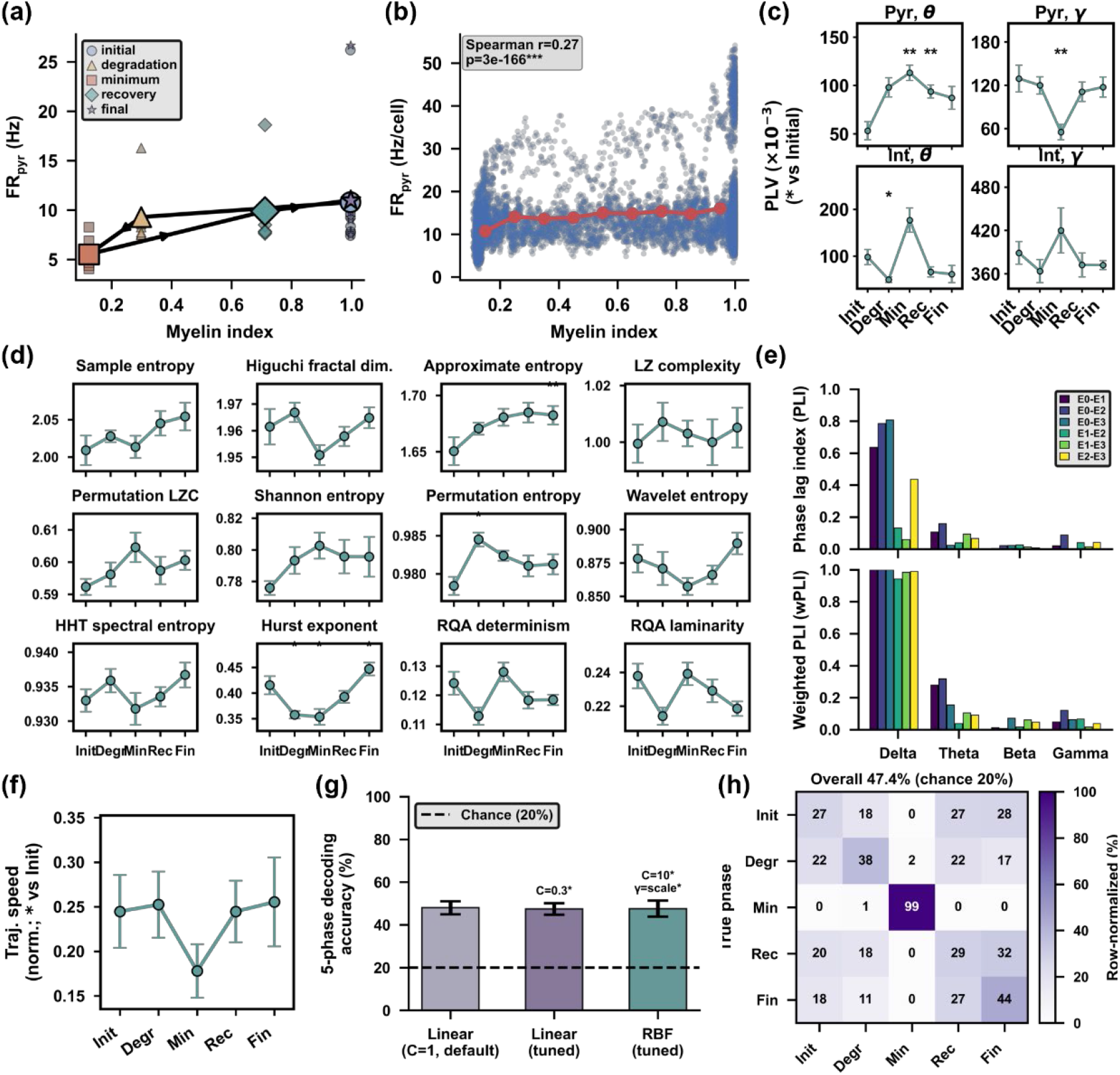
Rate-myelin relationship, phase locking, the full nonlinear descriptor set, inter-electrode connectivity and decoding controls for the demyelination and remyelination trajectory. Phases, n and pairing as in Figure 6. (a) Phase-mean pyramidal firing rate plotted against phase-mean myelin index, with the five phases as distinct markers and lines joining them in temporal order. The descending and ascending limbs do not coincide: Degradation and Recovery differ 2.4-fold in mean myelin index but only 6 % in rate. Because the two transition phases average over different distributions of instantaneous myelin index, this panel is descriptive; the direct test of path dependence is Initial against Final, which is null. (b) Instantaneous pyramidal firing rate per cell in 20 ms bins against the concurrent myelin index, pooled over all networks (blue points, n = 9968 bins), with binned means in red. Rates in this panel are taken from the raw detected spike train shipped with each simulation and therefore sit about 1.5-fold above the quality-controlled rates in Fig. 6c; only the rank correlation is reported. (c) Phase-locking value between spikes and field phase, in the theta and gamma bands for pyramidal cells and interneurons, on the same spike-count-matched trains as Fig. 6g and 6i. The phase-locking value reproduces the pyramidal theta-up and gamma-down dissociation independently; it diverges from pairwise phase consistency only for interneuron theta during Degradation, where its residual spike-count sensitivity is the likely explanation. (d) All twelve non-degenerate nonlinear descriptors of the local field potential by phase, mean ± SEM across networks. The recurrence rate is omitted because it is fixed at 5 % by construction of the recurrence threshold. Only the Hurst exponent, approximate entropy and permutation entropy reach significance after Holm–Bonferroni correction within descriptor and Benjamini-Hochberg correction across descriptors; these three are summarised as effect sizes in Fig. 6f. (e) Phase lag index (top) and weighted phase lag index (bottom) between all six pairs of the four tetrode contacts of one network, by frequency band, over the full 25 s recording. Both indices are insensitive to zero-lag coupling and therefore to volume conduction. Only delta shows a consistent non-zero-lag phase relationship (weighted index 0.942 to 1.000 across pairs); above delta both indices collapse (weighted index 0.037 to 0.317 in theta and 0.016 to 0.121 in gamma), indicating that inter-contact coherence at higher frequencies reflects a shared field source rather than coupling between distinct generators. The four contacts span tens of micrometres within one tetrode, so this panel constrains the interpretation of the local field and not long-range connectivity. (f) Normalised speed of the population-state trajectory by phase, compared against Initial. The dip at Minimum does not reach corrected significance (p = 0.129). (g) Five-phase decoding accuracy for three classifiers under leave-one-network-out cross-validation with nested hyperparameter tuning: a linear kernel at the default regularisation, a tuned linear kernel and a tuned radial-basis kernel. Annotations give the modal selected hyperparameters across the eight outer folds. Bars are the mean over folds, error bars the SD, and the dashed line the 20 % chance level. All three agree, and a control using only the three leading principal components decodes at 21.1 ± 10.4 %, that is, at chance. (h) Row-normalised confusion matrix of the tuned linear classifier, reproduced from Fig. 6m for reference alongside (g).

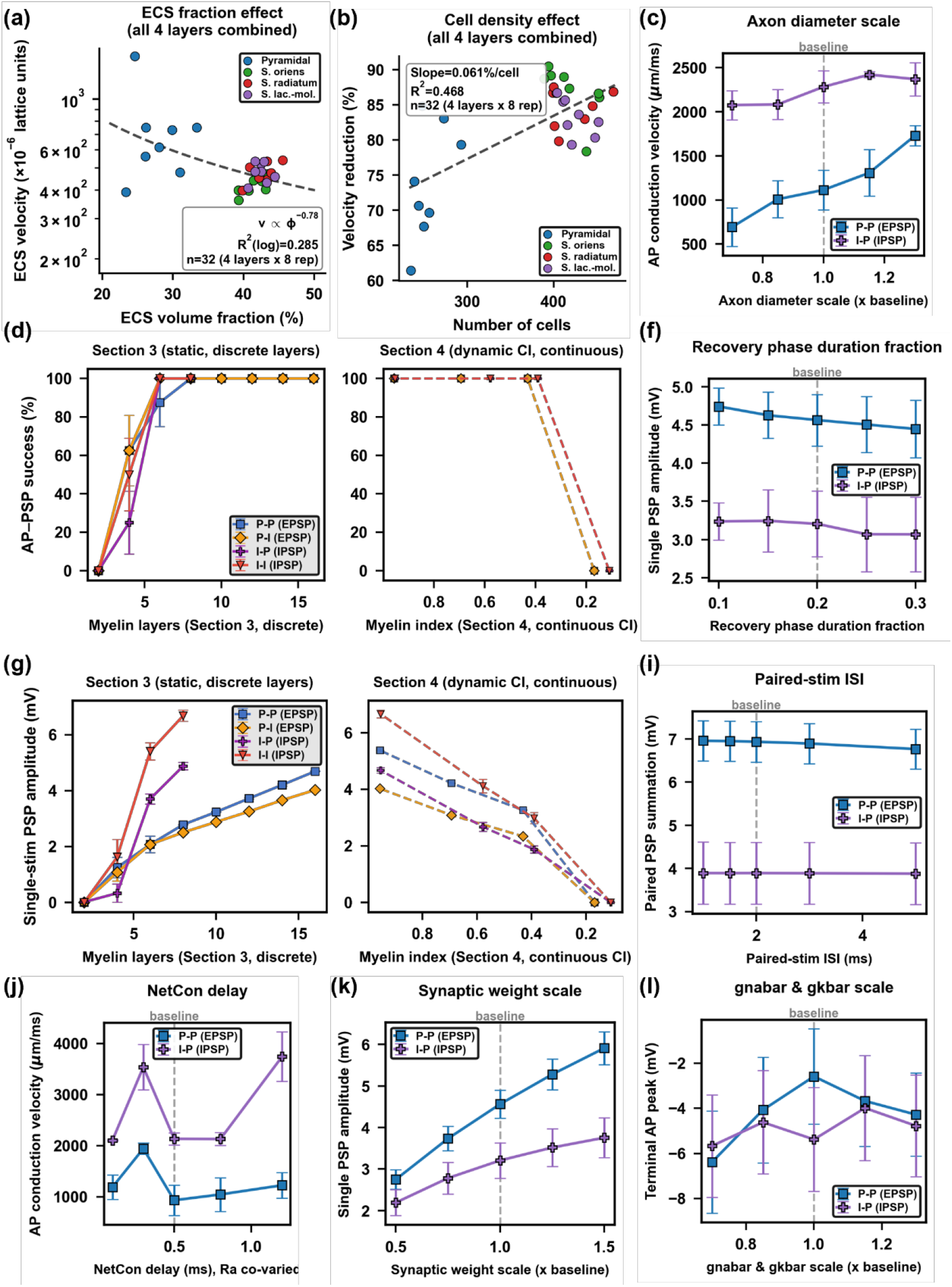

**Supplementary Figure 8.**
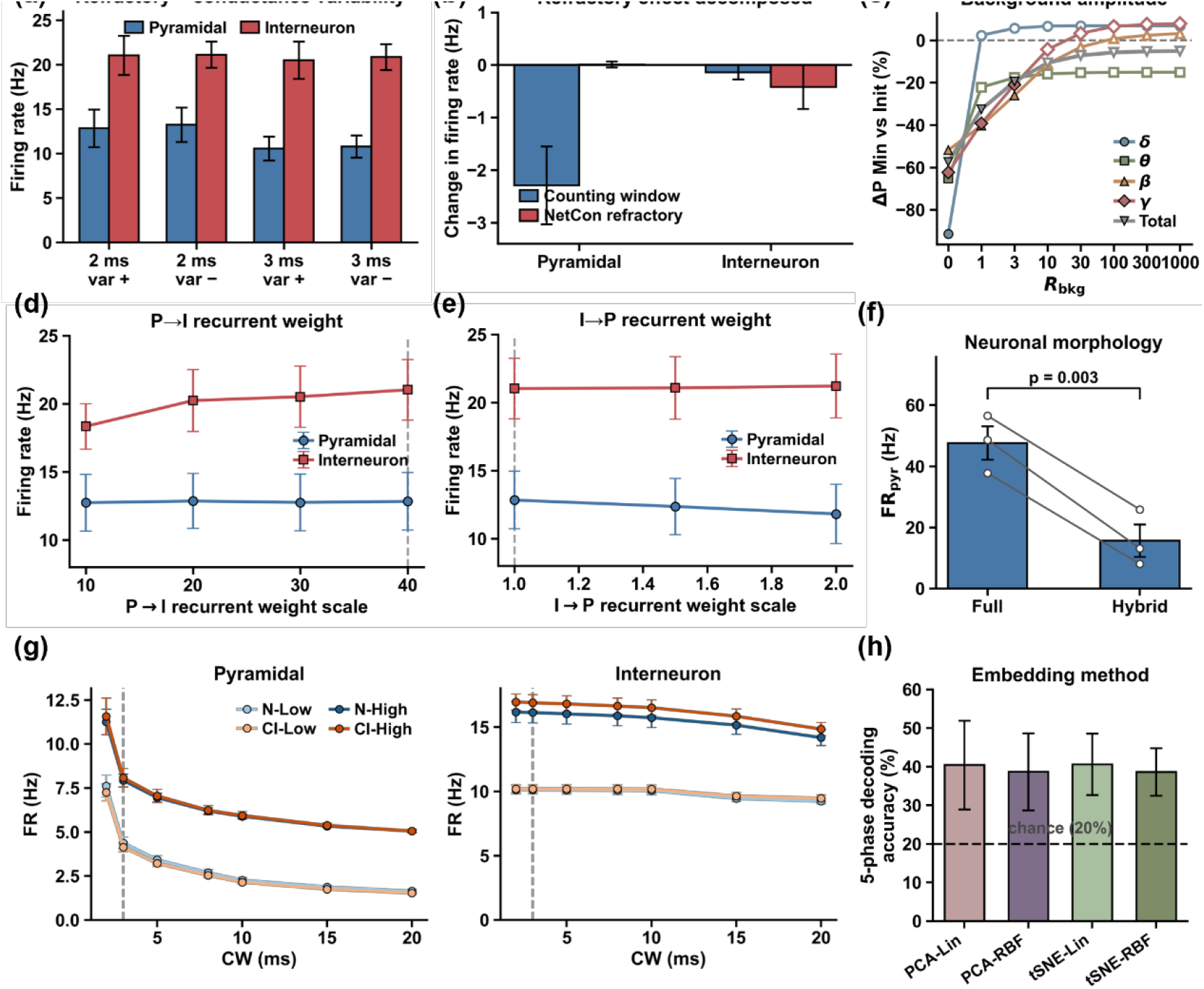
Sensitivity and robustness analyses for the network simulations of Sections 5 and 6. (a, b, d, e) Parameter sensitivity of the Section 5 network. Each condition was run with three independent network seeds, and within a seed every condition shares the same cell census and wiring, so conditions are compared as paired same-network contrasts; bars and points give the mean with the SEM across the three networks, and p values are paired t-tests against the baseline condition. With three networks the design resolves only large effects, so the informative quantities here are the direction of the change and its consistency across seeds rather than significance. (a) Mean firing rate under the two refractory conventions (2 ms and 3 ms) crossed with the presence (var +) or absence (var −) of cell-to-cell variability in the Na⁺ and K⁺ conductances. Removing conductance variability changed pyramidal rate by +0.40 Hz (p = 0.46). Moving to the 3 ms convention lowered pyramidal rate by 2.28 Hz with variability present and by 2.06 Hz with it absent, in the same direction in all three networks in both cases (p = 0.098 and 0.163). Interneuron rate was insensitive to both factors (all |Δ| ≤ 0.56 Hz). (b) The 2 ms to 3 ms difference of (a) decomposed into the two mechanisms it conflates, by re-detecting spikes in the identical 2 ms simulation with a 3 ms window (counting-window component) and taking the remainder as the effect of the 3 ms refractory period in synaptic transmission. For pyramidal cells the entire change is the counting window (−2.29 ± 0.74 Hz) and the transmission component is nil (+0.01 ± 0.06 Hz); for interneurons both components are small (−0.14 ± 0.14 and −0.42 ± 0.42 Hz). The apparent sensitivity of pyramidal rate to the refractory setting is therefore a counting convention applied to genuine high-frequency discharge, not a change in how spikes propagate. (c) Robustness of the Section 6 band-power result to additive background activity. Synthetic background was added to the field potential at a background-to-signal power ratio R (R_bkg on the axis) from 0 to 1000 and the change in band power at Minimum against Initial recomputed. Filled symbols mark comparisons that remain significant across the eight networks (Wilcoxon signed-rank, *p* < 0.05), open symbols those that do not. The measured reductions of roughly 60 % are recovered in full only when the background is negligible; delta and theta lose significance once R reaches 1, beta, gamma and total power hold to R = 3, and no band remains significant at R = 10. This is a deliberately adverse bound rather than an estimate of in vivo conditions, since the added background is broadband and uncorrelated with the signal. (d) Recurrent pyramidal-to-interneuron weight scaled from 10 to 40, with 40 the baseline. Interneuron rate fell as the drive was weakened, most clearly at the lowest setting (−2.69 Hz, p = 0.054, same direction in all three networks), and pyramidal rate was unchanged (all |Δ| ≤ 0.10 Hz). (e) Recurrent inhibitory-to-pyramidal weight scaled 1.5-fold and 2-fold above baseline. Pyramidal rate fell monotonically and in the same direction in all three networks (−0.47 and −1.02 Hz; p = 0.17 and 0.26) and interneuron rate was unchanged, which is the expected sign for stronger feedback inhibition. (f) Mean pyramidal firing rate (FR_pyr) with the full reconstructed morphology imposed on every neuron (Full) against the spatially graded configuration used in production (Hybrid), in which only neurons inside the core volume around the electrodes carry their full morphology. Bars give the mean with the SEM across three network seeds, and the connecting lines join the two configurations within a seed, the cell census being held fixed inside each seed so that morphology is the only variable. The full morphology yields rates about three-fold higher (47.7 against 15.7 Hz, paired t-test p = 0.003), in the same direction in all three seeds. The magnitude should be read as a bracket rather than as a calibration of the production offset, because at twenty cells the number of pyramidal cells that happen to fall inside the core volume, and therefore keep their full morphology in the graded configuration, varies strongly between seeds: 0 of 8, 2 of 7 and 5 of 11. Rate tracks that proportion monotonically (8.1, 13.1 and 25.9 Hz respectively), which identifies the amount of morphological detail carried by the population as the quantity that sets the absolute rate scale. What the panel establishes is the direction and the order of magnitude, and that both are stable across seeds. It is an offset of the baseline rate rather than a change in how the network responds to demyelination: every comparison in this work is made either between groups simulated under one and the same morphological configuration or between phases within a single network, so the offset cancels in all reported contrasts. Only the pyramidal comparison is shown, because this reduced configuration contains between zero and two interneurons per run and so does not support a population estimate for that class. (g) Dependence of the mean firing rates of Figure 5f on the width of the window used to collapse repetitive discharge into a single counted event (CW), from 2 ms to 20 ms; the dashed line marks the 3 ms convention used throughout. The four groups carry the labels of Figure 5, N-Low and CI-Low being the low-myelination groups without and with inflammation and N-High and CI-High their high-myelination counterparts. Absolute rates fall as the window widens, as they must, but the two myelination states remain separated at every window tested, with the two low-myelination groups below the two high-myelination groups throughout and no crossing anywhere in the range, so the myelination effect is not an artefact of the particular value chosen. (h) Robustness of the five-phase decoding of Section 6 to the choice of embedding, comparing a three-dimensional principal component analysis with a two-dimensional t-distributed stochastic neighbour embedding, each with a linear and a radial-basis kernel (PCA-Lin, PCA-RBF, tSNE-Lin, tSNE-RBF), all under the same leave-one-network-out cross-validation. Accuracy is 38.7 % to 40.8 % in all four combinations against a chance level of 20 %. Three accuracies should be kept distinct. The main analysis of Figure 6m uses the seven population descriptors directly, with no embedding at all, and reaches 47.5 %. This panel embeds the Procrustes-aligned trajectories in three or two dimensions first, which costs about 7 percentage points but is insensitive to which embedding is used. Omitting the alignment, so that each network is described in its own arbitrary principal-component basis, collapses accuracy to chance (21.1 %, reported in the text). The panel therefore shows that the low-dimensional view of Figure 6l is not what carries the phase information, and that the choice between principal component analysis and t-distributed stochastic neighbour embedding for that view is immaterial.

